# Enhancing the detection of HTT1a with neoepitope antibodies in mouse models of Huntington’s disease

**DOI:** 10.64898/2026.03.01.708805

**Authors:** Georgina F. Osborne, Edward J. Smith, Kirupa Sathasivam, Zixin Kang, Iulia M. Nita, Maria Cañibano-Pico, Jemima Phillips, Gillian P. Bates, Christian Landles

**Author notes:** Correspondence to: Christian Landles, Huntington’s Disease Centre and Department of Neurodegenerative Disease, Queen Square Institute of Neurology, University College London, Queen Square, London, WC1N 3BG, United Kingdom.

## Abstract

Huntington’s disease is an inherited neurodegenerative disorder caused by a CAG repeat expansion in exon 1 of the Huntingtin (*HTT*) gene, encoding an expanded polyglutamine tract in the huntingtin (HTT) protein. The pathogenic CAG repeat of *HTT* is unstable and undergoes progressive somatic expansion in specific brain cells and peripheral tissues throughout life. Genes involved in DNA mismatch repair pathways, which promote repeat expansion, have been identified as genetic modifiers of the disease. Consequently, the rate of CAG repeat expansion is a key determinant driving the age of onset and disease progression. As the CAG repeat expands, alternative processing of *HTT* pre-mRNA increasingly favours production of the *HTT1a* transcript, which encodes the highly pathogenic and aggregation-prone HTT1a protein. This process provides a mechanistic link between CAG repeat expansion and disease pathogenesis, as increased HTT1a production accelerates HTT aggregation and neuronal dysfunction.

HTT1a has previously been detected in Huntington’s disease mouse models by using immunoprecipitation coupled with western blotting, homogeneous time-resolved fluorescence (HTRF) and Meso Scale Discovery (MSD) bioassays, and immunohistochemistry. These approaches were developed using MW8, a neoepitope antibody that specifically recognizes the C-terminus of HTT1a. MW8 is a relatively weak antibody with limited detection sensitivity. To generate more robust HTT1a-specific reagents, two novel recombinant antibodies, 1B12 and 11G2, have been developed for evaluation. Using an allelic series of knock-in (*Hdh*Q20, *Hdh*Q50, *Hdh*Q80, *Hdh*Q111, CAG140 and zQ175) mice, alongside transgenic YAC128 and N171-82Q models, we extensively evaluated and compared the performance of MW8, 1B12 and 11G2. We demonstrate that 1B12 and 11G2 function as HTT1a-specific neoepitope antibodies by immunoprecipitation with western blotting, and by immunohistochemistry. To enhance HTT1a detection using HTRF and MSD technology platforms, we further evaluated the performance of 1B12 and 11G2 in HTT bioassays using cortical lysates from zQ175 and YAC128 mice. In zQ175 mice, enhanced detection of aggregated HTT1a by HTRF and MSD revealed that HTT fragments longer than HTT1a can be incorporated into HTT1a-containing aggregates. The most sensitive assays were subsequently applied across the allelic series of knock-in mice to assess the effect of polyglutamine length on bioassay performance. For optimal sensitivity, we recommend the preferential use of 1B12 for HTRF assays and 11G2 for MSD assays. Collectively, these findings establish 1B12 and 11G2 as robust antibodies to reliably detect and track HTT1a pathology *in vivo* and promotes the replacement of previously used MW8-based experimental approaches.

**GRAPHICAL ABSTRACT:** 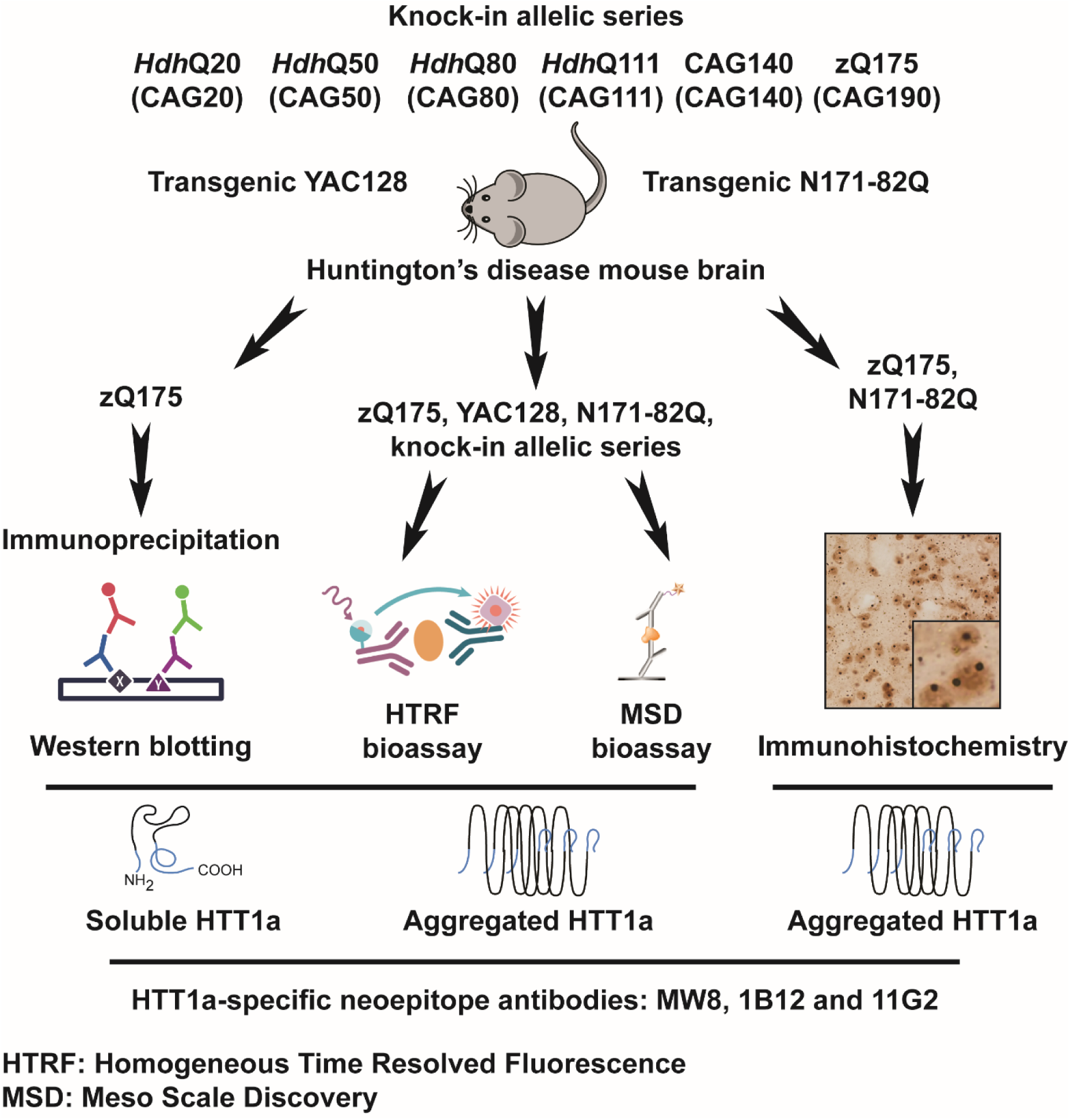

Osborne *et al*. used Huntington’s disease mouse models to evaluate and compare the performance of HTT1a-specific neoepitope antibodies by using immunoprecipitation with western blotting, bioassays, and immunohistochemistry. In contrast to MW8, they establish that 1B12 and 11G2 are robust antibodies to reliably detect and track HTT1a pathology *in vivo*.

## INTRODUCTION

Huntington’s disease is an autosomal dominant neurodegenerative disorder characterized by motor dysfunction, cognitive decline, and psychiatric disturbances.^1^ The disease is caused by the expansion of a glutamine-coding CAG trinucleotide repeat in exon 1 of the huntingtin (*HTT*) gene, which leads to an abnormally long polyglutamine (polyQ) tract in the huntingtin (HTT) protein.^2^ *HTT* alleles with 40 or more CAGs are fully penetrant, while those with 35 CAGs or less do not cause disease.^3^ Individuals carrying alleles with 35–39 CAGs have an increased risk of developing Huntington’s disease within a normal lifespan,^4^ whilst expansions of 60 CAGs or more repeats are typically associated with juvenile-onset forms of the disorder.^5^ Genetic evidence suggests that expression of HTT containing an expanded polyQ tract confers a toxic gain-of-function leading to selective neuronal dysfunction and cell death.^2,6^

The CAG repeat within the *HTT* gene is somatically unstable and undergoes age-dependent expansions in specific brain regions.^7–10^ Multiple DNA mismatch repair pathway genes have been identified as genetic modifiers of Huntington’s disease that can influence the rate of somatic repeat expansion, and as a result, CAG instability is widely considered a key driver of both age-of-onset and the rate of disease progression.^11–14^ Amongst the cell types assessed, medium spiny neurons are the most susceptible to somatic instability and show the earliest and most pronounced CAG repeat expansion,^10,15,16^ although the mechanisms underlying this selective vulnerability remain unclear. Expanded CAG tracts can promote the alternative processing of *HTT* pre-mRNA to generate the *HTT1a* transcript, which encodes the highly pathogenic and aggregation-prone HTT1a protein.^17–19^ Notably, HTT1a levels increase as CAG repeat length expands,^17,20^ providing a mechanistic link through which somatic CAG repeat expansion may exert its pathogenic effects, as supported by a recent study demonstrating that prevention of *HTT1a* production ameliorated disease phenotypes.^21^

The accurate quantification of HTT protein isoforms across disease models and clinical samples is essential for mechanistic studies and for evaluating preclinical and clinical interventions, and is critical for interpreting therapeutic strategies designed to directly reduce *HTT* expression.^6^ Numerous immunoassays have been developed to measure soluble and aggregated HTT species using pairs of anti-HTT antibodies that recognize distinct epitopes. In cellular lysates and tissue extracts from Huntington’s disease models, assays have been successfully established on multiple analytical platforms, including homogeneous time-resolved fluorescence (HTRF),^22–24^ Meso Scale Discovery (MSD),^24–26^ and amplified luminescent proximity homogeneous assay (AlphaLISA).^24,27^ However, because HTT concentrations in cerebrospinal fluid are substantially lower, more sensitive single-molecule counting assays have been required to accurately detect and quantify HTT.^28,29^ To enable the measurement of HTT1a using the HTRF and MSD platforms, we previously developed and published two HTT1a-specific assays: (i) a soluble HTT1a assay using the N-terminal HTT 2B7 antibody, and (ii) an aggregated HTT1a assay using the human-specific proline-rich domain HTT 4C9 antibody, each paired with the C-terminal HTT1a MW8 antibody.^20,24,30^ Recently, to further improve the sensitivity of HTT1a detection, two recombinant antibodies (namely, 1B12 and 11G2) were generated, which like MW8,^31^ recognize the C-terminus of HTT1a.^32^

In this study, we used immunoprecipitation and western blotting of cortical lysates from zQ175 mice, together with immunohistochemistry of zQ175 and N171-82Q mouse tissue sections, to demonstrate that 1B12 and 11G2, like MW8, function as neoepitope-specific antibodies for the detection of soluble and aggregated HTT1a. Using cortical lysates from zQ175 and YAC128 mice on the HTRF and MSD platforms, we showed that assays incorporating 1B12 or 11G2 detected soluble and aggregated HTT1a with greater sensitivity than our previously published MW8-based assays.^24^ Similarly, immunohistochemical analysis of zQ175 tissue sections demonstrated that 1B12 and 11G2 detected a diffuse nuclear aggregate signal, in addition to nuclear and cytoplasmic inclusions, with greater sensitivity than our MW8-based approach.^30^ Notably, the improved detection of aggregated HTT1a by HTRF and MSD assays revealed that HTT fragments longer than HTT1a can be incorporated into HTT1a-containing aggregates in zQ175 mice. Finally, using the most sensitive HTRF and MSD assays on hippocampal lysates from an allelic series of knock-in mice (*Hdh*Q20, *Hdh*Q50, *Hdh*Q80, *Hdh*Q111, CAG140 and zQ175 mice), we assessed the effect of polyglutamine length on HTT1a bioassay performance. Collectively, these findings establish 1B12 and 11G2 as robust antibody reagents to reliably detect and track HTT1a pathology *in vivo*.

## MATERIALS AND METHODS

### Ethical statement

All procedures complied with the Animals (Scientific Procedures) Act, 1986 and were approved by the University College London (UCL) Ethical Review Process Committee.

### Mouse breeding and maintenance

Heterozygous, homozygous and wild-type littermates from the *Hdh*Q20^33^ and zQ175^34,35^ knock-in lines on a C57BL/6J background were imported from CHDI Foundation colonies at the Jackson Laboratory (Bar Harbor, Maine) and housed at UCL until tissue collection at 2, 6, or 12 months of age. Transgenic YAC128 or transgenic N171-82Q mouse colonies were bred at UCL by pairing transgenic males with C57BL/6J females (Charles River), and tissues were collected at either 2, 6, or 12 (YAC128) or 1.5 and 3 (N171-82Q) months of age. Mouse husbandry and health monitoring were performed as previously described.^36^ Animals were group-housed by sex, with mixed genotypes per individually ventilated cage, containing Aspen Chips 4 bedding and environmental enrichment that consisted of a play tunnel and chew sticks (Datesand). Mice had *ad libitum* access to water and food (Teklad global 18% protein rodent diet, Inotiv). The environment was maintained at 21°C ± 1°C under a 12 h light/dark cycle. Animals were housed under barrier-maintained conditions with quarterly non-sacrificial screening (Federation of European Laboratory Animal Science Associations) to confirm pathogen-free status. Following euthanasia, tissues were rapidly dissected, snap-frozen in liquid nitrogen, and stored at-70°C.

### DNA extractions, genotyping, and CAG repeat sizing

Genomic DNA was extracted from ear biopsies.^37^ The expanded *HTT* allele in YAC128 mice contains a stable polyQ repeat of 125 residues, encoded by (CAG)_23_(CAA)_3_CAGCAA(CAG)_80_(CAA)_3_CAGCAA(CAG)_10_CAACAG,^38^ and mice were genotyped as previously described.^24^ Transgenic N171-82Q mice express a *HTT* transgene that contains a stable polyQ repeat of 82 residues encoded by (CAG)₈₀CAACAG, and mice were genotyped as previously described.^39^ For imported mice, tail biopsies were collected upon sacrifice for genotype confirmation. *Hdh*Q20 mice were genotyped as previously described and contain a stable repeat of 18 CAGs.^24^ zQ175 mice were genotyped as previously described and repeat sizing was performed by the CHDI Foundation prior to shipment to UCL.^24^ The mean CAG repeat length for all zQ175 mice used in this study was 192.9 ± 4.8(SD).

### Antibodies

The details of all antibodies used in this study are provided (Supplementary Table 1), and the location of the HTT epitopes recognized by these antibodies on the HTT protein is illustrated (Fig. 1A).

**Figure 1.**
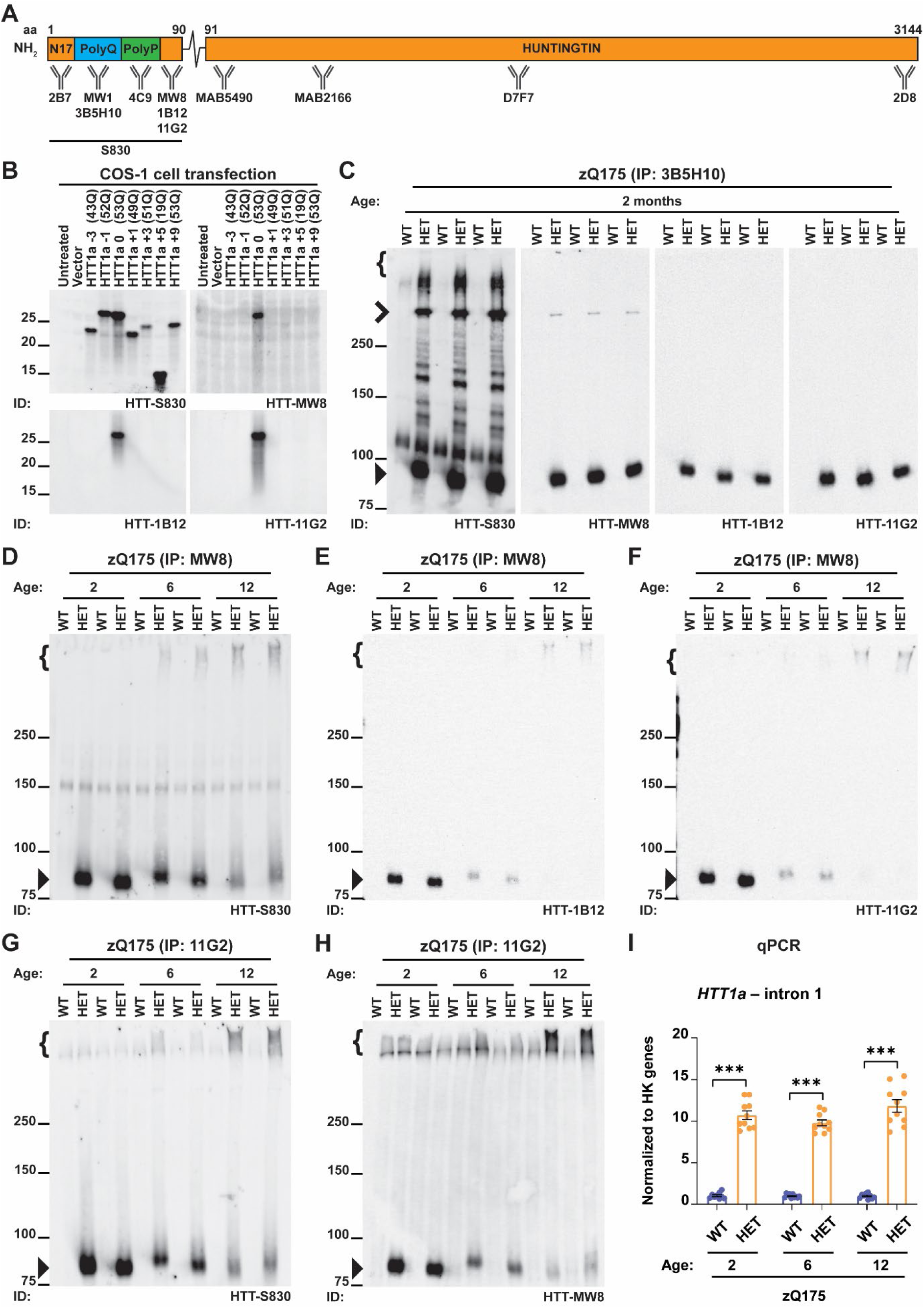
MW8, 1B12 and 11G2 are HTT1a-specific neoepitope antibodies by immunoprecipitation and western blotting. **(A)** Schematic representation of the location of the HTT epitopes recognized by the antibodies used in this study. The HTT protein is 3144 amino acids long and HTT1a extends to amino acid 90 (with 21 CAGs). Details of all immunogens and antibody epitopes are provided in Supplementary Table 1. **(B)** Mutant pSG5-HTT1a constructs (-3 to +9) were transiently expressed in COS-1 cells for 48 h, after which lysates were prepared in HEPES buffer and resolved by 12% Criterion-TGX™ (Bio-Rad) Tris-Glycine SDS-PAGE for western blotting. Proteins were transferred onto nitrocellulose membranes and were probed with either S830, MW8, 1B12 or 11G2 antibodies. An extended version of this experiment is shown (Supplementary Fig. 1). **(C)** Mutant HTT was immunoprecipitated from cortical lysates of zQ175 mice at 2 months of age using the 3B5H10 antibody that detects expanded polyQ repeats. Lysates were prepared and immunoprecipitated in HEPES buffer, resolved by 8% Criterion-TGX™ (Bio-Rad) Tris-Glycine SDS-PAGE, and transferred onto nitrocellulose membrane (0.45 μm) for western blotting. Blots were cut into strips and probed with either S830, MW8, 1B12 or 11G2 antibodies in PBST (n = 3/genotype). An extended version of this experiment is shown (Supplementary Fig. 2). **(D-H)** HTT1a was immunoprecipitated from cortical lysates of zQ175 mice at 2, 6, and 12 months of age using either **(D-F)** MW8 or **(G-H)** 11G2 neoepitope antibodies that specifically recognize the mutant HTT1a protein isoform. Lysates were prepared and immunoprecipitated in HEPES buffer, resolved by 8% Novex™ (Thermo Fisher Scientific) Tris-Glycine SDS-PAGE, and transferred onto nitrocellulose membranes for western blotting (n = 2/genotype/age). For MW8 immunoprecipitations, blots were probed with either **(D)** S830, **(E)** 1B12, or **(F)** 11G2, whilst for 11G2 immunoprecipitations, blots were probed with either **(G)** S830 or **(H)** MW8. **(I)** qPCR for *HTT1a* transcript levels in cDNA prepared from the cortex of wild-type and heterozygous zQ175 mice at 2, 6, and 12 months of age (*n* = 5/gender/genotype/age). Data analysis was by two-way ANOVA with Bonferroni *post hoc* correction. The test statistic, degrees of freedom and *P* values for the ANOVA are reported in Supplementary Table 4. Error bars are mean ± SEM. \*\*\**P* ≤ 0.001. aa = amino acid, N17 = first 17 amino acids, PolyQ = polyglutamine tract, PolyP = proline-rich domain, IP = immunoprecipitate, WT = wild type, HET = heterozygote, } = stacking gel, **>** = full-length mutant HTT, ⏵ = HTT1a, ID = immunodetect, HK = housekeeping. Protein size markers are indicated in kDa.

### Immunoprecipitation and western blotting

Cortical brain lysates were prepared in ice-cold HEPES buffer (50 mM HEPES/NaOH (pH 7.0), 150 mM NaCl, 10 mM EDTA, 1.0% NonidetP-40, 0.5% sodium deoxycholate, 0.1% sodium dodecyl sulphate (SDS), 0.1% bovine serum albumin) supplemented with 10 mM dithiothreitol and cOmplete™ protease inhibitor cocktail tablets (Roche). Tissues were homogenized three times for 30 s using Lysing Matrix D tubes at 6.5 m/s (Fast-Prep-24™, MP Biomedicals), lysates were spun at 13000x*g* for 20 min at 4°C, the supernatant collected, and protein concentrations were determined using the Pierce™ BCA protein assay kit (Thermo Fisher Scientific). Lysates were used immediately and never frozen. For HTT immunoprecipitation, either the anti-polyglutamine 3B5H10 antibody (Sigma-Aldrich) or a HTT1a neoepitope antibody (MW8, 1B12 or 11G2) was used, following previously described protocols.^17,31^ For western blotting, washed immunoprecipitates were resuspended in 2x Laemmli buffer and denatured at 75°C for 10 min. Samples were resolved by either Novex™ (Thermo Fisher Scientific) or Criterion-TGX™ (Bio-Rad) Tris-Glycine SDS-polyacrylamide gel electrophoresis (SDS-PAGE), and transferred onto either nitrocellulose or polyvinylidene difluoride (PVDF) membranes (Bio-Rad) using a submerged transfer system (Bio-Rad) in transfer buffer (25 mM Tris; 192 mM glycine; 20% methanol). Membranes were blocked for 1 h at room temperature in 3% non-fat dried milk in phosphate buffered saline (PBS) containing 0.1% Tween-20 (PBST) and incubated overnight with gentle agitation in primary antibody diluted in 1% PBST at 4°C. After three PBST washes, membranes were incubated with horseradish peroxidase-conjugated secondary antibodies in PBST containing 0.5% non-fat dried milk for 1 h at room temperature, washed again three times in PBST, and detected by chemiluminescence as previously described.^40^ Primary antibodies were used at the following dilutions: S830 (1:2000); 2B7, 4C9, or D7F7 (1:1000); MW8, 1B12 or 11G2 (1:500).

### Cell culture

Sequential site-directed mutagenesis of pSG5-HTT1a was generated as previously described.^31^ COS-1 cells were transiently transfected with 1 μg of DNA using Lipofectamine™ 3000 (Thermo Fisher Scientific), harvested 48 h post-transfection, and lysates were prepared in ice-cold HEPES buffer. For western blot analysis, 25 μg of lysate was resuspended in 2x Laemmli buffer, samples were denatured at 95°C for 10 min and resolved by SDS-PAGE. Primary antibodies were used at the same dilutions as before.

### RNA extractions and real-time quantitative PCR (qPCR)

Cortical brain tissue was homogenized three times for 30 s using Lysing Matrix D tubes at 6.5 m/s (Fast-Prep-24™, MP Biomedicals). Total RNA was extracted using an RNeasy kit (Qiagen), DNase I treated and reverse transcribed with oligodT_18_ (Thermo Fisher Scientific) to amplify the *HTT1a* transcript as previously described.^41^ *HTT1a* primer and probe sequences were: Forward: TCCTCATCAGGCCTAAGAGCTGG, Reverse: GAGACCTCCTAAAAGCATTATGTCATC, Probe: 5’FAM-AGTGCAGGACAGCGTGA-3’BHQ1. Housekeeping (HK) reference genes (*Atp5b, Saha, Eif4a2*) were amplified using custom-designed Taqman assays (Thermo Fisher Scientific). qPCR reactions were performed in triplicate, amplified on a Real-Time PCR system (CFX-Opus384, Bio-Rad), and datasets were analysed as previously described.^42^

### HTT bioassay protein lysate preparation

10% (w/v) cortical brain homogenates were prepared in ice-cold bioassay buffer (PBS containing 1% Triton-X-100) supplemented with cOmplete protease inhibitor cocktail tablets (Roche), as previously described.^20,24^ Lysates were snap frozen, stored overnight at-70°C, and used entirely the following day. For assays detecting aggregated HTT isoforms, crude lysate was used directly. For assays detecting soluble HTT isoforms, lysates were centrifuged at 3500x*g* for 10 min, the supernatant collected and used as previously described.^24^

### Homogenous time resolved fluorescence (HTRF)

Cortical homogenates were dispensed in triplicate into 384-well proxiplates (Greiner Bio-One) to a final volume of 10 μL per well. Lysate dilutions and antibody concentrations are listed (Supplementary Table 2). For HTRF assays, the plate preparation, antibody incubation steps, and detection parameters on an EnVision plate reader (Revvity) were performed as previously described.^20,24^

### Meso Scale Discovery (MSD)

Cortical homogenates were dispensed in triplicate into 384-well, 1-spot custom printed MSD plates to a final volume of 18 μL per well. Lysate dilutions and antibody concentrations are listed (Supplementary Table 3). For MSD assays, the plate blocking, washing, HTT capture, sulfo-tag secondary antibody incubation steps, and detection parameters on a SectorS600 plate reader (Meso Scale Discovery) were performed as previously described.^20^

### Immunohistochemistry

Mice were transcardially perfusion-fixed with 4% paraformaldehyde and tissue processing was as previously described.^40^ Immunohistochemical staining of coronal sections from transgenic N171-82Q or zQ175 knock-in mouse lines was performed as previously described,^30^ except that the ABC reagent was diluted 1:4. Primary antibodies were used at the following dilutions: S830 (1:2000); MW8, 1B12 or 11G2 (1:500). Sections from each experiment were stained together, and images were acquired as previously described.^43^

### Statistical analysis

The data was initially assessed for outliers using Grubbs’ test, and identified outliers were removed prior to any between-group comparisons. Normal Gaussian data distribution was evaluated using the Shapiro-Wilk test and confirmed by Quantile-Quantile plot. Statistical analyses were performed using either Student’s *t*-test, one-way ANOVA, or two-way ANOVA with Bonferroni *post-hoc* correction. All analyses and graphical displays were performed in GraphPad Prism (v10).

### Data availability

The authors confirm that all data supporting the findings of this study are included in this article and its supplementary materials. Raw datasets generated from this study are available from the corresponding author upon request.

## RESULTS

To enhance the detection sensitivity of HTT1a protein isoforms, we evaluated the performance of the MW8 antibody alongside two newly generated recombinant antibodies, 1B12 and 11G2. MW8 is a neoepitope antibody that specifically recognizes the C-terminus of HTT1a that terminates in a proline residue (Supplementary Fig. 1), and does not detect this peptide sequence when present within longer N-terminal proteolytic HTT fragments or full-length HTT protein isoforms.^31^ Like MW8, 1B12 and 11G2 were screened to function as HTT1a-specific neoepitope antibodies.^32^ Because mutant HTT can aggregate in tissue lysates even when stored at −70°C, we implemented a standardized workflow whereby freshly prepared lysates were frozen once and processed within 24 h. The location of the HTT epitopes recognized by the antibodies used in this study is illustrated (Fig. 1A).

### MW8, 1B12 and 11G2 are HTT1a-specific neoepitope antibodies

The MW8 antibody epitope was originally mapped to a region of HTT encompassing the last eight C-terminal amino acids of the HTT1a protein.^44^ Using site-directed mutagenesis of HTT1a constructs, we demonstrated by western blotting that MW8 functions as a neoepitope-specific antibody for detecting the C-terminus of HTT1a.^31^ We subsequently confirmed that MW8 functions as a neoepitope antibody for HTT1a across the HTRF, MSD and AlphaLISA platforms, demonstrating that the 2B7-MW8 assay specifically detected soluble HTT1a, whereas the 4C9-MW8 assay specifically detected aggregated HTT1a.^24,30^

To confirm that the 1B12 and 11G2 antibodies exhibited similar binding properties to MW8, we used our previously established site-directed mutagenesis of HTT1a. In this system, a mammalian expression pSG5-HTT1a construct was sequentially modified by deletion or addition of amino acids relative to the C-terminal proline residue (−3 to +9).^31^ Mutated HTT1a-constructs were transiently expressed in COS-1 cells, lysates were resolved by SDS-PAGE for western blotting, and immunoblotted with either S830, 2B7, 4C9, MW8, 1B12 or 11G2. Consistent with previous findings,^31^ S830, 2B7 and 4C9 detected all mutant constructs. In contrast, 1B12 and 11G2, like MW8, detected only the HTT1a protein terminating at the native C-terminal proline residue and failed to detect the −1, +1, or any other mutant constructs, demonstrating that they function as neoepitope-specific antibodies for soluble HTT1a (Fig. 1B and Supplementary Fig. 1).

To determine whether the 1B12 and 11G2 antibodies function as neoepitope-specific antibodies for detecting soluble and aggregated HTT1a *in vivo*, we applied an established HTT immunoprecipitation coupled with western blotting workflow. This approach had been previously applied to characterize how the pattern of mutant full-length HTT, N-terminal proteolytic HTT fragments, and HTT1a protein species change during disease progression.^31^ In zQ175 mice, endogenous mouse exon 1 *Htt* has been substituted with mutant human exon 1 *HTT.*^33,45^ Mutant HTT was immunoprecipitated from cortical lysates of zQ175 mice at 2 months of age using 3B5H10, which selectively binds soluble polyQ-expanded tracts, resolved by SDS-PAGE for western blotting, and immunoblotted with either S830, MW8, 1B12 or 11G2. S830 is a polyclonal antibody raised against human HTT1a(53Q) and detected full-length HTT, N-terminal proteolytic HTT fragments and HTT1a (Fig. 1C). MW8 detected a single band at approximately 90 kDa, confirming specific recognition of the soluble HTT1a protein. Strikingly, 1B12 and 11G2 also detected this single band, demonstrating that they function as neoepitope-specific antibodies for soluble HTT1a *in vivo* (Fig. 1C). During SDS-PAGE optimisation, we compared the effect of transferring gels to either nitrocellulose or PVDF membrane resins. Unexpectedly, and contrary to the detection of full-length HTT and N-terminal proteolytic HTT fragments, HTT1a failed to bind PVDF membranes (Supplementary Fig. 2). Based on these findings, we recommend that PVDF resins should not be used for the western blotting of HTT proteins.

To further investigate the sensitivity of the MW8, 1B12 and 11G2 antibodies for detecting HTT1a by western blot, reciprocal immunoprecipitation experiments were performed from cortical lysates of zQ175 mice at 2 months of age (Supplementary Fig. 3). Relative to the mouse immunoglobulin G (IgG) control, all three antibodies successfully immunoprecipitated soluble HTT1a, validating their neoepitope specificity. However, the most effective conditions for immunoprecipitating and detecting HTT1a with minimal nonspecific background was achieved when reciprocal species-specific antibody combinations were used (i.e., immunoprecipitation with mouse MW8 followed by immunoblotting with either rabbit 1B12 or 11G2, or *vice versa*, Supplementary Fig. 3). Applying these optimized conditions, HTT1a was then immunoprecipitated from cortical lysates of zQ175 mice at 2, 6, and 12 months of age to investigate how HTT1a protein isoforms changed during disease pathogenesis (Fig. 1D-H). When immunoblotting with S830 following MW8 (Fig. 1D) or 11G2 (Fig. 1G) immunoprecipitation, soluble HTT1a levels detected in the resolving gel progressively decreased, whereas the appearance of insoluble aggregated HTT1a in the stacking gel increased from 2 to 12 months of age. A similar pattern was observed when MW8 immunoprecipitates were immunoblotted with either 1B12 (Fig. 1E) or 11G2 (Fig. 1F), or when 11G2 immunoprecipitates were immunoblotted with MW8 (Fig. 1H). Notably at 6 and 12 months of age, MW8 was more sensitive than 11G2 for detecting the decreasing levels of soluble HTT1a and increasing abundance of aggregated HTT1a.

On occasion, a faint full-length mutant HTT signal was detected above the IgG control when samples were either immunoprecipitated or probed with HTT1a-specific antibodies by western blotting (Fig. 1C, MW8 panel and Supplementary Fig. 3A, MW8, 1B12 and 11G2 immunoprecipitates), prompting further investigation. To assess this, HTT1a was immunoprecipitated from cortical lysates of zQ175 mice at 2 months of age using MW8, 1B12 or 11G2, resolved by SDS-PAGE for western blotting, and immunoblotted with either S830, 2B7, or D7F7 (Supplementary Fig. 4). Consistent with previous findings (Supplementary Fig. 3), all three HTT1a-specific antibodies immunoprecipitated soluble HTT1a above IgG control levels, and when immunoblotted with S830 a small amount of full-length mutant HTT signal was detected (Supplementary Fig. 4A, 1B12 and 11G2 immunoprecipitates). When immunoblotted with 2B7 or D7F7, a small amount of full-length HTT was detected in both wild-type and heterozygous zQ175 samples but was absent in IgG controls (Supplementary Fig. 4B, C). Together, these results demonstrate that under highly enriched immunoprecipitation conditions, MW8, 1B12 and 11G2 antibodies can bind a minimal amount of full-length HTT (wild-type and mutant), but this affinity is likely to be negligible in comparison to their functional specificity as HTT1a neoepitope antibodies.

### Reduction in soluble HTT1a levels cannot be attributed to decreased *HTT1a* transcript levels

To determine whether the reduction in soluble HTT1a levels observed by western blot during disease progression was attributable to its recruitment into detergent-insoluble HTT1a aggregates, we quantified *HTT1a* transcript levels. Cortical cDNA from heterozygous zQ175 mice was generated at 2, 6, and 12 months of age, alongside age-matched wild-type littermate controls and analyzed by real-time quantitative PCR (qPCR). Consistent with previous findings, ^17,20^ the *HTT1a* transcript was only produced in mutant knock-in mice and was absent in wild-type controls. Importantly, a notable change in steady-state *HTT1a* transcript levels was not observed, confirming that the reduction in soluble HTT1a levels during disease progression cannot be attributed to a decrease in *HTT1a* transcript production (Fig. 1I).

### Enhancing the sensitivity of HTRF and MSD assays that detect soluble HTT1a

We previously demonstrated that because MW8 functions as a HTT1a-specific neoepitope antibody, it could be used to develop a 2B7-MW8 assay capable of selectively detecting the soluble HTT1a isoform in zQ175 mice across three technology platforms (HTRF, MSD and AlphaLISA).^24^ In an allelic series of knock-in mouse lines (*Hdh*Q20, *Hdh*Q50, *Hdh*Q80, *Hdh*Q111, CAG140 and zQ175) we subsequently showed that although the 2B7-MW8 HTRF assay was sufficiently sensitive to detect increasing soluble HTT1a levels with increasing CAG repeat length, that the equivalent MSD assay was weak and not technically viable.^20^ Therefore, we sought to assess whether substituting MW8 with either 1B12 or 11G2 could improve assay sensitivity for the detection of soluble HTT1a by HTRF and MSD.

To support this evaluation, we included transgenic N171-82Q mice that express a cDNA transgene encoding the first 171 amino acids of HTT.^46^ Because these mice lack *HTT* intron 1, they cannot generate the *HTT1a* transcript and consequently do not produce HTT1a, thereby serving as a biologically relevant negative control for HTT1a-specific assays. In addition to zQ175 mice, to further challenge assay sensitivity, we incorporated transgenic YAC128 mice into our experimental design. ^47^ YAC128 mice express full-length human *HTT* with an expanded CAG repeat encoding 125 glutamines and produces HTT1a at substantially lower levels than zQ175 mice.^20^

Cortical lysates from zQ175 mice at 2, 6, and 12 months of age, and from N171-82Q mice at 1.5 and 3 months of age, alongside age-matched wild-type littermate controls were analyzed using the HTRF and MSD platforms (Fig. 2). To verify sample genotypes, ‘total soluble mutant HTT’ levels were initially assessed using the 2B7-MW1 assay, which detects all soluble mutant HTT isoforms, including full-length HTT, HTT1a, the N171-82Q transgene protein, and any N-terminal proteolytic HTT fragments. Consistent with previously published data,^24^ both HTRF (Fig. 2A) and MSD (Fig. 2B) assay measurements revealed a continual decline in total soluble mutant HTT protein levels with disease progression, with no signal detected in wild-type controls.

**Figure 2.**
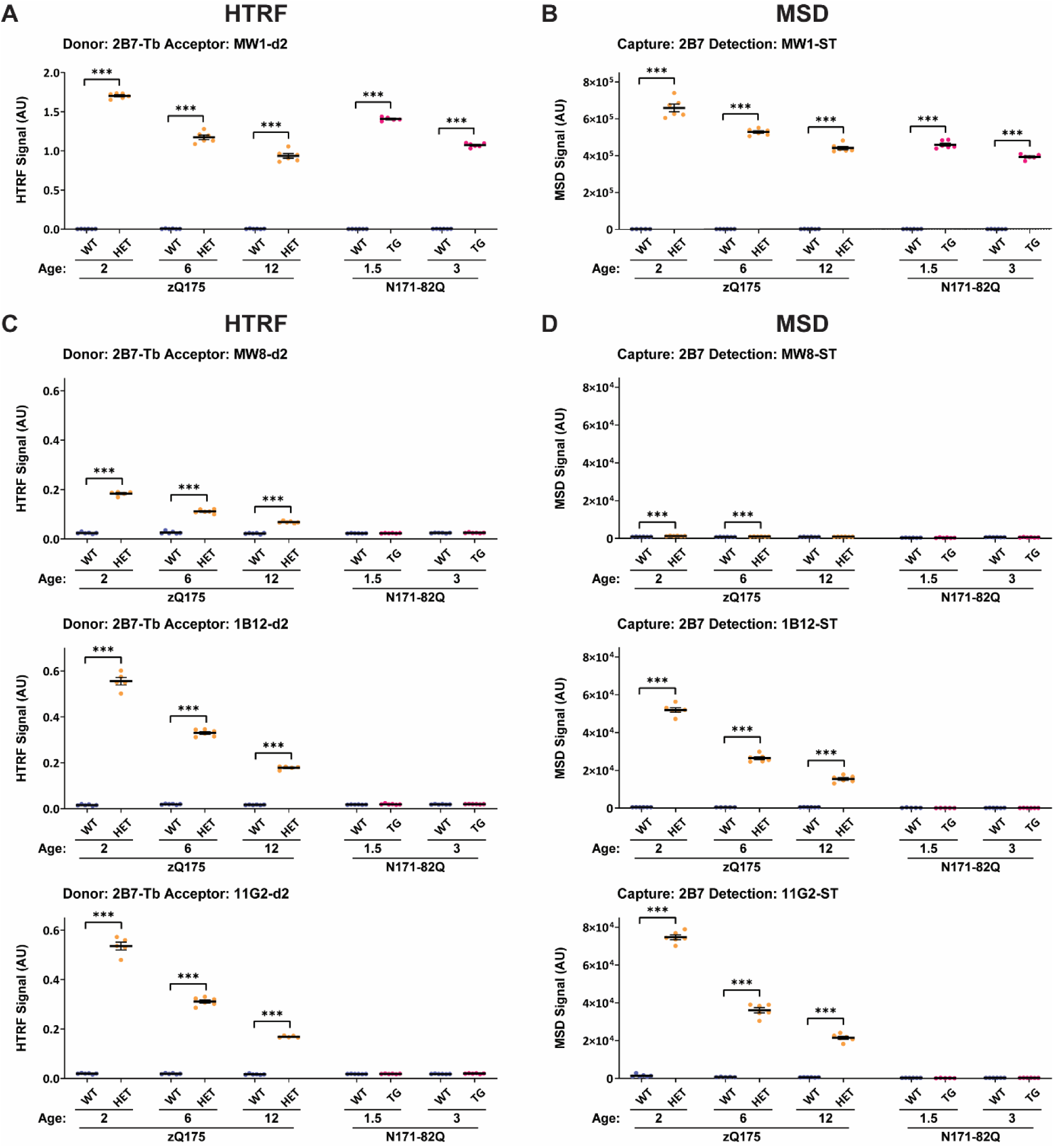
Enhancing the detection of soluble HTT1a by HTRF and MSD. (A-B) To verify sample genotypes, the 2B7-MW1 ‘total mutant HTT’ assay was evaluated by HTRF **(A)** and MSD **(B)** using cortical lysates from heterozygous zQ175 mice at 2, 6, or 12 months of age, and from transgenic N171-82Q mice at 1.5 and 3 months, alongside age-matched wild-type littermate controls. **(C-D)** To detect soluble HTT1a, antibody pairings of 2B7-MW8, 2B7-1B12 and 2B7-11G2 were assessed by HTRF **(C)** and MSD **(D)** using the same cortical lysates (n = 3/gender/genotype/age). HTT1a was not detected in transgenic N171-82Q or wild-type mice, which do not produce the HTT1a protein. Per dataset, the *y*-axis legends were scaled the same. Data analysis was by two-way ANOVA with Bonferroni *post hoc* correction. The test statistic, degrees of freedom and *P* values for the ANOVA are reported in Supplementary Table 5. Error bars are mean ± SEM. \*\*\**P* ≤ 0.001. Tb = terbium cryptate, ST = sulfo-tag, AU = arbitrary units, WT = wild type, HET = heterozygote, TG = transgenic.

To evaluate soluble HTT1a detection we performed HTRF with 2B7 as the donor antibody (Fig. 2C) and MSD with 2B7 as the capture antibody (Fig. 2D) paired with either MW8, 1B12 or 11G2, and for comparative purposes, each dataset was scaled to whichever antibody pairing had the highest signal. HTT1a was not detected in wild-type samples. In zQ175 mice at 2 months of age, substituting MW8 with either 1B12 or 11G2 resulted in an approximate 3-fold improvement in HTRF assay signal, with 1B12 being marginally more sensitive than 11G2 (Fig. 2C). In contrast, substituting MW8 with either 1B12 or 11G2 on the MSD platform substantially enhanced assay performance, whereby 11G2 was significantly more sensitive than 1B12 and resulted in an approximate 60-fold improvement in signal (Fig. 2D). Across both assay platforms, soluble HTT1a levels continually decreased from 2 to 12 months of age in zQ175 mice and tracked with disease progression (Fig. 2C, D), consistent with our immunoprecipitation and western blot findings (Fig. 1D-H). Importantly, HTT1a was not detected in N171-82Q lysates by any assay configuration, confirming specificity for detecting soluble HTT1a (Fig. 2C, D).

To further evaluate assay sensitivity, cortical lysates from YAC128 mice at 2, 6, and 12 months of age, and from N171-82Q mice at 1.5 and 3 months of age, alongside age-matched wild-type littermate controls were analyzed using the HTRF and MSD platforms (Supplementary Fig. 5). The 2B7-MW1 ‘total soluble mutant HTT’ assay confirmed that both YAC128 and N171-82Q mice expressed mutant HTT by HTRF (Supplementary Fig. 5A) and MSD (Supplementary Fig. 5B). The detection of soluble HTT1a by each assay configuration remained consistent with observations in zQ175 lysates (Fig. 2), and substituting MW8 with either 1B12 or 11G2 significantly enhanced assay performance across both technology platforms (Supplementary Fig. 5C, D). Furthermore, as detectable soluble HTT1a levels continually decreased from 2 to 12 months of age in YAC128 mice and tracked disease progression, a viable MSD assay had been generated (Supplementary Fig. 5D).

### HTRF and MSD assays with reciprocal 2B7 pairings detect either soluble or aggregated HTT1a

We previously demonstrated that reversing the soluble HTT1a antibody pairings (2B7-Tb: MW8-d2 for HTRF or 2B7-Capture: MW8-ST for MSD) to the opposite orientation (MW8-Tb: 2B7-d2 for HTRF or MW8-Capture: 2B7-ST for MSD) produced assays that tracked with an aggregated form of HTT1a in zQ175 mice, whereby signal intensity increased with disease progression.^24^ To further investigate this observation, cortical lysates from zQ175 and N171-82Q mice were analysed using the HTRF and MSD platforms (Fig. 3).

**Figure 3.**
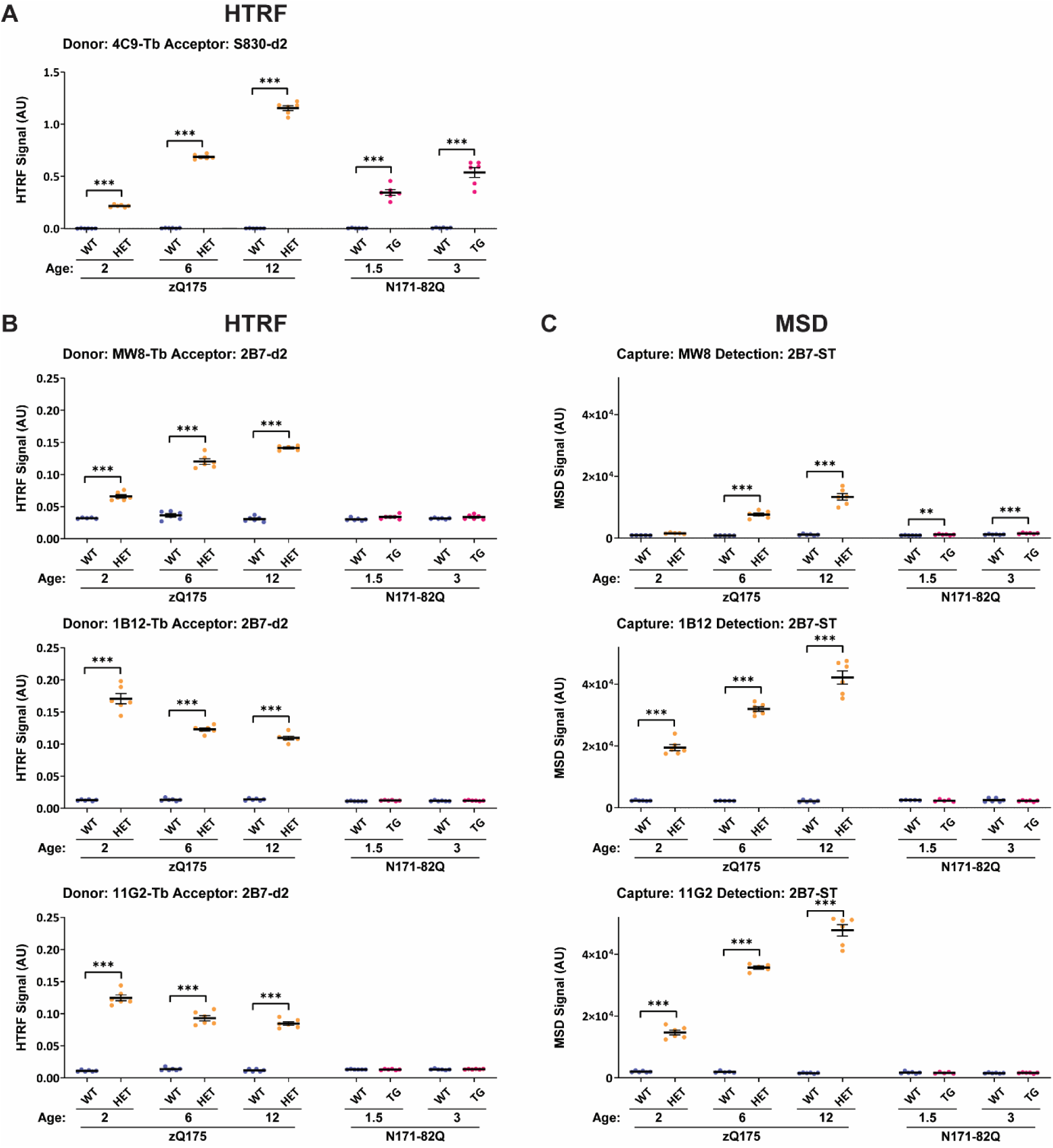
Detection of soluble or aggregated HTT1a protein isoforms with reciprocal 2B7 pairings by HTRF and MSD. **(A)** To verify sample genotypes, the 4C9-S830 ‘total HTT aggregation’ assay was evaluated by HTRF using cortical lysates from heterozygous zQ175 mice at 2, 6, or 12 months of age, and from transgenic N171-82Q mice at 1.5 and 3 months, alongside age-matched wild-type littermate controls. **(B-C)** To detect HTT1a protein isoforms, antibody pairings of MW8-2B7, 1B12-2B7 and 11G2-2B7 were assessed by HTRF **(B)** and MSD **(C)** using the same cortical lysates (n = 3/gender/genotype/age). HTT1a was not detected in wild-type mice. Besides a small but significant signal in the MW8-2B7 MSD assay, HTT1a was not detected in transgenic N171-82Q mice, which do not produce the HTT1a protein. Per dataset, the *y*-axis legends were scaled the same. Data analysis was by two-way ANOVA with Bonferroni *post hoc* correction. The test statistic, degrees of freedom and *P* values for the ANOVA are reported in Supplementary Table 6. Error bars are mean ± SEM. \*\**P* ≤ 0.01 and \*\*\**P* ≤ 0.001. Tb = terbium cryptate, ST = sulfo-tag, AU = arbitrary units, WT = wild type, HET = heterozygote, TG = transgenic.

We first assessed, ‘total HTT aggregation’ levels by using the 4C9-S830 HTRF assay, which confirmed that aggregated HTT levels increased progressively with disease progression (Fig. 3A). We then performed HTRF with 2B7 as the acceptor antibody (Fig. 3B) and MSD with 2B7 as the detection antibody (Fig. 3C) paired with either MW8, 1B12 or 11G2, and for comparative purposes, each dataset was scaled to whichever antibody pairing had the highest signal. HTT1a was not detected in wild-type samples. Consistent with previous findings,^24^ in zQ175 mice the MW8-2B7 HTRF assay signal tracked with an aggregated HTT1a isoform. However, when MW8 was substituted with either 1B12 or 11G2, HTRF assay signals continued to track with soluble HTT1a (Fig. 3B). Notably, the HTRF signal intensities for both the 1B12-2B7 and 11G2-2B7 configurations were lower when compared to previous soluble HTT1a measurements (Fig. 2C). In contrast, substituting MW8 with either 1B12 or 11G2 on the MSD platform continued to produce assay signals that tracked with aggregated HTT1a, with a significantly improved assay window. At 2 months of age, signal intensities were observed above wild-type controls, whilst by 12 months, an approximate 3.5-fold improvement in signal was observed (Fig. 3C). Across both assay platforms, when assays detected an aggregated HTT1a isoform, levels continually increased from 2 to 12 months of age in zQ175 mice and tracked with disease progression (Fig. 3B, C). An absence of signal in N171-82Q lysates confirmed assay specificity for detecting the HTT1a isoforms (Fig. 3B, C).

As before, an additional dataset of cortical lysates from YAC128 and N171-82Q mice were analyzed using the HTRF and MSD platforms to further evaluate assay sensitivity (Supplementary Fig. 6). The 4C9-S830 ‘total HTT aggregation’ assay confirmed that aggregated HTT levels increased progressively in YAC128 and N171-82Q mice by HTRF (Supplementary Fig. 6A). The detection of soluble versus aggregated HTT1a isoforms by each assay configuration remained consistent with observations in zQ175 lysates (Fig. 3), and substituting MW8 with either 1B12 or 11G2 significantly enhanced assay performance on the MSD platform to detect aggregated HTT1a (Supplementary Fig. 6C).

### MW8, 1B12 and 11G2 are HTT1a-specific neoepitope antibodies by immunohistochemistry

We previously optimized our immunohistochemistry protocols that used the S830 polyclonal and MW8 monoclonal antibodies to investigate the temporal and spatial emergence of HTT aggregation in the zQ175 brain from 1 to 6 months of age.^30^ Using this protocol, coronal sections from heterozygous zQ175 mice and transgenic N171-82Q mice were immunostained with either S830, MW8, 1B12 or 11G2.

To determine whether the MW8, 1B12 and 11G2 antibodies function as neoepitope antibodies for detecting aggregated HTT1a by immunohistochemistry, tissue sections from zQ175 mice at 6 months of age and from N171-82Q mice at 3 months of age were immunostained, alongside age-matched wild-type littermate controls. Because N171-82Q mice express a cDNA transgene encoding the first 171 amino acids of HTT and lack *HTT* intron 1, they cannot generate the *HTT1a* transcript and consequently do not produce HTT1a, thereby serving as a biologically relevant negative control. In sections from 6-month-old zQ175 mice, HTT aggregation was readily detected with all four antibodies (Fig. 4). In sections from 3-month-old N171-82Q, S830 immunostaining detected diffuse HTT aggregation that filled cellular nuclei of the striatum and cortex (Fig. 4), and of the hippocampus and cerebellum (Supplementary Fig. 7), with no evidence of nuclear or cytoplasmic inclusions (Fig. 4 and Supplementary Fig. 7). In contrast, immunostaining with either MW8, 1B12 or 11G2 failed to detect any HTT aggregation in these brain regions and appeared indistinguishable from wild-type littermate controls (Fig. 4 and Supplementary Fig. 7), demonstrating that MW8, 1B12 and 11G2 function as neoepitope-specific antibodies for detecting aggregated HTT1a by immunohistochemistry.

**Figure 4.**
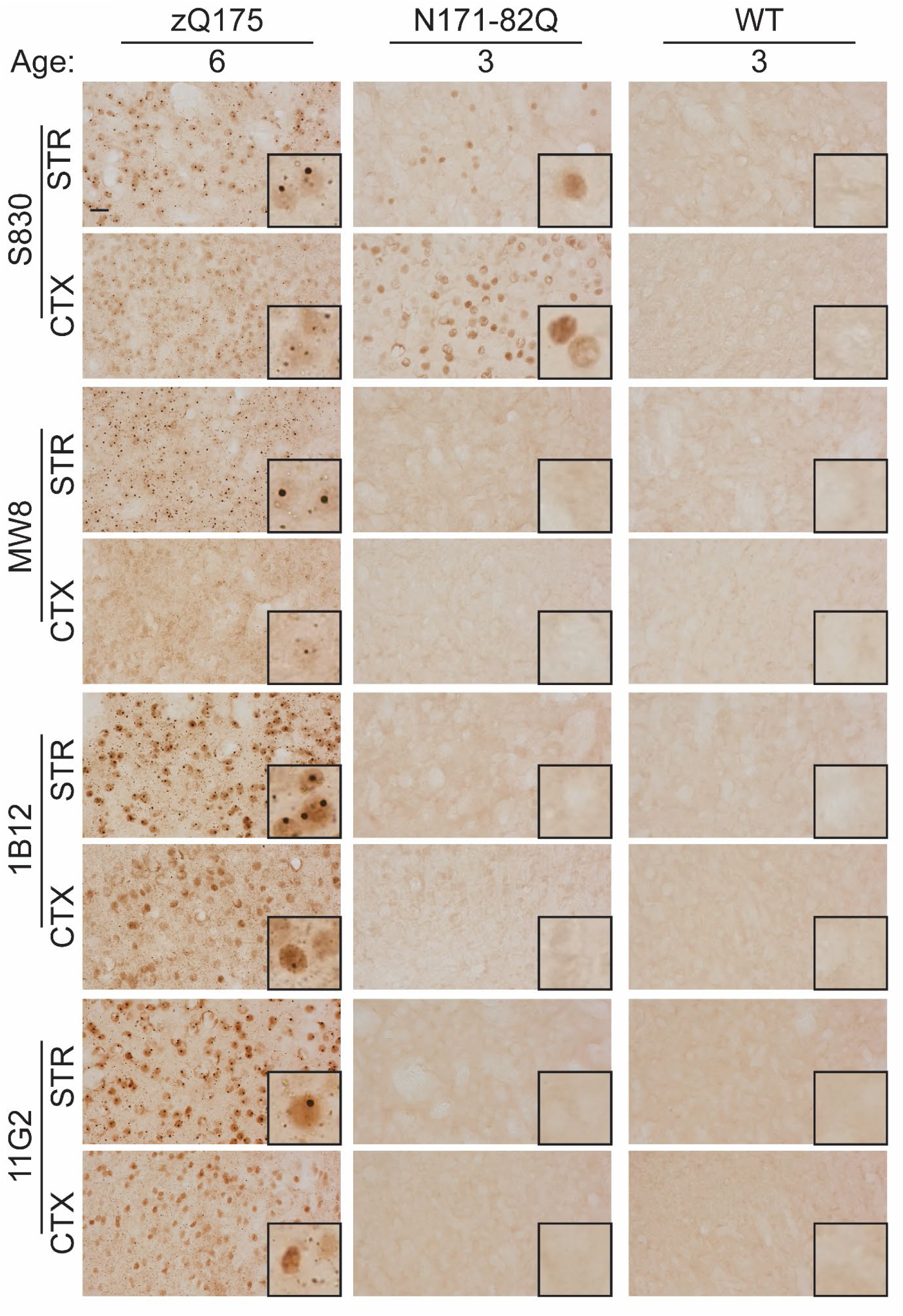
MW8, 1B12 and 11G2 are HTT1a-specific neoepitope antibodies by immunohistochemistry. Coronal brain sections at the level of the striatum and cortex (layers I-III) from heterozygous zQ175 mice at 6 months of age, and from transgenic N171-82Q mice at 3 months of age, alongside age-matched wild-type littermate controls were immunostained with either S830, MW8, 1B12 or 11G2 (n = 3/genotype). A HTT aggregation signal was not detected in wild-type mice. In zQ175 mice at 6 months, all four antibodies detected a diffuse nuclear aggregated HTT signal in addition to nuclear and cytoplasmic inclusion bodies in both the striatum and cortex. In N171-82Q mice at 3 months, S830 detected a diffuse nuclear aggregated HTT signal in the striatum and cortex, whereas MW8, 1B12 or 11G2 did not detect any aggregated HTT, with sections appearing indistinguishable from wild-type littermate controls. A nuclear counterstain was not applied to these sections as this would mask the nuclear signal. An extended version of this experiment with staining at the level of the hippocampus (CA1 and hilus regions) and cerebellum is shown (Supplementary Fig. 7). Scale bar = 20 μm, zoomed boxed area = 20 μm^2^. WT = wild type, STR = striatum, CTX = cortex.

To compare the patterns of HTT1a aggregate immunostaining detected by MW8, 1B12 and 11G2 antibodies, tissue sections at the level of the striatum and cortex from heterozygous zQ175 mice and wild-type littermates at 2, 6, and 12 months of age were immunostained, alongside S830. Consistent with previous findings,^30^ by 2 months of age, the S830 antibody detected a diffuse HTT aggregation signal that filled the nuclei of the striatum, with nuclear inclusions present in some cells. By 6 months of age, the diffuse HTT aggregation signal was more pronounced, with nuclear inclusions present in most cells and a substantial increase in the density of cytoplasmic inclusions, whilst by 12 months, both nuclear and cytoplasmic inclusions had increased in size (Fig. 5). In the cortex, a similar staining pattern was observed, however, relative to the striatum, detection of HTT aggregation was delayed (Fig. 5). Immunostaining with MW8 was markedly weaker at 2 and 6 months of age, as MW8 failed to detect the diffuse nuclear signal, but by 12 months was comparable to S830 (Fig. 5). In contrast, immunostaining with 1B12 or 11G2 was more pronounced than with S830 in both the striatum and cortex at all ages examined (Fig. 5). In wild-type control sections, no diffuse nuclear HTT aggregation signal or inclusions were observed (Fig. 5). Collectively, these findings demonstrate that, by immunohistochemistry, both 1B12 and 11G2 HTT1a-specific antibodies were more sensitive than MW8 to detect and track HTT1a aggregation.

**Figure 5.**
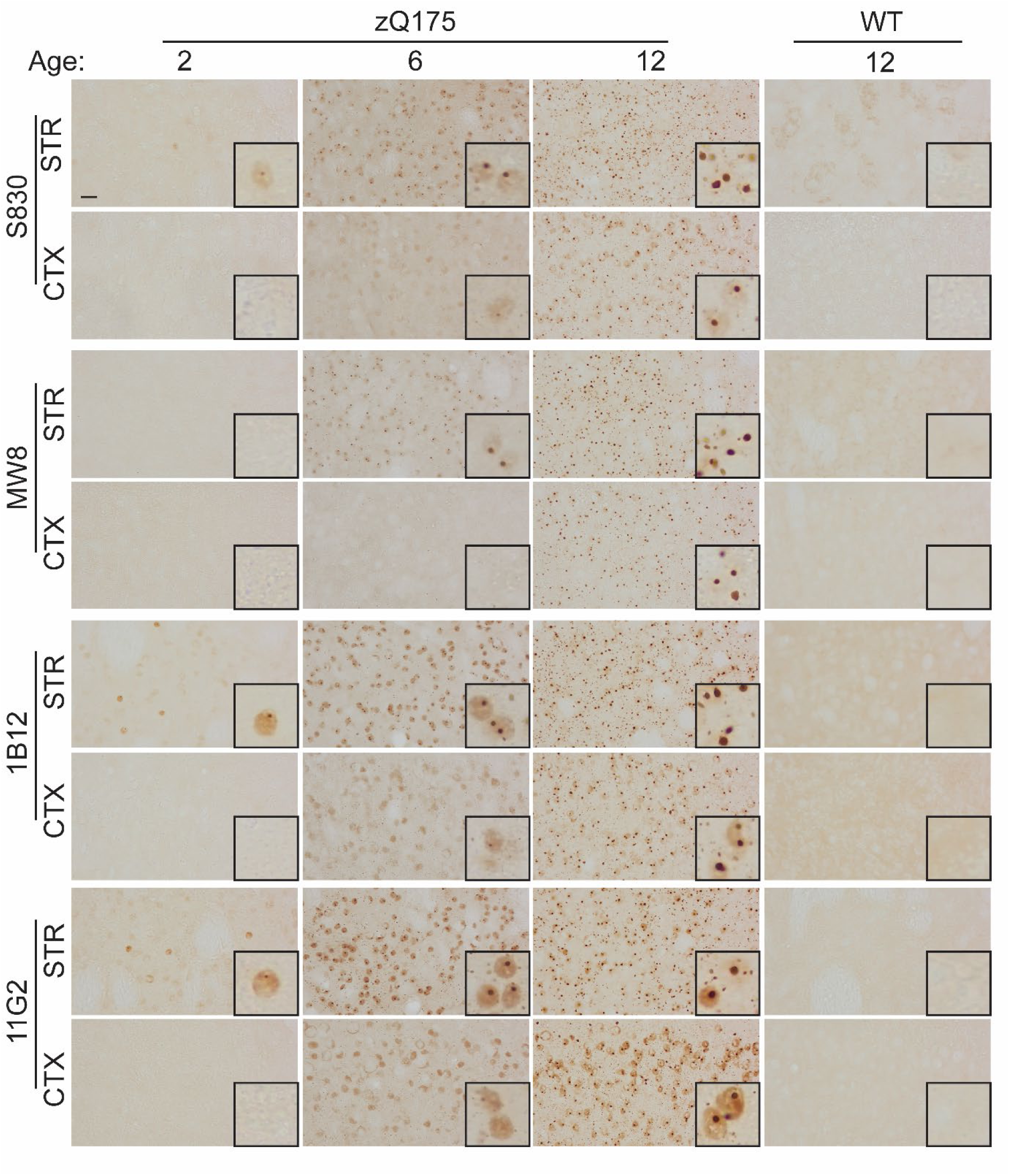
HTT1a aggregates accumulate in the brains of zQ175 mice by immunohistochemistry. Coronal brain sections at the level of the striatum and cortex (layers I-III) from wild-type and heterozygous zQ175 mice were immunostained with either S830, MW8, 1B12 or 11G2 at 2, 6, and 12 months of age (n = 3/genotype/age). A HTT1a aggregation signal was not detected in wild-type mice. With S830, occasional nuclear HTT inclusions and a diffuse nuclear aggregation staining pattern were detected in the striatum as early as 2 months of age and increased markedly between 6 and 12 months. Smaller cytoplasmic inclusions appeared from 6 months of age, and by 12 months, both nuclear and cytoplasmic inclusions had increased in size. In the cortex (layers I-III), aggregated HTT appeared later, becoming detectable by 6 months of age and increased substantially by 12 months. Staining with 1B12 and 11G2 produced a staining pattern similar to S830 but with a stronger signal intensity, whereas MW8 staining was weaker. A nuclear counterstain was not applied to these sections as this would mask the nuclear signal. Scale bar = 20 μm, zoomed boxed area = 20μm^2^. WT = wild type, STR = striatum, CTX = cortex.

### Enhancing the sensitivity of HTRF and MSD assays that detect aggregated HTT1a

The 4C9 antibody is specific for the human polyproline-rich domain in *HTT* exon 1 and has been used to develop mutant HTT-specific assays in mouse knock-in lines that carry a humanized exon 1.^23^ Previously, when developing HTT assays we demonstrated that pairing 4C9 with MW8 consistently tracked with HTT aggregation across three technology platforms (HTRF, MSD and AlphaLISA),^24^ and established that the 4C9-MW8 HTRF assay was specific for aggregated HTT1a.^30^ To comprehensively evaluate the detection of aggregated HTT1a, cortical lysates from zQ175 mice at 2, 6, and 12 months of age, and from N171-82Q mice at 1.5 and 3 months of age, alongside age-matched wild-type littermate controls, were analysed using the HTRF and MSD platforms (Fig. 6).

**Figure 6.**
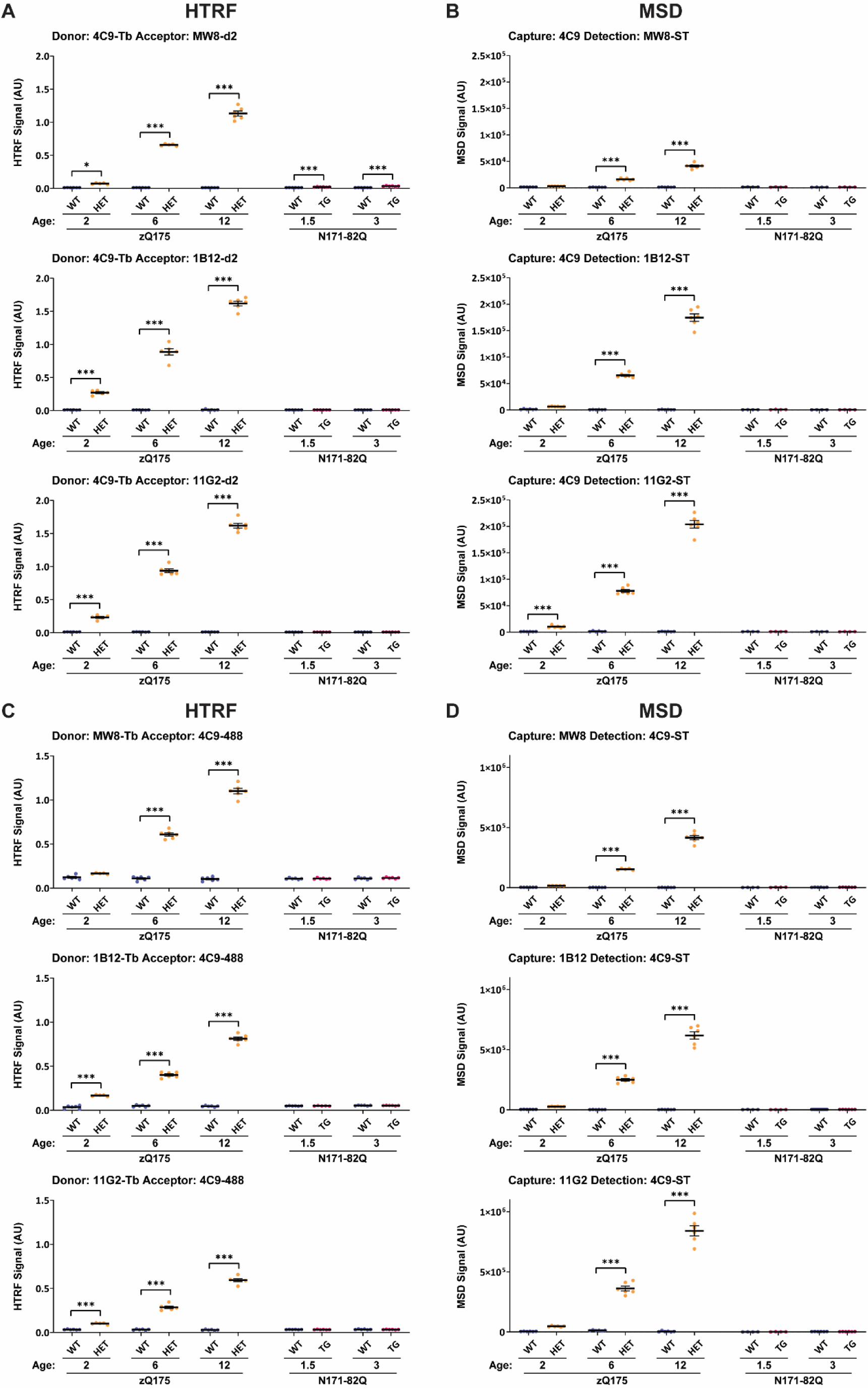
Enhancing the detection of aggregated HTT1a by HTRF and MSD. (A-D) To detect aggregated HTT1a, reciprocal antibody pairings of 4C9-MW8, 4C9-1B12 and 4C9-11G2 were assessed by HTRF **(A)** and MSD **(B)** and of MW8-4C9, 1B12-4C9 and 11G2-4C9 were assessed by HTRF **(C)** and MSD **(D).** Assays were performed using cortical lysates from heterozygous zQ175 mice at 2, 6, or 12 months of age, and from transgenic N171-82Q mice at 1.5 and 3 months, alongside age-matched wild-type littermate controls (n = 3/gender/genotype/age). Any signal detected in wild-type mice by each assay configuration represented assay background, since the endogenous mouse wild-type HTT protein does not contain the human HTT-specific sequences recognized by the 4C9 epitope. Besides a small but significant signal in the 4C9-MW8 HTRF assay, HTT1a was not detected in transgenic N171-82Q mice, which do not produce the HTT1a protein. Per dataset, the *y*-axis legends were scaled the same. Data analysis was by two-way ANOVA with Bonferroni *post hoc* correction. The test statistic, degrees of freedom and *P* values for the ANOVA are reported in Supplementary Table 7. An extended version of this experiment, further demonstrating HTT1a assay exclusivity is shown (Supplementary Fig. 9). Error bars are mean ± SEM. \**P* ≤ 0.5 and \*\*\**P* ≤ 0.001. Tb = terbium cryptate, ST = sulfo-tag, AU = arbitrary units, WT = wild type, HET = heterozygote, TG = transgenic.

HTRF was performed with 4C9 as the donor antibody (Fig. 6A) and MSD with 4C9 as the capture antibody (Fig. 6B) paired with either MW8, 1B12 or 11G2, together with reciprocal assays of HTRF with 4C9 as the acceptor antibody (Fig. 6C) and MSD with 4C9 as the detection antibody (Fig. 6D) paired with either MW8, 1B12 or 11G2. For comparative purposes, each dataset was scaled to whichever antibody pairing had the highest signal, and HTT1a was not detected in wild-type samples. In zQ175 mice, all assay configurations were specific for mutant HTT and correlated with increasing levels of aggregated HTT1a (Fig. 6). By HTRF, the greatest signal and lowest background was observed when 4C9 served as the terbium donor (Fig. 6A, C). In zQ175 mice at 2 months of age, substituting MW8 with either 1B12 or 11G2 resulted in a significantly improved assay window to detect aggregated HTT1a. Whilst by 12 months, an approximate 1.5-fold improvement assay signal was observed, with 1B12 being marginally more sensitive than 11G2 (Fig. 6A). By MSD, the greatest signal and lowest background was observed when 4C9 served as the sulfo-tag detector (Fig. 6B, D). In zQ175 at 2 months of age, substituting MW8 with either 1B12 or 11G2 also resulted in a significantly improved assay window to detect aggregated HTT1a, with 11G2 being significantly more sensitive than 1B12. Relative to our previously established MW8-4C9 assay, by 12 months, an approximate 2-fold improvement in signal was observed when substituting MW8 with 11G2 (Fig. 6D). Across both assay platforms, aggregated HTT1a levels continually increased from 2 to 12 months of age in zQ175 mice and tracked with disease progression (Fig. 6), consistent with our immunoprecipitation and western blot findings (Fig. 1D-H). An absence of signal in N171-82Q lysates confirmed assay specificity for detecting aggregated HTT1a (Fig. 6).

Once again, cortical lysates from YAC128 and N171-82Q mice were analyzed using the HTRF and MSD platforms to further evaluate assay sensitivity (Supplementary Fig. 8). The detection of aggregated HTT1a by each assay configuration remained consistent with observations in zQ175 lysates (Fig. 6), and substituting MW8 with either 1B12 or 11G2 significantly enhanced assay performance across both technology platforms (Supplementary Fig. 8). Finally, because the 4C9, MW8, 1B12 and 11G2 antibodies recognize epitopes that are present in full-length mutant HTT, we reasoned that exclusive HTT1a-specific aggregation assays should not detect any background signal in the *Hdh*Q20 knock-in line. This model carries a polyQ expansion of 20, which does not produce aggregated HTT1a, but does express full-length mutant HTT containing these epitopes, which could therefore be detected.^20^ To evaluate this and confirm HTT1a assay specificity, the same HTRF and MSD assays were reassessed on cortical lysates from wild-type, heterozygous and homozygous *Hdh*Q20 and zQ175 knock-in mice, alongside transgenic YAC128 mice and wild-type littermate controls, at 2 months of age (Supplementary Fig. 9).

### HTT fragments longer than HTT1a are incorporated into HTT1a aggregates

We previously demonstrated that pairing MW8 with either MAB5490 or MAB2166 produced assays capable of tracking HTT aggregation in zQ175 mice, whereby signal intensities increased with disease progression.^24^ To further investigate this phenomenon with enhanced sensitivity, we repeated these experiments using 1B12 and 11G2, alongside MW8, whilst simultaneously expanding our antibody panel beyond MAB5490 and MAB2166 to include D7F7 and 2D8, thereby increasing epitope coverage across the entire full-length HTT protein (Fig. 1A). Cortical lysates from zQ175 mice at 2, 6, and 12 months of age, alongside age-matched wild-type littermate controls, were analysed using both the HTRF (Supplementary Fig. 10) and MSD platforms (Fig. 7).

**Figure 7.**
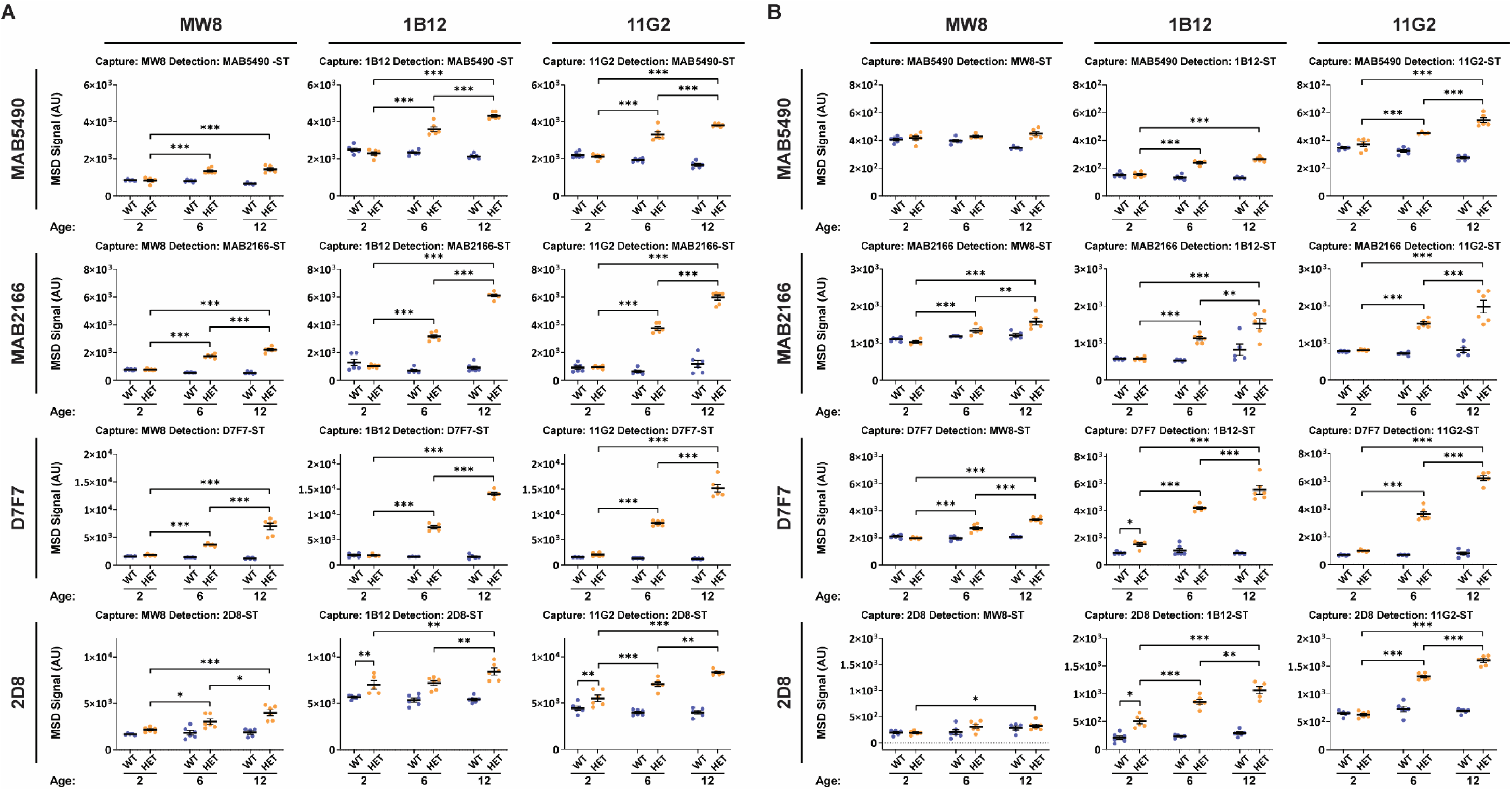
Detection of HTT fragments longer than HTT1a in HTT1a-containing aggregates by MSD. (A-B) Reciprocal antibody pairings of **(A)** MW8-MAB5490, 1B12-MAB5490, 11G2-MAB5490, MW8-MAB2166, 1B12-MAB2166, 11G2-MAB2166, MW8-D7F7, 1B12-D7F7, 11G2-D7F7, MW8-2D8, 1B12-2D8, 11G2-2D8 and of **(B)** MAB5490-MW8, MAB5490-1B12, MAB5490-11G2, MAB2166-MW8, MAB2166-1B12, MAB2166-11G2, D7F7-MW8, D7F7-1B12, D7F7-11G2, 2D8-MW8, 2D8-1B12, 2D8-11G2 were assessed by MSD. Assays were performed using cortical lysates from heterozygous zQ175 mice at 2, 6, or 12 months of age, alongside age-matched wild-type littermate controls (n = 3/gender/genotype/age). Per dataset, the *y*-axis legends were scaled the same. Data analysis was by two-way ANOVA with Bonferroni *post hoc* correction. The test statistic, degrees of freedom and *P* values for the ANOVA are reported in Supplementary Table 10. Error bars are mean ± SEM. \**P* ≤ 0.5, \*\**P* ≤ 0.01 and \*\*\**P* ≤ 0.001. ST = sulfo-tag, AU = arbitrary units, WT = wild type, HET = heterozygote.

Reciprocal HTRF assays were performed with either MW8, 1B12 or 11G2 as the donor (Supplementary Fig. 10A) or acceptor (Supplementary Fig. 10B) antibody, paired with either MAB5490, MAB2166, D7F7 or 2D8. Similarly, reciprocal MSD assays were performed with either MW8, 1B12 or 11G2 as the capture (Fig. 7A) or detection (Fig. 7B) antibody, paired with either MAB5490, MAB2166, D7F7 or 2D8. For comparative purposes, each dataset was scaled to whichever antibody pairing had the highest signal. Signals detected in wild-type samples represented assay background, since HTT1a is not produced in wild-type mice, whereas signals detected in zQ175 samples that increased with disease pathogenesis above wild-type controls represented HTT aggregation.

On the MSD platform there was compelling evidence that N-terminal HTT fragments longer than HTT1a were increasingly incorporated into HTT aggregates throughout disease progression, when detected with either MAB5490, MAB2166, D7F7 or 2D8. Substituting MW8 with either 1B12 or 11G2 significantly improved assay signal, with 11G2 being marginally more sensitive than 1B12 (Fig. 7). Consistent with previous findings,^24^ assay signals detected by HTRF were notably weaker than by MSD (Supplementary Fig. 10 and Fig. 7). Nevertheless, on the HTRF platform, there was some evidence that N-terminal proteolytic HTT fragments longer than HTT1a were increasingly incorporated into HTT aggregates throughout disease progression, particularly when detected with MAB2166 or D7F7. Substituting MW8 with either 1B12 or 11G2 produced a modest improvement in assay signal in some assay configurations (Supplementary Fig. 10).

### PolyQ-length dependent detection of soluble and aggregated HTT1a using enhanced HTRF and MSD assays

We previously investigated the effect of CAG repeat length on mutant HTT and HTT1a protein levels.^20^ In that study, the performance of HTRF and MSD assays was compared using cortical lysates at 11 weeks of age from an allelic series of heterozygous and homozygous *Hdh*Q20, *Hdh*Q50, *Hdh*Q80, *Hdh*Q111, CAG140 and zQ175 knock-in mice, alongside age-matched wild-type littermate controls. Cortical lysates from transgenic YAC128 mice were also included to control for the detection of full-length HTT levels exceeding wild-type expression levels. Since unused hippocampal material from that study remained available, we applied the most sensitive HTT1a assays identified here to further demonstrate the effect of polyQ length on HTT bioassay performance.

To evaluate the effect of polyQ length on the detection of soluble HTT1a, we performed HTRF using the 2B7-1B12 assay (Fig. 8A) and MSD using the 2B7-11G2 assay (Fig. 8B). By HTRF, soluble HTT1a levels increased with increasing CAG repeat length (Fig. 8A), and with enhanced sensitivity, this 2B7-1B12 assay data closely mirrored our previously published 2B7-MW8 data.^20^ In contrast, by MSD, the 2B7-11G2 assay substantially improved assay performance (Fig. 8B), relative to our previously published weak and nonviable 2B7-MW8 data.^20^ Strikingly, with enhanced sensitivity, the 2B7-11G2 MSD assay data closely mimicked the 2B7-1B12 HTRF assay data, such that the detection of increasing soluble HTT1a levels with increasing CAG repeat length across the allelic series of knock-in mice was indistinguishable between both technology platforms (Fig. 8A, B). Soluble HTT1a was not detected in wild-type or N171-82Q lysates, confirming HTT1a specificity. In the hippocampus of 9-week-old YAC128 mice, soluble HTT1a levels were comparable to those observed in heterozygous *Hdh*Q80 mice at 11 weeks of age (Fig. 8A, B).

**Figure 8.**
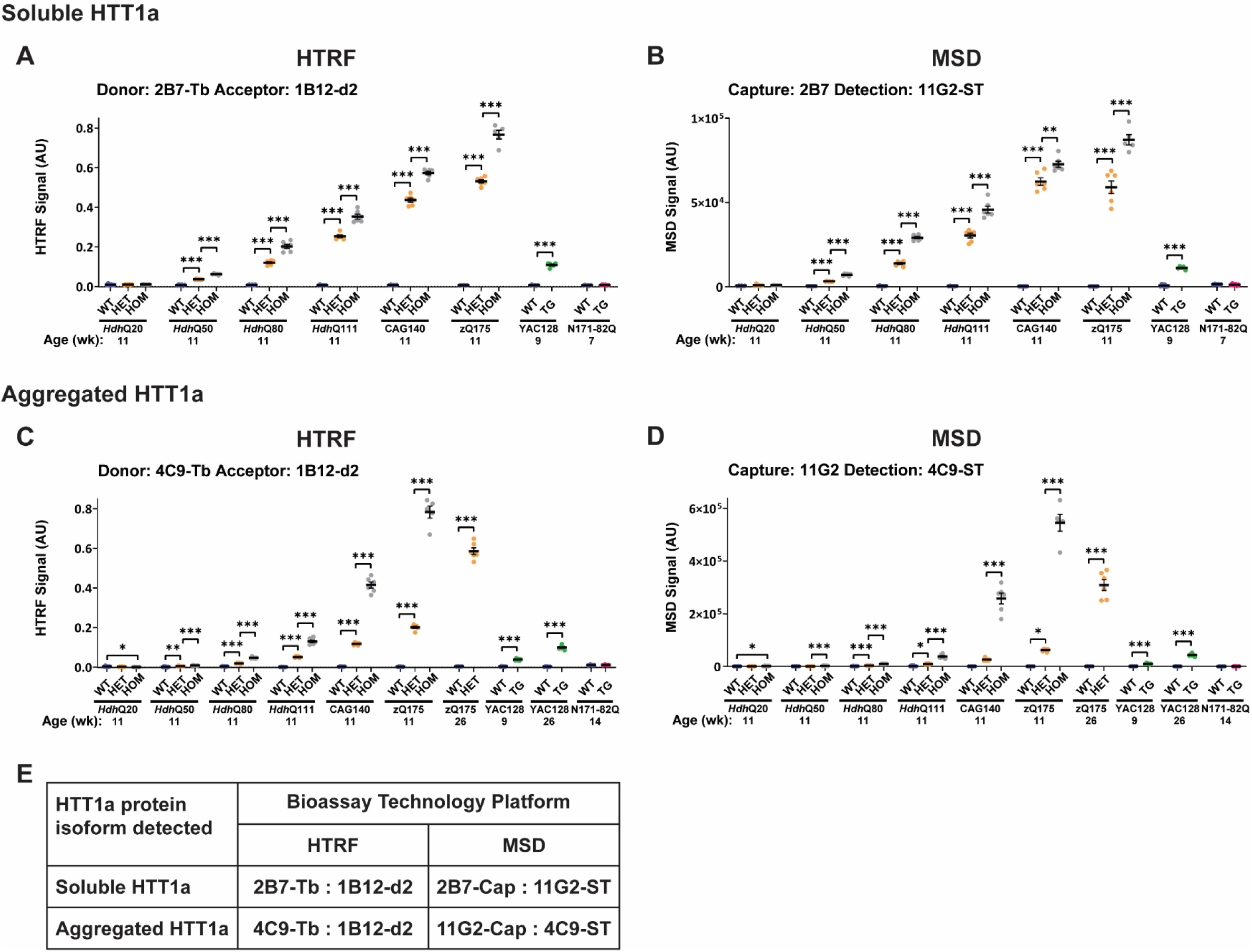
PolyQ-length dependence of HTRF and MSD assays that detect soluble and aggregated HTT1a. (A-B) For soluble HTT1a detection, 2B7-1B12 was assessed by HTRF **(A)** and 2B7-11G2 was assessed by MSD **(B)** using hippocampal lysates from heterozygous and homozygous *Hdh*Q20, *Hdh*Q50, *Hdh*Q80, *Hdh*Q111, CAG140 and zQ175 mice at 11 weeks of age, from transgenic YAC128 mice at 9 weeks of age, and from transgenic N171-82Q mice at 7 weeks of age, alongside age-matched wild-type littermate controls. **(C-D)** For aggregated HTT1a detection, 4C9-1B12 was assessed by HTRF **(C)** and 11G2-4C9 was assessed by MSD **(D)** using the same hippocampal lysates from the allelic series of knock-in mice, in addition to heterozygous zQ175 and transgenic YAC128 mice at 26 weeks of age, and from transgenic N171-82Q mice at 14 weeks of age, alongside age-matched wild-type littermate controls (*n* = 3/gender/genotype). HTT1a was not detected in transgenic N171-82Q or wild-type mice, which do not produce the HTT1a protein. **(E)** Summary of the recommended HTRF and MSD assays for detecting soluble and aggregated HTT1a. Data analysis was either by Student’s *t*-test (for YAC128 and N171-82Q) or one-way ANOVA with Bonferroni *post hoc* correction per mouse knock-in line. The test statistic, degrees of freedom and *P-*values for the ANOVA are reported in Supplementary Table 11. Error bars are mean ± SEM. \**P* ≤ 0.5, \*\**P* ≤ 0.01 and \*\*\**P* ≤ 0.001. Tb = terbium cryptate, ST = sulfo-tag, AU = arbitrary units, WT = wild type, HET = heterozygote, HOM = homozygote, TG = transgenic.

To evaluate the effect of polyQ length on the detection of aggregated HTT1a, we performed HTRF using the 4C9-1B12 assay (Fig. 8C) and MSD using the 11G2-4C9 assay (Fig. 8D) on the same hippocampal lysates. Even with these enhanced assays, and consistent with our previously published 4C9–MW8 data,^20^ the HTRF platform remained more sensitive than MSD for detecting HTT1a aggregation. At 11 weeks of age, statistically significant levels of aggregated HTT1a were first detected in heterozygous *Hdh*Q50 mice by HTRF (Fig. 8C), with aggregation levels in homozygous mice increasing more rapidly with polyQ length than in heterozygous mice (Fig. 8C, D). Aggregated HTT1a was not detected in wild-type or N171-82Q lysates, confirming HTT1a specificity. In the hippocampus of 26-week-old YAC128 mice, aggregated HTT1a levels were comparable to those observed in heterozygous CAG140 mice at 11 weeks of age (Fig. 8C, D). The recommended HTRF and MSD assays for detecting soluble and aggregated HTT1a, incorporating the most sensitive antibody pairings identified in this study, are summarized (Fig. 8E).

## DISCUSSION

Therapeutic strategies for Huntington’s disease have increasingly focused on lowering HTT protein levels by targeting the *HTT* gene or its RNA transcript. Depending on the location within the *HTT* mRNA sequence targeted by any given agent, these approaches may reduce levels of full-length HTT, HTT1a, or both protein isoforms. Recent preclinical HTT-lowering studies have demonstrated that strategies that lower HTT1a levels are substantially more efficacious than those that lower full-length HTT levels alone.^48,49^ This observation is consistent with emerging clinical trial data using Tominersen, an antisense oligonucleotide that targets full-length *HTT*, which worsened clinical endpoints in a large phase III trial.^50^ In contrast, treatment of individuals with Huntington’s disease for 36 months with AMT-130, an engineered microRNA that targets exon 1 *HTT* to lower both full-length HTT and HTT1a,^51^ has recently entered phase I/II clinical evaluation (ClinicalTrials.gov identifier: NCT04120493), and reported significant improvements in both primary and secondary clinical endpoints (https://www.uniqure.com/investors-media/press-releases).

As these therapeutic modalities advance, there is a growing need to accurately measure specific soluble and aggregated HTT protein isoforms as pharmacodynamic biomarkers in both preclinical and clinical settings. In particular, the detection of HTT1a is important, as current assays lack sufficient sensitivity in mouse models of Huntington’s disease and have not yet been developed for clinical biofluids. Using mouse models of Huntington’s disease, we have shown that 1B12 and 11G2 are *bona fide* neoepitope-specific antibodies for detecting the C-terminus of soluble and aggregated HTT1a. Furthermore, by using complementary *in vitro* (site-directed mutagenesis in COS-1 cells) and *in vivo* (immunoprecipitation, western blotting, immunohistochemistry, and HTT bioassays) experimental approaches, we have demonstrated that 1B12 and 11G2 consistently outperformed our previously published MW8-based methods for detecting HTT1a.^17,20,24,30,31,52^

For immunoprecipitation experiments coupled with western blotting in zQ175 mice, we chose the stringent detergent-rich HEPES-based buffer for lysate preparation to ensure complete solubilization of membrane-associated and detergent-resistant HTT species. Contrary to published work, ^53^ both 1B12 and 11G2 readily immunoprecipitated HTT1a with high affinity, comparable to MW8. During disease progression, detectable soluble HTT1a levels in the resolving gel decreased whilst the appearance of SDS-insoluble high-molecular-mass (HMM) HTT1a-aggregates accumulated in the stacking gel. Recently, the direct detection of HTT1a in mouse knock-in tissues by western blotting using 1B12 and 11G2 has been reported, whereby “HTT1a appeared as a band and SDS-soluble HMM smear”.^53^ In contrast, across all western blotting analyses performed here, HTT1a never resolved as a soluble HMM smear; instead, aggregated HTT1a consistently remained SDS-insoluble under standard denaturing conditions. Differences in sample preparation may account for this discrepancy, because mild sucrose-based buffers that lack detergent may only partially solubilize the highly-aggregation prone HTT1a protein.^53^ Importantly, we found that HTT1a transferred inefficiently to PVDF membranes, emphasizing the need to avoid PVDF resins when assaying HTT proteins by western blotting.

By immunohistochemical analysis of N171-82Q and zQ175 mice, we showed that 1B12 and 11G2, like MW8, function as neoepitope-specific antibodies to visualize aggregated HTT1a. Using these HTT1a-specific antibodies in zQ175 mice, we further revealed that aggregated HTT staining appears both as a diffuse nuclear signal and as distinct inclusion bodies, both of which contained HTT1a protein. This observation is consistent with atomic force microscopy analyses showing that the nanometre-scale globular structures of HTT aggregates in transgenic R6/2 mice, which express HTT1a, have dimensions identical to those observed in *Hdh*Q150 knock-in mice.^54^ For maximal sensitivity, both 1B12 and 11G2 were markedly superior to MW8 in visualizing the spatial emergence and accumulation of aggregated HTT1a by immunohistochemistry, as both antibodies readily detect the diffuse nuclear HTT1a-aggregate signal in addition to HTT inclusion pathology. This observation is consistent with a recent immunofluorescence study using MW8 and 11G2 to assess the accumulation of aggregated HTT1a in the forebrains of zQ175 mice, which also detected HTT1a inclusions in *postmortem* Huntington’s disease tissue.^55^

Developing bioassays capable of distinguishing distinct soluble and aggregated HTT isoforms is critically important. Once validated for HTT isoform specificity, their direct in-well format enables rapid, reliable and highly reproducible quantification of HTT isoforms from large numbers of tissue lysates. These attributes make such bioassays well suited for interpreting mechanistic studies and for evaluating pharmacodynamic responses in preclinical therapeutic interventions.^43,48,49,51,56,57^ Using a comprehensive evaluation of HTRF and MSD assays in zQ175 and YAC128 mouse models, we have shown that the 1B12 and 11G2 antibodies could serve as effective substitutes for, and substantially outperform, the MW8 antibody in detecting soluble and aggregated HTT1a whilst maintaining HTT1a isoform specificity. For maximal sensitivity in detecting soluble HTT1a, we recommend using the 2B7-1B12 pairing for HTRF and 2B7-11G2 pairing for MSD. Similarly, for detecting aggregated HTT1a, we recommend using the 4C9-1B12 pairing for HTRF and 11G2-4C9 pairing for MSD (Fig. 8E). The rationale for recommending these pairings was further supported by evaluating the most sensitive HTRF and MSD assays across an allelic series of knock-in mice, whereby increasing CAG length was correlated with an increase in both soluble and aggregated HTT1a levels on both technology platforms.

When pairing these more sensitive HTT1a-specific antibodies to a panel of antibodies that recognized epitopes C-terminal to HTT1a across the entire full-length HTT protein we further showed that, in zQ175 mice, N-terminal HTT fragments longer than HTT1a were increasingly being incorporated into HTT1a aggregates during disease progression. Historically, HTT inclusions have not been detected by immunohistochemistry using antibodies targeting epitopes C-terminal to HTT1a in zQ175 mouse brains,^30^ or in *postmortem* Huntington’s disease tissue.^58^ These immunoassays could be detecting aggregated HTT material that is inaccessible by conventional immunohistochemistry due to differences in epitope accessibility and detection sensitivity thresholds.

In summary, the data presented here, across multiple experimental approaches, establish 1B12 and 11G2 as robust neoepitope antibodies to reliably track HTT1a pathology *in vivo*. Both antibodies consistently outperformed our previously established MW8-based approaches,^20,24,30,31^ demonstrating superior sensitivity. Collectively, these findings support the preferential use of 1B12 or 11G2 for detecting both soluble and aggregated HTT1a, thereby strengthening the interpretation of mechanistic studies and the assessment of pharmacodynamic responses in therapeutic interventions. Whether these antibodies, when applied to more sensitive bioassay platforms,^59^ will enable the detection of HTT1a in clinical samples remains to be determined.

## ACKNOWLEDGEMENTS

We thank Brenda Lager and Britt Callahan for providing mice from the CHDI colonies at the Jackson Laboratory and David Rubinsztein and Farah Siddiqi for providing N171-82Q mice from their colony at the University of Cambridge. We acknowledge Alexandre Jean (Revvity) and Joao Nunes (MSD) for valuable technical discussions and are grateful to Kerry Chapman (Revvity) and Touran Shahbazi (MSD) for their assistance with ordering and logistical processing.

## FUNDING INFORMATION

This work was supported by grants from the CHDI Foundation.

## COMPETING INTERESTS

The authors report there are no competing interests.

## SUPPLEMENTARY MATERIAL

Supplementary material is available at *Brain* online.

## ABBREVIATIONS

aa: amino acid
AU: arbitrary units
AlphaLISA: amplified luminescent proximity homogeneous assay
BCA: bicinchoninic acid
CTX: cortex
FRET: fluorescence energy transfer
HET: heterozygous
HMM: high-molecular-mass
HOM: homozygous
HTRF: homogeneous time resolved fluorescence
HK: housekeeping
ID: immunodetect
IgG: immunoglobulin G
IP: immunoprecipitated
kDa: molecular weight in kilo Daltons
MSD: Meso Scale Discovery
N17: first 17 amino acids
PBS: phosphate buffered saline
PBST: phosphate buffered saline containing 0.1% Tween-20
PolyP: polyproline-rich domain
PolyQ: polyglutamine
Q: glutamine
PVDF: polyvinylidene difluoride
qPCR: real-time quantitative PCR
SDS: sodium dodecyl sulphate
SDS-PAGE: SDS-polyacrylamide gel electrophoresis
SD: standard deviation of the mean
SEM: standard error of the mean
ST: sulfo-tag
STR: striatum
Tb: terbium cryptate
TG: transgenic
UCL: University College London
WT: wild-type.

## Notes

### Competing Interest Statement

The authors have declared no competing interest.

## REFERENCES

1. Bates GP, Dorsey R, Gusella JF, et al. Huntington disease. Nat Rev Dis Primers. 2015;1:15005.

2. HDCRG. A novel gene containing a trinucleotide repeat that is expanded and unstable on Huntington’s disease chromosomes. The Huntington’s Disease Collaborative Research Group. Cell. 1993;72(6):971–83.

3. Duyao M, Ambrose C, Myers R, et al. Trinucleotide repeat length instability and age of onset in Huntington’s disease. Nat Genet. 1993;4(4):387–92.

4. Rubinsztein DC, Leggo J, Coles R, et al. Phenotypic characterization of individuals with 30-40 CAG repeats in the Huntington disease (HD) gene reveals HD cases with 36 repeats and apparently normal elderly individuals with 36-39 repeats. Am J Hum Genet. 1996;59(1):16–22.

5. Telenius H, Kremer HP, Theilmann J, et al. Molecular analysis of juvenile Huntington disease: the major influence on (CAG)n repeat length is the sex of the affected parent. Hum Mol Genet. 1993;2(10):1535–40.

6. Tabrizi SJ, Estevez-Fraga C, van Roon-Mom WMC, et al. Potential disease-modifying therapies for Huntington’s disease: lessons learned and future opportunities. Lancet Neurol. 2022;21(7):645–658.

7. Telenius H, Kremer B, Goldberg YP, et al. Somatic and gonadal mosaicism of the Huntington disease gene CAG repeat in brain and sperm. Nat Genet. 1994;6(4):409–14.

8. Kennedy L, Evans E, Chen CM, et al. Dramatic tissue-specific mutation length increases are an early molecular event in Huntington disease pathogenesis. Hum Mol Genet. 2003;12(24):3359–67.

9. Matlik K, Baffuto M, Kus L, et al. Cell-type-specific CAG repeat expansions and toxicity of mutant Huntingtin in human striatum and cerebellum. Nat Genet. 2024;56(3):383–394.

10. Handsaker RE, Kashin S, Reed NM, et al. Long somatic DNA-repeat expansion drives neurodegeneration in Huntington’s disease. Cell. 2025;188(3):623–639 e19.

11. Genetic Modifiers of Huntington’s Disease C. Identification of Genetic Factors that Modify Clinical Onset of Huntington’s Disease. Cell. 2015;162(3):516–26.

12. Genetic Modifiers of Huntington’s Disease Consortium. Electronic address ghmhe, Genetic Modifiers of Huntington’s Disease C. CAG Repeat Not Polyglutamine Length Determines Timing of Huntington’s Disease Onset. Cell. 2019;178(4):887–900 e14.

13. Moss DJH, Pardinas AF, Langbehn D, et al. Identification of genetic variants associated with Huntington’s disease progression: a genome-wide association study. Lancet Neurol. 2017;16(9):701–711.

14. Lee JM, Chao MJ, Harold D, et al. A modifier of Huntington’s disease onset at the MLH1 locus. Hum Mol Genet. 2017;26(19):3859–3867.

15. Shelbourne PF, Keller-McGandy C, Bi WL, et al. Triplet repeat mutation length gains correlate with cell-type specific vulnerability in Huntington disease brain. Hum Mol Genet. 2007;16(10):1133–42.

16. Pressl C, Matlik K, Kus L, et al. Selective Vulnerability of Layer 5a Corticostriatal Neurons in Huntington’s Disease. bioRxiv. 2023:2023.04.24.538096.

17. Sathasivam K, Neueder A, Gipson TA, et al. Aberrant splicing of HTT generates the pathogenic exon 1 protein in Huntington disease. Proc Natl Acad Sci U S A. 2013;110(6):2366–70.

18. Neueder A, Landles C, Ghosh R, et al. The pathogenic exon 1 HTT protein is produced by incomplete splicing in Huntington’s disease patients. Sci Rep. 2017;7(1):1307.

19. Papadopoulou AS, Gomez-Paredes C, Mason MA, Taxy BA, Howland D, Bates GP. Extensive Expression Analysis of Htt Transcripts in Brain Regions from the zQ175 HD Mouse Model Using a QuantiGene Multiplex Assay. Sci Rep. 2019;9(1):16137.

20. Landles C, Osborne GF, Phillips J, et al. Mutant huntingtin protein decreases with CAG repeat expansion: implications for therapeutics and bioassays. Brain Commun. 2024;6(6):fcae410.

21. Papadopoulou AS, Landles C, Smith EJ, et al. The HTT1a protein initiates HTT aggregation in a knock-in mouse model of Huntington’s disease. Brain. 2026.

22. Weiss A, Abramowski D, Bibel M, et al. Single-step detection of mutant huntingtin in animal and human tissues: a bioassay for Huntington’s disease. Anal Biochem. 2009;395(1):8–15.

23. Baldo B, Paganetti P, Grueninger S, et al. TR-FRET-based duplex immunoassay reveals an inverse correlation of soluble and aggregated mutant huntingtin in huntington’s disease. Chem Biol. 2012;19(2):264–75.

24. Landles C, Milton RE, Jean A, et al. Development of novel bioassays to detect soluble and aggregated Huntingtin proteins on three technology platforms. Brain Commun. 2021;3(1):fcaa231.

25. Macdonald D, Tessari MA, Boogaard I, et al. Quantification assays for total and polyglutamine-expanded huntingtin proteins. PLoS One. 2014;9(5):e96854.

26. Reindl W, Baldo B, Schulz J, et al. Meso scale discovery-based assays for the detection of aggregated huntingtin. PLoS One. 2019;14(3):e0213521.

27. Baldo B, Sajjad MU, Cheong RY, et al. Quantification of Total and Mutant Huntingtin Protein Levels in Biospecimens Using a Novel alphaLISA Assay. eNeuro. 2018;5(4).

28. Fodale V, Boggio R, Daldin M, et al. Validation of Ultrasensitive Mutant Huntingtin Detection in Human Cerebrospinal Fluid by Single Molecule Counting Immunoassay. J Huntingtons Dis. 2017;6(4):349–361.

29. Wild EJ, Boggio R, Langbehn D, et al. Quantification of mutant huntingtin protein in cerebrospinal fluid from Huntington’s disease patients. J Clin Invest. 2015;125(5):1979–86.

30. Smith EJ, Sathasivam K, Landles C, et al. Early detection of exon 1 huntingtin aggregation in zQ175 brains by molecular and histological approaches. Brain Commun. 2023;5(1):fcad010.

31. Landles C, Sathasivam K, Weiss A, et al. Proteolysis of mutant huntingtin produces an exon 1 fragment that accumulates as an aggregated protein in neuronal nuclei in Huntington disease. J Biol Chem. 2010;285(12):8808–23.

32. Baldo B, Peladan J, Albers J, et al. Development of an HTT exon1-selective MSD immunoassay with novel HTT P90 neo-epitope-specific antibodies. J Neurol Neurosurg Psychiat. 2024;95(Suppl 1)(D: Wet biomarkers):A32.

33. Wheeler VC, Auerbach W, White JK, et al. Length-dependent gametic CAG repeat instability in the Huntington’s disease knock-in mouse. Hum Mol Genet. 1999;8(1):115–22.

34. Menalled LB, Kudwa AE, Miller S, et al. Comprehensive behavioral and molecular characterization of a new knock-in mouse model of Huntington’s disease: zQ175. PLoS One. 2012;7(12):e49838.

35. Heikkinen T, Lehtimaki K, Vartiainen N, et al. Characterization of neurophysiological and behavioral changes, MRI brain volumetry and 1H MRS in zQ175 knock-in mouse model of Huntington’s disease. PLoS One. 2012;7(12):e50717.

36. Fienko S, Landles C, Sathasivam K, et al. Alternative processing of human HTT mRNA with implications for Huntington’s disease therapeutics. Brain. 2022;145(12):4409–4424.

37. Mason MA, Gomez-Paredes C, Sathasivam K, Neueder A, Papadopoulou AS, Bates GP. Silencing Srsf6 does not modulate incomplete splicing of the huntingtin gene in Huntington’s disease models. Sci Rep. 2020;10(1):14057.

38. Pouladi MA, Stanek LM, Xie Y, et al. Marked differences in neurochemistry and aggregates despite similar behavioural and neuropathological features of Huntington disease in the full-length BACHD and YAC128 mice. Hum Mol Genet. 2012;21(10):2219–32.

39. Schilling G, Becher MW, Sharp AH, et al. Intranuclear inclusions and neuritic aggregates in transgenic mice expressing a mutant N-terminal fragment of huntingtin. Hum Mol Genet. 1999;8(3):397–407.

40. Landles C, Milton RE, Ali N, et al. Subcellular Localization And Formation Of Huntingtin Aggregates Correlates With Symptom Onset And Progression In A Huntington’S Disease Model. Brain Commun. 2020;2(2):fcaa066.

41. Fienko S, Landles C, Sathasivam K, et al. Alternative processing of human HTT mRNA with implications for Huntington’s disease therapeutics. Brain. 2022; 145(12):4409–4424.

42. Livak KJ, Schmittgen TD. Analysis of relative gene expression data using real-time quantitative PCR and the 2(-Delta Delta C(T)) Method. Methods. 2001;25(4):402–408.

43. Aldous SG, Smith EJ, Landles C, et al. A CAG repeat threshold for therapeutics targeting somatic instability in Huntington’s disease. Brain. 2024;147(5):1784–1798.

44. Ko J, Ou S, Patterson PH. New anti-huntingtin monoclonal antibodies: implications for huntingtin conformation and its binding proteins. Brain Res Bull. 2001;56(3-4):319–29.

45. Menalled LB, Sison JD, Dragatsis I, Zeitlin S, Chesselet MF. Time course of early motor and neuropathological anomalies in a knock-in mouse model of Huntington’s disease with 140 CAG repeats. J Comp Neurol. 2003;465(1):11–26.

46. Schilling G, Wood JD, Duan K, et al. Nuclear accumulation of truncated atrophin-1 fragments in a transgenic mouse model of DRPLA. Neuron. 1999;24(1):275–86.

47. Slow EJ, van Raamsdonk J, Rogers D, et al. Selective striatal neuronal loss in a YAC128 mouse model of Huntington disease. Hum Mol Genet. 2003;12(13):1555–67.

48. Bragg RM, Landles C, Smith EJ, et al. Selective targeting of mutant *huntingtin* intron-1 improves rescue provided by antisense oligonucleotides. bioRxiv. 2025:2025.07.29.665998.

49. Papadopoulou AS, Alterman J, Landles C, et al. Lowering the *HTT1a* transcript as an effective therapy for Huntington’s disease. bioRxiv. 2025:2025.06.10.658804.

50. Kingwell K. Double setback for ASO trials in Huntington disease. Nat Rev Drug Discov. 2021;20(6):412–413.

51. Sogorb-Gonzalez M, Landles C, Caron NS, et al. Exon 1-targeting miRNA reduces the pathogenic exon 1 HTT protein in Huntington’s disease models. Brain. 2024;147(12):4043–4055.

52. Franich NR, Hickey MA, Zhu C, et al. Phenotype onset in Huntington’s disease knock-in mice is correlated with the incomplete splicing of the mutant huntingtin gene. J Neurosci Res. 2019;97(12):1590–1605.

53. Sapp E, Boudi A, Iwanowicz A, et al. Mutant huntingtin exon 1 protein detected in mouse brain with neoepitope antibody: effects of CAG repeat expansion, MutS Homolog 3 silencing and aggregation. Brain Commun. 2025;7(5):fcaf314.

54. Sathasivam K, Lane A, Legleiter J, et al. Identical oligomeric and fibrillar structures captured from the brains of R6/2 and knock-in mouse models of Huntington’s disease. Hum Mol Genet. 2010;19(1):65–78.

55. Deng Y, Joni M, Wang H, et al. Localization of mutant huntingtin with HTT Exon1 P90 C-terminal neoepitope antibodies in relation to regional and neuronal vulnerability in forebrain in Q175 mice and human huntington’s disease. J Huntingtons Dis. 2025:18796397251404999.

56. Murillo A, Alpaugh M, Peter Durairaj RR, et al. Cas9 Nickase-Mediated Contraction of CAG/CTG Repeats *in Vivo* is Accompanied by Improvements in Huntington’s Disease Pathology. bioRxiv. 2025:2024.02.19.580669.

57. Belgrad J, Summers A, Landles C, et al. Blocking somatic repeat expansion and lowering huntingtin via RNA interference synergize to prevent Huntington’s disease pathogenesis in mice. bioRxiv. 2025:2025.06.24.661398.

58. Schilling G, Klevytska A, Tebbenkamp AT, et al. Characterization of huntingtin pathologic fragments in human Huntington disease, transgenic mice, and cell models. J Neuropathol Exp Neurol. 2007;66(4):313–20.

59. Feng W, Beer J, Hao Q, et al. NULISA: a novel proteomic liquid biopsy platform with attomolar sensitivity and high multiplexing. bioRxiv. 2023.

