## Supplementary material for "Enhancing the detection of HTT1a with neoepitope antibodies in mouse models of Huntington’s disease"

### Supplementary Table 1. Summary of the antibodies used in this study.

| Name | Immunogen | Epitope | Species | Source |
|---|---|---|---|---|
| **2B7** | Human HTT peptide, amino acids: 1-17.[1] | Within amino acids: 7-13 LMKAFE*. | Mouse, monoclonal | CHDI Foundation |
| **MW1** | Human DRPLA (19Q) fusion protein. | Polyglutamine (PolyQ).[2] | Mouse, monoclonal | CHDI Foundation |
| **3B5H10** | HTT fusion protein, amino acids: 1-171 (66Q).[3] | Polyglutamine (PolyQ). | Mouse, monoclonal | Sigma-Aldrich, P1874 |
| **4C9** | Human HTT peptide, amino acids: 51-71.[4] | Human polyproline-rich domain, (does not detect mouse). | Mouse, monoclonal | CHDI Foundation |
| **S830** | Human HTT exon 1 fusion protein (53Q).[5] | | Sheep, polyclonal | In-house |
| **MW8** | HTT exon 1 fusion protein, amino acids 1-90 (67Q).[2] | Within amino acids: 83-90 AEEPLHRP, ends at proline.[6] | Mouse, monoclonal | Merck MABN2529 |
| **1B12** | Human HTT peptide, C-AEEPLHRP(OH). | Terminates precisely at amino acids: 83-90, AEEPLHRP(OH). | Rabbit, monoclonal | CHDI Foundation |
| **11G2** | Human HTT peptide, C-AEEPLHRP(OH). | Terminates precisely at amino acids: 83-90, AEEPLHRP(OH). | Rabbit, monoclonal | CHDI Foundation |
| **MAB5490** | Human HTT peptide, amino acids: 115-129.[7] | Within amino acids: 115-129 QSVRNSPEFQKLLGI (mouse L). | Mouse, monoclonal | Sigma-Aldrich, MAB5490 |
| **MAB2166** | HTT fusion protein, amino acids: 181-810.[8] | Within amino acids: 443-457, GKVLLGEEEALEDDS.[9] | Mouse, monoclonal | Sigma-Aldrich, MAB2166 |
| **D7F7** | Human HTT peptide. | Within amino acids: 1214-1223, QSDTSGPVTT* (mouse A). | Rabbit, monoclonal | Cell Signaling Technology, #5656 |
| **2D8** | Human HTT peptide, amino acids: 3093-3115. | Within amino acids: 3093-3115, LVATDFYRHQIEEELDRRAFQSV (mouse F). | Rabbit, monoclonal | CHDI Foundation |

*Information provided by CHDI Foundation, DRPLA = Dentatorubral-pallidoluysian atrophy.

### References:


1. Weiss A, Abramowski D, Bibel M, *et al*. Single-step detection of mutant huntingtin in animal and human tissues: a bioassay for Huntington's disease. *Anal Biochem*. 2009;395(1):8-15.

2. Ko J, Ou S, Patterson PH. New anti-huntingtin monoclonal antibodies: implications for huntingtin conformation and its binding proteins. *Brain research bulletin*. 2001;56(3-4):319-329.

3. Peters-Libeu C, Newhouse Y, Krishnan P, *et al*. Crystallization and diffraction properties of the Fab fragment of 3B5H10, an antibody specific for disease-causing polyglutamine stretches. *Acta Crystallogr Sect F Struct Biol Cryst Commun*. 2005;61(Pt 12):1065-8.

4. Baldo B, Paganetti P, Grueninger S, *et al*. TR-FRET-based duplex immunoassay reveals an inverse correlation of soluble and aggregated mutant huntingtin in huntington's disease. *Chemistry & biology*. 2012;19(2):264-275.

5. Sathasivam K, Woodman B, Mahal A, *et al*. Centrosome disorganization in fibroblast cultures derived from R6/2 Huntington's disease (HD) transgenic mice and HD patients. *Hum Mol Genet*. 2001;10(21):2425-35.

6. Landles C, Sathasivam K, Weiss A, *et al*. Proteolysis of mutant huntingtin produces an exon 1 fragment that accumulates as an aggregated protein in neuronal nuclei in Huntington disease. *J Biol Chem*. 2010;285(12):8808-8823.

7. Lunkes A, Lindenberg KS, Ben-Haiem L, *et al*. Proteases acting on mutant huntingtin generate cleaved products that differentially build up cytoplasmic and nuclear inclusions. *Molecular cell*. 2002;10(2):259-269.

8. Trottier Y, Devys D, Imbert G, *et al*. Cellular localization of the Huntington's disease protein and discrimination of the normal and mutated form [see comments]. *Nature genetics*. 1995;10(1):104-110.

9. Cong SY, Pepers BA, Roos RA, Van Ommen GJ, Dorsman JC. Epitope mapping of monoclonal antibody 4C8 recognizing the protein huntingtin. *Hybridoma (Larchmt)*. 2005;24(5):231-5.

### Supplementary Table 2. Antibody and lysate concentrations used for HTRF assays.

| Figure number | Antibody pairing | Donor (ng / well) | Acceptor (ng / well) | Lysate dilution concentration |
|---|---|---|---|---|
| Figure 2: zQ175 & N171-82Q Supplementary Figure 5: YAC128 & N171-82Q | 2B7-Tb : MW1-d2 | 1 ng | 20 ng | 2.5% |
| | 2B7-Tb : MW8-d2 | 1 ng | 40 ng | 10% |
| | 2B7-Tb : 1B12-d2 | 1 ng | 20 ng | 10% |
| | 2B7-Tb : 11G2-d2 | 1 ng | 20 ng | 10% |
| Figure 3: zQ175 & N171-82Q Supplementary Figure 6: YAC128 & N171-82Q | 4C9-Tb : S830-d2 | 1 ng | 40 ng | 10% Crude Lysate |
| | MW8-Tb : 2B7-d2 | 1 ng | 20 ng | 10% Crude Lysate |
| | 1B12-Tb : 2B7-d2 | 1 ng | 20 ng | 10% |
| | 11G2-Tb : 2B7-d2 | 1 ng | 20 ng | 10% |
| Figure 6: zQ175 & N171-82Q Supplementary Figure 8: YAC128 & N171-82Q | 4C9-Tb : MW8-d2 | 1 ng | 20 ng | 10% Crude Lysate |
| | 4C9-Tb : 1B12-d2 | 1 ng | 20 ng | 10% Crude Lysate |
| | 4C9-Tb : 11G2-d2 | 1 ng | 20 ng | 10% Crude Lysate |
| | MW8-Tb : 4C9-Alexa488 | 1 ng | 20 ng | 10% Crude Lysate |
| | 1B12-Tb : 4C9-Alexa488 | 1 ng | 20 ng | 10% Crude Lysate |
| | 11G2-Tb : 4C9-Alexa488 | 1 ng | 20 ng | 10% Crude Lysate |
| Supplementary Figure 9: *Hdh*Q20, zQ175 & N171-82Q | D7F7-Tb : 4C9-Alexa488 | 1 ng | 20 ng | 10% |
| | 4C9-Tb : MW8-d2 | 1 ng | 20 ng | 10% Crude Lysate |
| | 4C9-Tb : 1B12-d2 | 1 ng | 20 ng | 10% Crude Lysate |
| | 4C9-Tb : 11G2-d2 | 1 ng | 20 ng | 10% Crude Lysate |
| | MW8-Tb : 4C9-Alexa488 | 1 ng | 20 ng | 10% Crude Lysate |
| | 1B12-Tb : 4C9-Alexa488 | 1 ng | 20 ng | 10% Crude Lysate |
| | 11G2-Tb : 4C9-Alexa488 | 1 ng | 20 ng | 10% Crude Lysate |
| Supplementary Figure 10: zQ175 | MW8-Tb : MAB5490-d2 | 1 ng | 20 ng | 10% Crude Lysate |
| | 1B12-Tb : MAB5490-d2 | 1 ng | 20 ng | 10% Crude Lysate |
| | 11G2-Tb : MAB5490-d2 | 1 ng | 20 ng | 10% Crude Lysate |
| | MW8-Tb : MAB2166-d2 | 1 ng | 20 ng | 10% Crude Lysate |
| | 1B12-Tb : MAB2166-d2 | 1 ng | 20 ng | 10% Crude Lysate |
| | 11G2-Tb : MAB2166-d2 | 1 ng | 20 ng | 10% Crude Lysate |
| | MW8-Tb : D7F7-d2 | 1 ng | 20 ng | 10% Crude Lysate |
| | 1B12-Tb : D7F7-d2 | 1 ng | 20 ng | 10% Crude Lysate |
| | 11G2-Tb : D7F7-d2 | 1 ng | 20 ng | 10% Crude Lysate |
| | MW8-Tb : 2D8-d2 | 1 ng | 20 ng | 10% Crude Lysate |
| | 1B12-Tb : 2D8-d2 | 1 ng | 20 ng | 10% Crude Lysate |
| | 11G2-Tb : 2D8-d2 | 1 ng | 20 ng | 10% Crude Lysate |
| | MAB5490-Tb : MW8-d2 | 1 ng | 20 ng | 10% Crude Lysate |
| | MAB5490-Tb : 1B12-d2 | 1 ng | 20 ng | 10% Crude Lysate |
| | MAB5490-Tb: 11G2-d2 | 1 ng | 20 ng | 10% Crude Lysate |
| | MAB2166-Tb : MW8-d2 | 1 ng | 20 ng | 10% Crude Lysate |
| | MAB2166-Tb : 1B12-d2 | 1 ng | 20 ng | 10% Crude Lysate |
| | MAB2166-Tb : 11G2-d2 | 1 ng | 20 ng | 10% Crude Lysate |
| | D7F7-Tb : MW8-d2 | 1 ng | 20 ng | 10% Crude Lysate |
| | D7F7-Tb : 1B12-d2 | 1 ng | 20 ng | 10% Crude Lysate |
| | D7F7-Tb : 11G2-d2 | 1 ng | 20 ng | 10% Crude Lysate |
| | 2D8-Tb : MW8-d2 | 1 ng | 20 ng | 10% Crude Lysate |
| | 2D8-Tb : 1B12-d2 | 1 ng | 20 ng | 10% Crude Lysate |
| | 2D8-Tb : 11G2-d2 | 1 ng | 20 ng | 10% Crude Lysate |
| Figure 8 Knock-in allelic series | 2B7-Tb : 1B12-d2 | 1 ng | 20 ng | 10% |
| | 4C9-Tb : 1B12-d2 | 1 ng | 20 ng | 10% Crude Lysate |

NB: Final antibody and lysate dilution conditions must be optimized by the user, as instrument sensitivity may differ depending on the plate reader make/model. Tb = terbium cryptate.

**Supplementary Table 3. Antibody and lysate concentrations used for MSD assays.**

| Figure number | Antibody pairing | Capture | Sulfo-tag (μg/mL) | Lysate dilution concentration |
|---|---|---|---|---|
| Figure 2: zQ175 & N171-82Q Supplementary Figure 5: YAC128 & N171-82Q | 2B7-Capture : MW1-ST | MSD default | 1.5 μg/mL | 12% |
| | 2B7-Capture : MW8-ST | MSD default | 2.5 μg/mL | 18% |
| | 2B7-Capture : 1B12-ST | MSD default | 2.5 μg/mL | 18% |
| | 2B7-Capture : 11G2-ST | MSD default | 2.5 μg/mL | 18% |
| Figure 3: zQ175 & N171-82Q Supplementary Figure 6: YAC128 & N171-82Q | MW8-Capture : 2B7-ST | MSD default | 2.5 μg/mL | 18% Crude Lysate |
| | 1B12-Capture : 2B7-ST | MSD default | 2.5 μg/mL | 18% Crude Lysate |
| | 11G2-Capture : 2B7-ST | MSD default | 2.5 μg/mL | 18% Crude Lysate |
| Figure 6: zQ175 & N171-82Q Supplementary Figure 8: YAC128 & N171-82Q | 4C9-Capture : MW8-ST | MSD default | 2.5 μg/mL | 18% Crude Lysate |
| | 4C9-Capture : 1B12-ST | MSD default | 2.5 μg/mL | 18% Crude Lysate |
| | 4C9-Capture : 11G2-ST | MSD default | 2.5 μg/mL | 18% Crude Lysate |
| | MW8-Capture : 4C9-ST | MSD default | 2.5 μg/mL | 18% Crude Lysate |
| | 1B12-Capture : 4C9-ST | MSD default | 2.5 μg/mL | 18% Crude Lysate |
| | 11G2-Capture : 4C9-ST | MSD default | 2.5 μg/mL | 18% Crude Lysate |
| Supplementary Figure 9: *Hdh*Q20, zQ175 & N171-82Q | D7F7-Capture : 4C9-ST | MSD default | 2.5 μg/mL | 18% |
| | 4C9-Capture : MW8-ST | MSD default | 2.5 μg/mL | 18% Crude Lysate |
| | 4C9-Capture : 1B12-ST | MSD default | 2.5 μg/mL | 18% Crude Lysate |
| | 4C9-Capture : 11G2-ST | MSD default | 2.5 μg/mL | 18% Crude Lysate |
| | MW8-Capture : 4C9-ST | MSD default | 2.5 μg/mL | 18% Crude Lysate |
| | 1B12-Capture : 4C9-ST | MSD default | 2.5 μg/mL | 18% Crude Lysate |
| | 11G2-Capture : 4C9-ST | MSD default | 2.5 μg/mL | 18% Crude Lysate |
| Figure 7: zQ175 | MW8-Capture : MAB5490-ST | MSD default | 2.5 μg/mL | 18% Crude Lysate |
| | 1B12-Capture : MAB5490-ST | MSD default | 2.5 μg/mL | 18% Crude Lysate |
| | 11G2-Capture : MAB5490-ST | MSD default | 2.5 μg/mL | 18% Crude Lysate |
| | MW8-Capture : MAB2166-ST | MSD default | 2.5 μg/mL | 18% Crude Lysate |
| | 1B12-Capture : MAB2166-ST | MSD default | 2.5 μg/mL | 18% Crude Lysate |
| | 11G2-Capture : MAB2166-ST | MSD default | 2.5 μg/mL | 18% Crude Lysate |
| | MW8-Capture : D7F7-ST | MSD default | 2.5 μg/mL | 18% Crude Lysate |
| | 1B12-Capture : D7F7-ST | MSD default | 2.5 μg/mL | 18% Crude Lysate |
| | 11G2-Capture : D7F7-ST | MSD default | 2.5 μg/mL | 18% Crude Lysate |
| | MW8-Capture : 2D8-ST | MSD default | 2.5 μg/mL | 18% Crude Lysate |
| | 1B12-Capture : 2D8-ST | MSD default | 2.5 μg/mL | 18% Crude Lysate |
| | 11G2-Capture : 2D8-ST | MSD default | 2.5 μg/mL | 18% Crude Lysate |
| | MAB5490-Capture : MW8-ST | MSD default | 2.5 μg/mL | 18% Crude Lysate |
| | MAB5490-Capture : 1B12-ST | MSD default | 2.5 μg/mL | 18% Crude Lysate |
| | MAB5490-Capture : 11G2-ST | MSD default | 2.5 μg/mL | 18% Crude Lysate |
| | MAB2166-Capture : MW8-ST | MSD default | 2.5 μg/mL | 18% Crude Lysate |
| | MAB2166-Capture : 1B12-ST | MSD default | 2.5 μg/mL | 18% Crude Lysate |
| | MAB2166-Capture : 11G2-ST | MSD default | 2.5 μg/mL | 18% Crude Lysate |
| | D7F7-Capture : MW8-ST | MSD default | 2.5 μg/mL | 18% Crude Lysate |
| | D7F7-Capture : 1B12-ST | MSD default | 2.5 μg/mL | 18% Crude Lysate |
| | D7F7-Capture : 11G2-ST | MSD default | 2.5 μg/mL | 18% Crude Lysate |
| | 2D8-Capture : MW8-ST | MSD default | 2.5 μg/mL | 18% Crude Lysate |
| | 2D8-Capture : 1B12-ST | MSD default | 2.5 μg/mL | 18% Crude Lysate |
| | 2D8-Capture : 11G2-ST | MSD default | 2.5 μg/mL | 18% Crude Lysate |
| Figure 8: Knock-in allelic series | 2B7-Capture : 11G2-ST | MSD default | 2.5 μg/mL | 18% |
| | 11G2-Capture : 4C9-ST | MSD default | 2.5 μg/mL | 18% Crude Lysate |

MSD = Meso Scale Discovery, ST = sulfo-tag.

### Supplementary Figure 1.

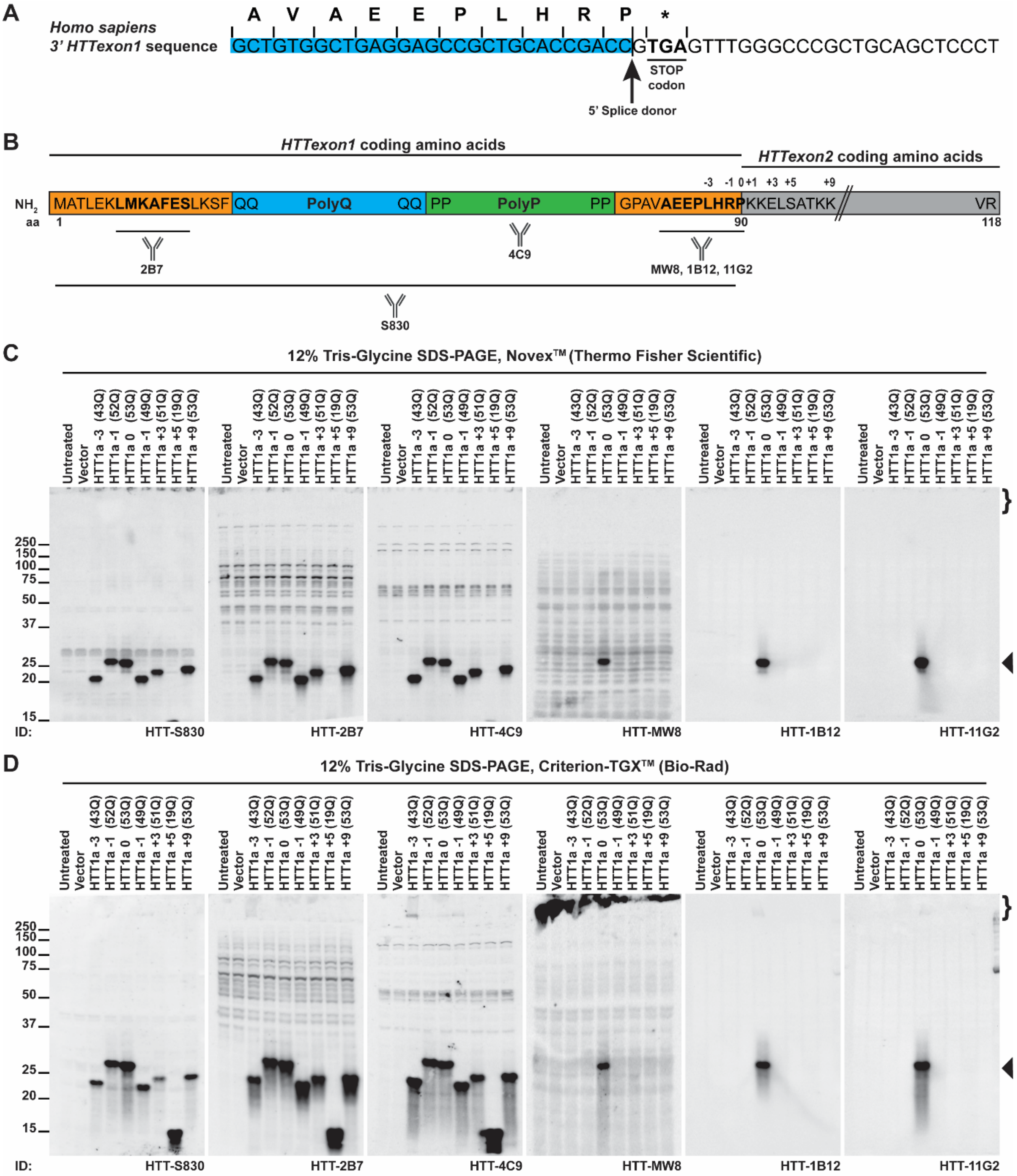

**Supplementary Figure 1. MW8, 1B12 and 11G2 are HTT1a-specific neoepitope antibodies by western blotting. (A)** Schematic illustration of the human 3′ *HTT* exon1 (blue shaded) and intron1 splice junction DNA sequence. The amino acid codon usage is indicated above the DNA sequence, with the 5′ splice donor site and STOP codon (*) highlighted. HTT1a terminates at a C-terminal proline residue. **(B)** Schematic representation of the locations of the HTT epitopes recognized by the antibodies that bind to HTT1a used in this study. HTT1a is 90 amino acids long (with 21 CAGs), terminates at a C-terminal proline residue, and does not contain any additional amino acids that are not present in full-length HTT. The numbers above the amino acid sequence indicate positions at which the pSG5-HTT1a construct was sequentially modified by the deletion or addition of amino acids relative to the C-terminal HTT proline residue (-3 to +9). The antigens used to generate 4C9 and S830, as well as the epitopes recognized by 2B7, MW8, 1B12 and 11G2 in bold, are shown. Details of all immunogens and antibody epitopes are provided in Supplementary Table 1. **(C-D)** Mutant pSG5-HTT1a constructs (-3 to +9) were transiently expressed in COS-1 cells for 48 h, after which lysates were prepared in HEPES buffer and resolved by either **(C)** 12% Novex™ (Thermo Fisher Scientific) or **(D)** 12% Criterion-TGX™ (Bio-Rad) Tris-Glycine SDS-polyacrylamide gel electrophoresis (SDS-PAGE). Proteins were transferred onto nitrocellulose membranes, and probed with either S830, 2B7, 4C9, MW8, 1B12 or 11G2 antibodies in phosphate buffered saline (PBS) containing 0.1% Tween-20 (PBST). When probed with either S830, 2B7, or 4C9, all mutant HTT constructs were detected. In contrast, when probed with either MW8, 1B12 or 11G2, only the HTT1a protein that terminated at the native C-terminal proline residue was detected (◀, closed arrow), demonstrating that these antibodies function as neoepitope antibodies specific for soluble HTT1a. aa = amino acid, PolyQ = polyglutamine tract, PolyP = polyproline-rich domain, } = stacking gel, ▶ = HTT1a, ID = immunodetect. Protein size markers are indicated in kDa.

**Supplementary Figure 2.**

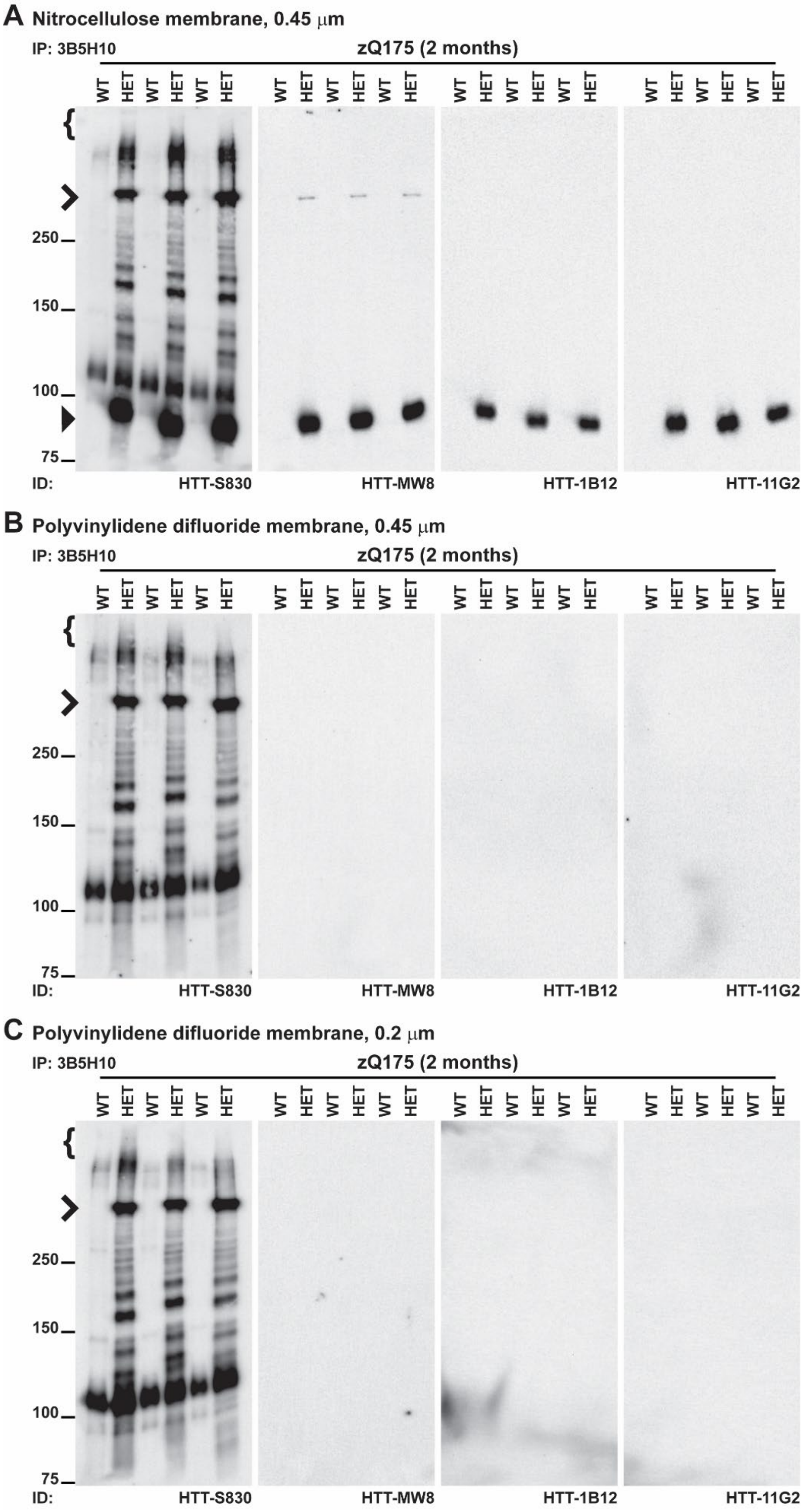

**Supplementary Figure 2. Comparison of the immunoblotting efficiency for immunoprecipitated mutant HTT using different membrane resins by western blotting. (A-C)** Mutant HTT was immunoprecipitated from cortical lysates of zQ175 mice at 2 months of age using the 3B5H10 antibody that detects expanded polyQ repeats. Lysates were prepared and immunoprecipitated in HEPES buffer, resolved by 8% Novex™ (Thermo Fisher Scientific) Tris-Glycine SDS-PAGE, and transferred onto either **(A)** Nitrocellulose (0.45 mm), **(B)** Polyvinylidene difluoride (PVDF) (0.45 mm), or **(C)** PVDF (0.2 mm) membranes (Bio-Rad). Blots were cut into strips and probed with either S830, MW8, 1B12 or 11G2 antibodies in PBST (n = 3/genotype). When zQ175 samples were transferred onto a nitrocellulose membrane and probed with S830, a characteristic pattern of full-length mutant HTT (**>,** open arrow), N-terminally cleaved mutant HTT fragments, and soluble HTT1a (▶, closed arrow) was observed. However, when transferred onto PVDF membranes (0.45 µm or 0.2 µm), whilst full-length mutant HTT and N-terminally cleaved mutant HTT fragments remained detected, soluble HTT1a was absent. Similarly, when probed with either MW8, 1B12 or 11G2, soluble HTT1a was only detected on nitrocellulose membranes and absent on both PVDF membrane types. IP = immunoprecipitate, WT = wild type, HET = heterozygote, } = stacking gel, **>** = full-length mutant HTT, ▶ = soluble HTT1a, ID = immunodetect. Protein size markers are indicated in kDa.

**Supplementary Figure 3.**

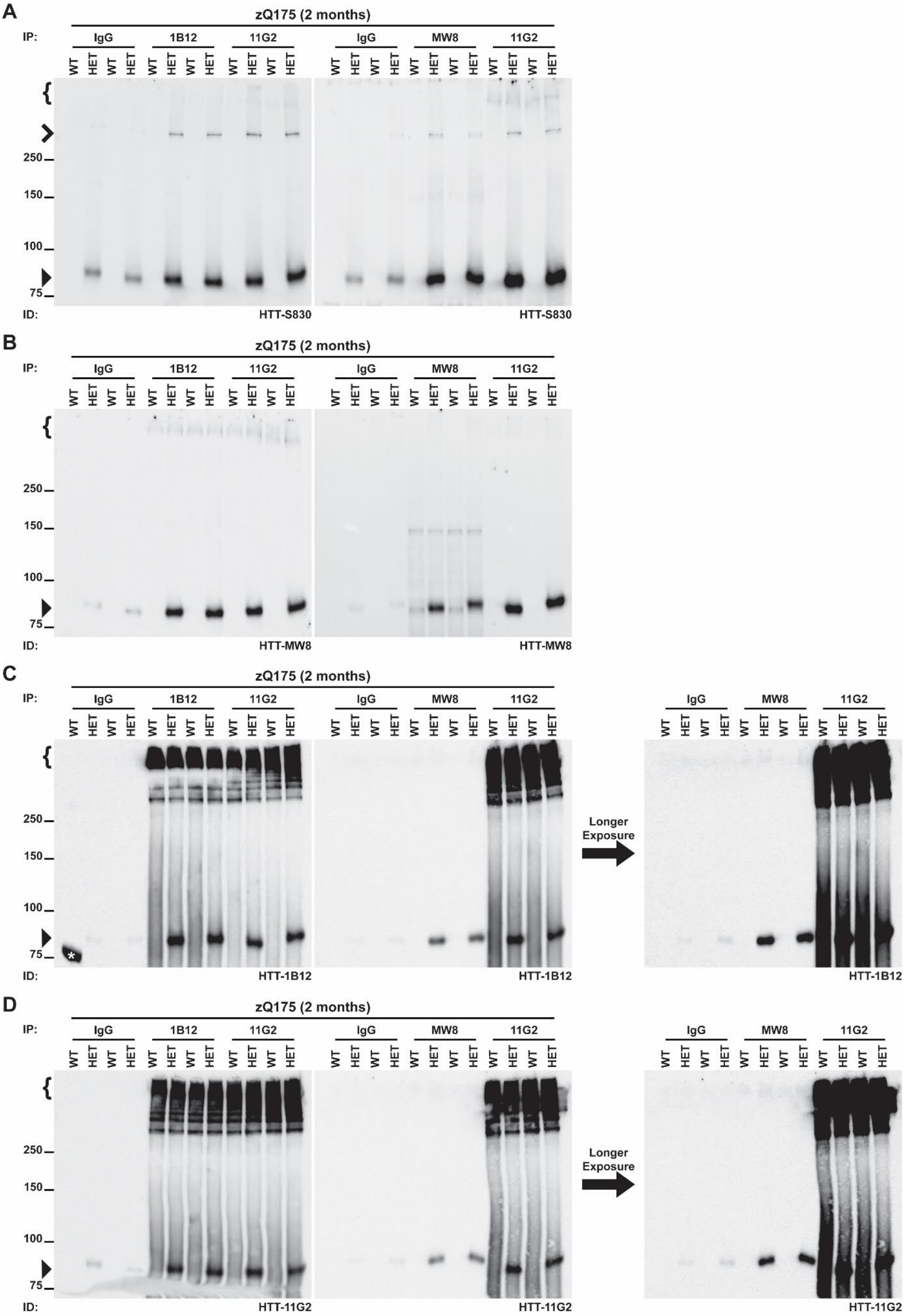

**Supplementary Figure 3. Comparison of the immunoblotting efficiency for reciprocally immunoprecipitated HTT1a by western blotting. (A-D)** HTT1a was immunoprecipitated from cortical lysates of zQ175 mice at 2 months of age using either MW8, 1B12 or 11G2 neoepitope antibodies that specifically recognizes the mutant HTT1a protein isoform. Lysates were prepared and immunoprecipitated in HEPES buffer, resolved by 8% Novex™ (Thermo Fisher Scientific) Tris-Glycine SDS-PAGE, and transferred onto nitrocellulose membranes. Mouse immunoglobulin G (IgG) served as a negative background control. Blots were probed with either **(A)** S830, **(B)** MW8, **(C)** 1B12, or **(D)** 11G2 antibodies in PBST (n = 2/genotype). HTT1a was not detected in wild-type mice that do not produce the HTT1a protein. In zQ175 samples, relative to the IgG control, all three HTT1a neoepitope antibodies successfully immunoprecipitated soluble HTT1a (▶, closed arrow). When probed with S830 a faint full-length mutant HTT (**>,** open arrow) signal was detected above the IgG control. The most effective conditions for immunoprecipitating and detecting HTT1a with minimal nonspecific background was achieved when reciprocal species-specific (mouse ↔ rabbit) antibody combinations were used. Specifically, to either immunoprecipitate with mouse MW8 followed by probing with either rabbit 1B12 or 11G2, or immunoprecipitate with either rabbit 1B12 or 11G2 followed by probing with mouse MW8. IP = immunoprecipitate, IgG = immunoglobulin G, WT = wild type, HET = heterozygote, } = stacking gel, **>** = full-length mutant HTT, ▶ = soluble HTT1a, ID = immunodetect, ★ = nonspecific protein size marker band co-loaded in lane 1. Protein size markers are indicated in kDa.

**Supplementary Figure 4.**

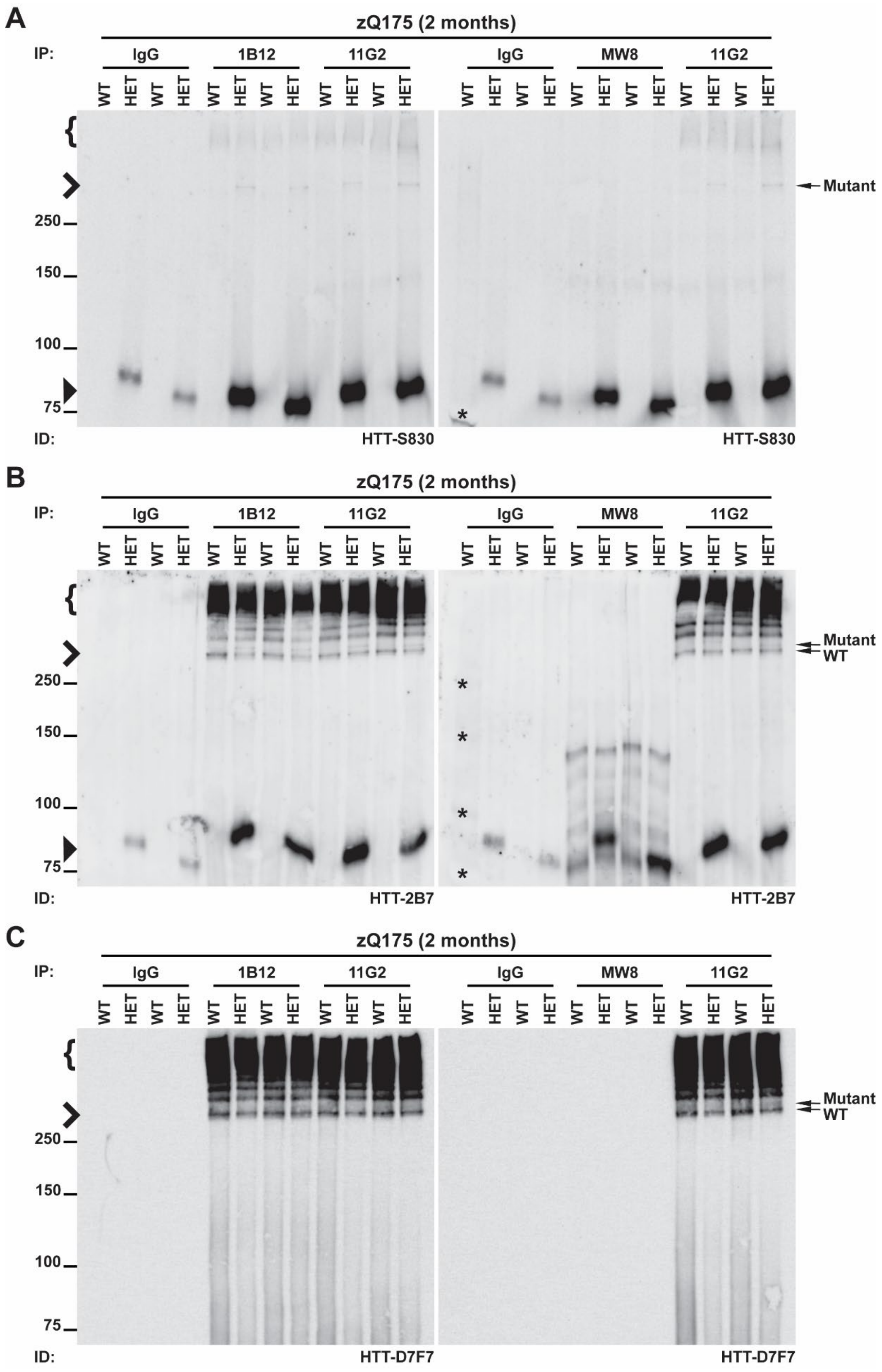

**Supplementary Figure 4. A small amount of full-length HTT (wild-type and mutant) is detected when HTT1a is immunoprecipitated by western blotting. (A-C)** HTT1a was immunoprecipitated from cortical lysates of zQ175 mice at 2 months of age using either MW8, 1B12 or 11G2 neoepitope antibodies that specifically recognizes the mutant HTT1a protein isoform. Lysates were prepared and immunoprecipitated in HEPES buffer, resolved by 8% Novex™ (Thermo Fisher Scientific) Tris-Glycine SDS-PAGE, and transferred onto nitrocellulose membranes. Mouse IgG served as a negative background control. Blots were probed with either **(A)** S830, **(B)** 2B7, or **(C)** D7F7 antibodies in PBST (n = 2/genotype). HTT1a was not detected in wild-type mice that do not produce the HTT1a protein. Consistent with previous findings, in zQ175 samples, all three HTT1a neoepitope antibodies successfully immunoprecipitated soluble HTT1a (▶, closed arrow) above IgG control levels. When probed with S830 a faint full-length mutant HTT (**>,** open arrow) signal was detected above the IgG control. To evaluate whether full-length HTT isoforms co-purified under these conditions, membranes were probed with either 2B7 (amino acids 7-13) or D7F7 (aa 1214-1223). A small amount of both full-length HTT isoforms (wild-type and mutant) were detected when probed with either 2B7 or D7F7 (←, arrows). IP = immunoprecipitate, IgG = immunoglobulin G, WT = wild type, HET = heterozygote, } = stacking gel, **>** = full-length HTT (wild-type or mutant), ▶ = soluble HTT1a, ID = immunodetect, ★ = nonspecific protein size marker band co-loaded in lane 1. Protein size markers are indicated in kDa.

**Supplementary Figure 5.**

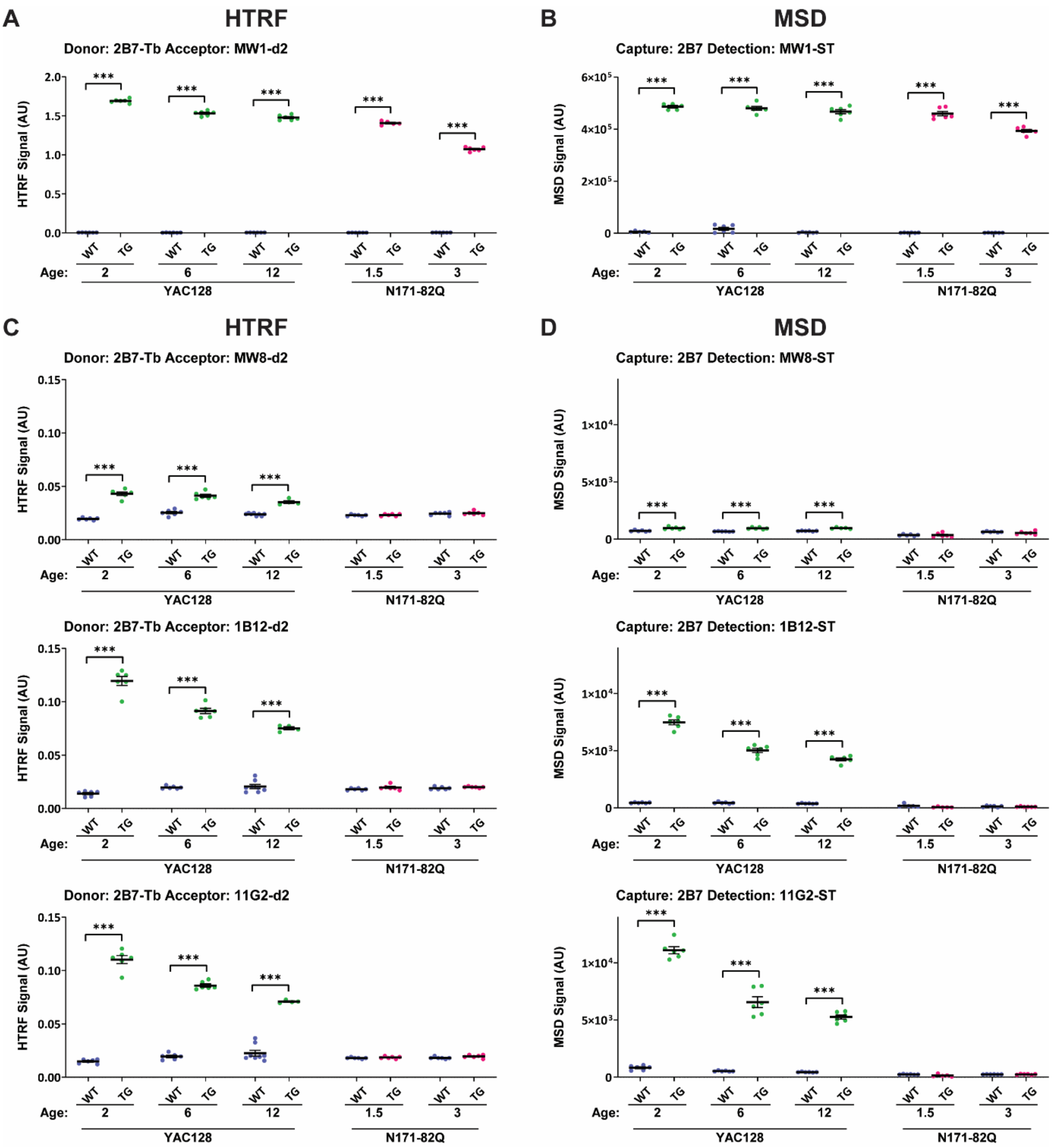

**Supplementary Figure 5. Enhancing the detection of soluble HTT1a by HTRF and MSD. (A-B)** To verify sample genotypes, the 2B7-MW1 'total mutant HTT' assay was evaluated by HTRF **(A)** and MSD **(B)** using cortical lysates from transgenic YAC128 mice at 2, 6, or 12 months of age, and from transgenic N171-82Q mice at 1.5 and 3 months, alongside age-matched wild-type littermate controls. **(C-D)** To detect soluble HTT1a, antibody pairings of 2B7-MW8, 2B7-1B12 and 2B7-11G2 were assessed by HTRF **(C)** and MSD **(D)** using the same cortical lysates (n = 3/gender/genotype/age). HTT1a was not detected in transgenic N171-82Q or wild-type mice, which do not produce the HTT1a protein. Per dataset, the *y*-axis legends were scaled the same. Data analysis was by two-way ANOVA with Bonferroni *post hoc* correction. The test statistic, degrees of freedom and *P* values for the ANOVA are reported in Supplementary Table 5. Error bars are mean ± SEM. ***$P \leq 0.001$. Tb = terbium cryptate, ST = sulfo-tag, AU = arbitrary units, WT = wild type, TG = transgenic.

**Supplementary Figure 6.**

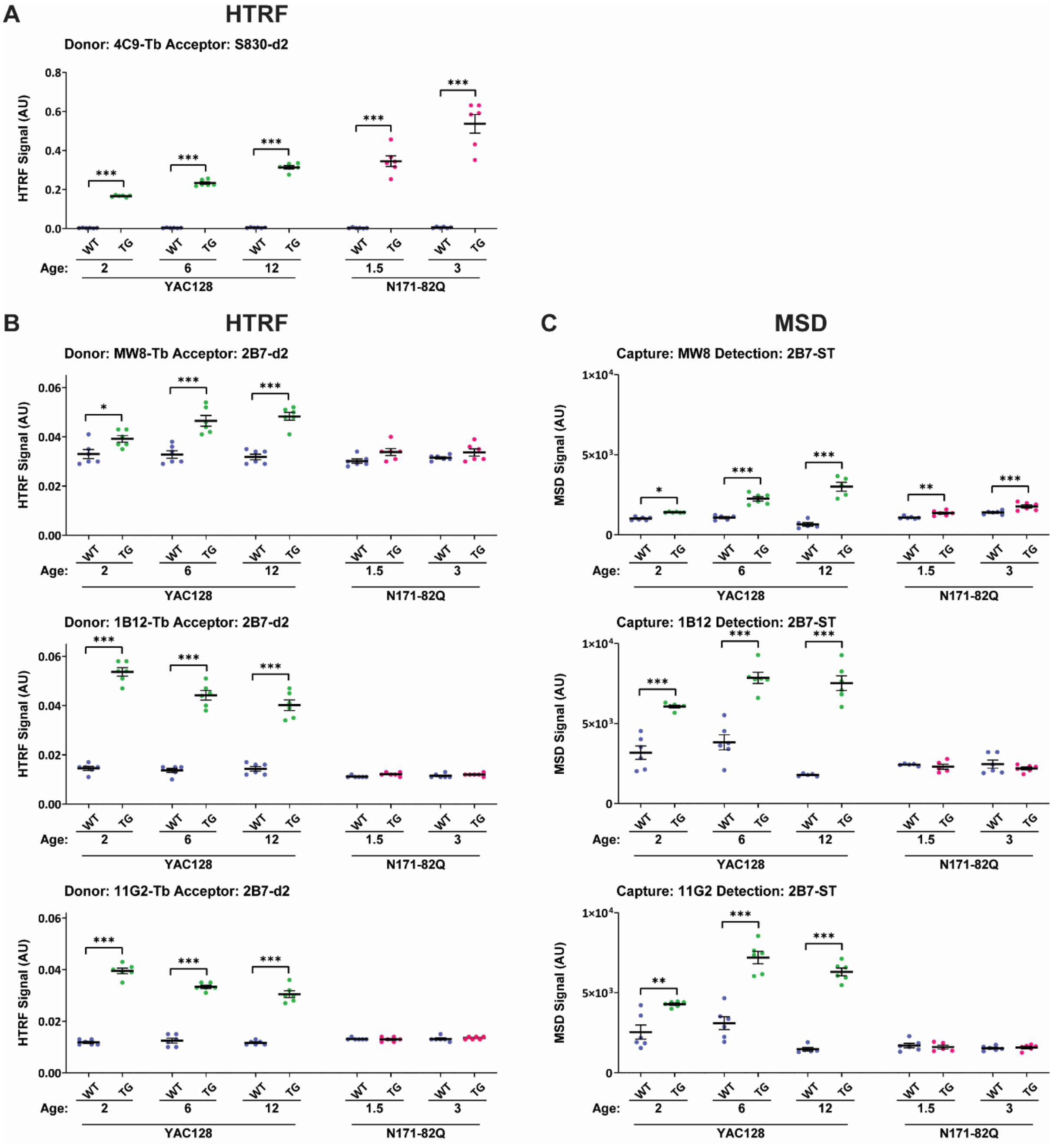

**Supplementary Figure 6. Detection of soluble or aggregated HTT1a protein isoforms with reciprocal 2B7 pairings by HTRF and MSD**. **(A)** To verify sample genotypes, the 4C9-S830 'total HTT aggregation' assay was evaluated by HTRF using cortical lysates from transgenic YAC128 mice at 2, 6, or 12 months of age, and from transgenic N171-82Q mice at 1.5 and 3 months, alongside age-matched wild-type littermate controls. **(B-C)** To detect HTT1a protein isoforms, antibody pairings of MW8-2B7, 1B12-2B7 and 11G2-2B7 were assessed by HTRF **(B)** and MSD **(C)** using the same cortical lysates (n = 3/gender/genotype/age). HTT1a was not detected in wild-type mice. Besides a small but significant signal in the MW8-2B7 MSD assay, HTT1a was not detected in transgenic N171-82Q mice, which do not produce the HTT1a protein. Per dataset, the *y*-axis legends were scaled the same. Data analysis was by two-way ANOVA with Bonferroni *post hoc* correction. The test statistic, degrees of freedom and *P* values for the ANOVA are reported in Supplementary Table 6. Error bars are mean ± SEM. $^*P \leq 0.5$, $^{**}P \leq 0.01$ and $^{***}P \leq 0.001$. Tb = terbium cryptate, ST = sulfo-tag, AU = arbitrary units, WT = wild type, TG = transgenic.

**Supplementary Figure 7.**

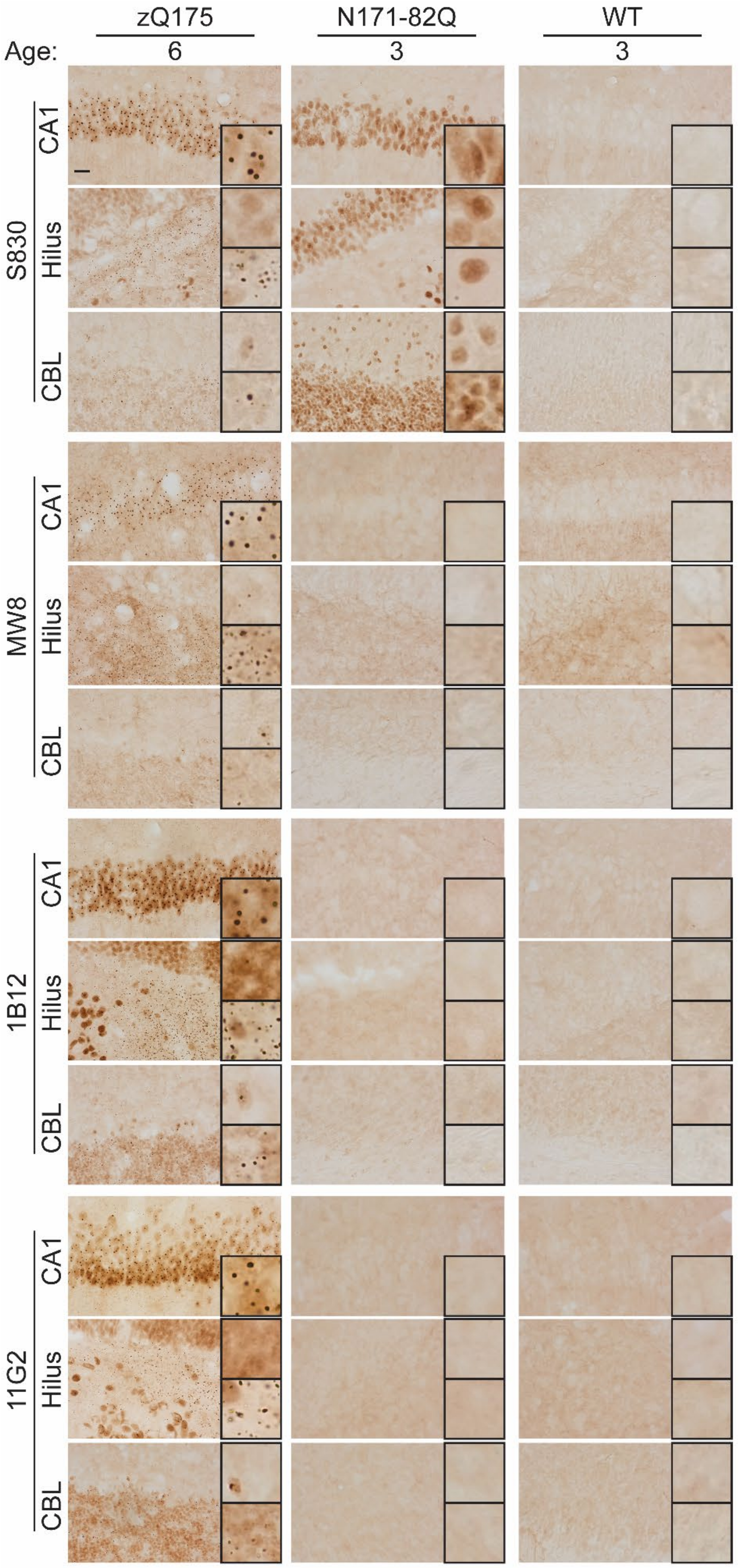

**Supplementary Figure 7. MW8, 1B12 and 11G2 are HTT1a-specific neoepitope antibodies by immunohistochemistry.** Coronal brain sections at the level of the hippocampus (CA1 and hilus regions) and cerebellum from heterozygous zQ175 mice at 6 months of age, and from transgenic N171-82Q mice at 3 months of age, alongside age-matched wild-type littermate controls were immunostained with either S830, MW8, 1B12 or 11G2 (n = 3/genotype). A HTT aggregation signal was not detected in wild-type mice. In zQ175 brain sections at 6 months, all four antibodies detected a diffuse nuclear aggregated HTT signal in addition to nuclear and cytoplasmic inclusion bodies in the hippocampus and cerebellum. In N171-82Q brain sections at 3 months, S830 detected a diffuse nuclear aggregated HTT signal in the hippocampus and cerebellum, whereas MW8, 1B12 or 11G2 did not detect any aggregated HTT, with sections appearing indistinguishable from wild-type littermate controls. These findings confirm that MW8, 1B12 and 11G2 function as neoepitope antibodies that specifically detect aggregated HTT1a. A nuclear counterstain was not applied to these sections as this would mask the nuclear signal. Scale bar = 20 μm, zoomed boxed area = 20 $\mu m^2$.WT = wild type, CA1 = hippocampal CA1 region, Hilus = hippocampal dentate gyrus, CBL = cerebellum.

### Supplementary 8.

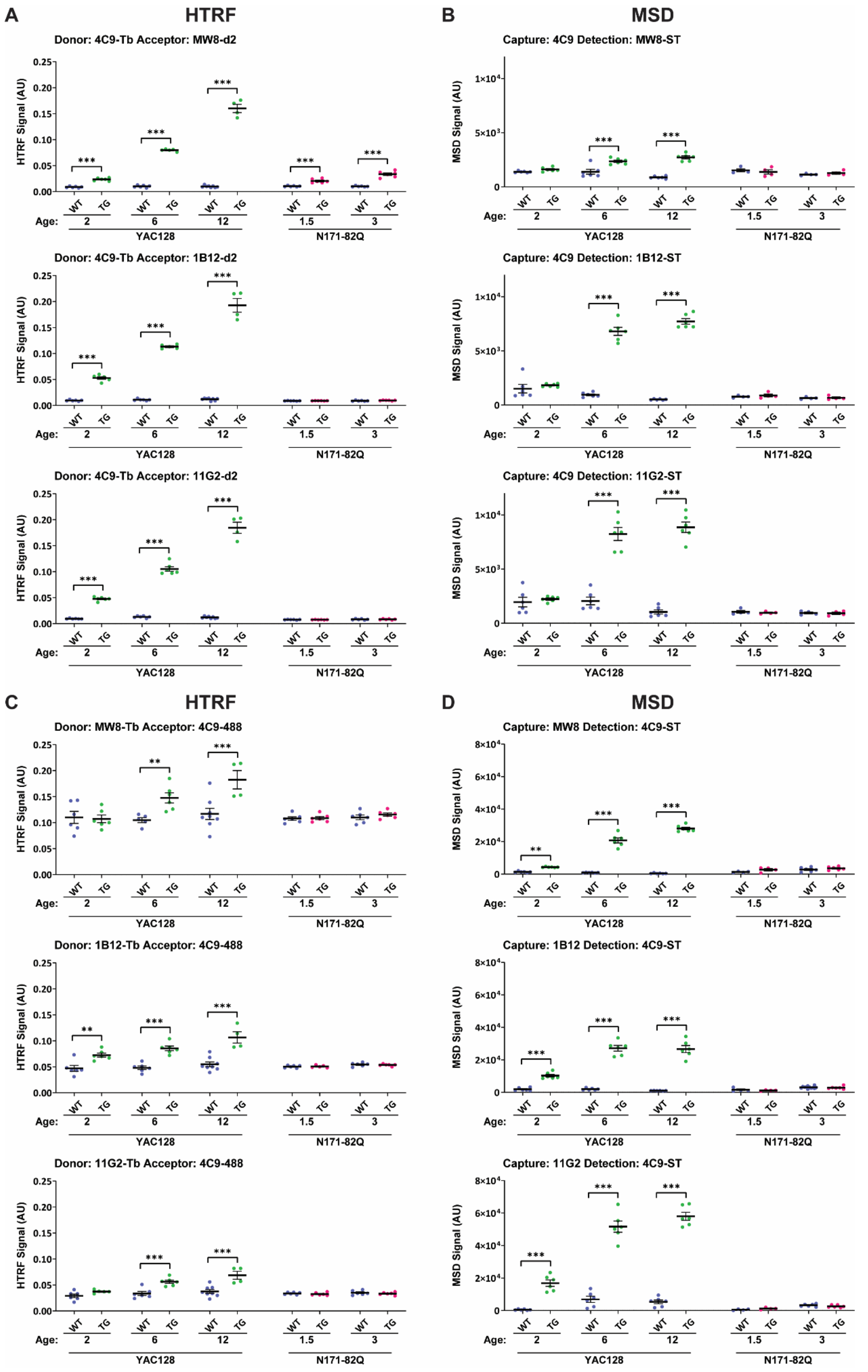

**Supplementary Figure 8. Enhancing the detection of aggregated HTT1a by HTRF and MSD. (A-D)** To detect aggregated HTT1a, reciprocal antibody pairings of 4C9-MW8, 4C9-1B12 and 4C9-11G2 were assessed by HTRF **(A)** and MSD **(B)** and of MW8-4C9, 1B12-4C9 and 11G2-4C9 were assessed by HTRF **(C)** and MSD **(D).** Assays were performed using cortical lysates from transgenic YAC128 mice at 2, 6, or 12 months of age, and from transgenic N171-82Q mice at 1.5 and 3 months, alongside age-matched wild-type littermate controls (n = 3/gender /genotype/age). Any signal detected in wild-type mice by each assay configuration represented assay background, since the endogenous mouse wild-type HTT protein does not contain the human HTT-specific sequences recognized by the 4C9 epitope. Besides a small but significant signal in the 4C9-MW8 HTRF assay, HTT1a was not detected in transgenic N171-82Q mice, which do not produce the HTT1a protein. Per dataset, the *y*-axis legends were scaled the same. Data analysis was by two-way ANOVA with Bonferroni *post hoc* correction. The test statistic, degrees of freedom and *P* values for the ANOVA are reported in Supplementary Table 7. Error bars are mean ± SEM. $^{**}P \leq 0.01$ and $^{***}P \leq 0.001$. Tb = terbium cryptate, ST = sulfo-tag, AU = arbitrary units, WT = wild type, TG = transgenic.

### Supplementary Figure 9.

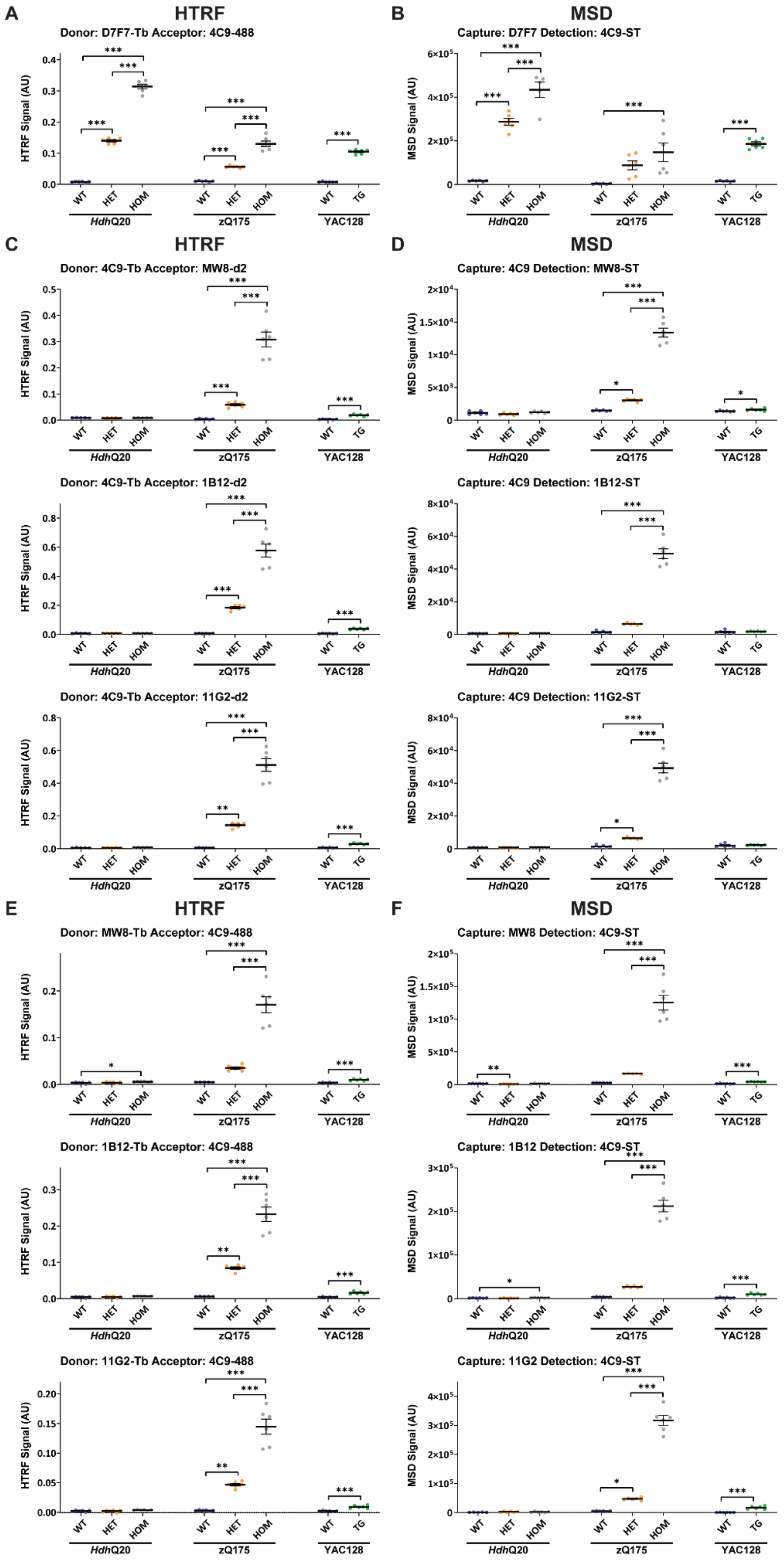

**Supplementary Figure 9. When paired with the human-specific 4C9 antibody, MW8, 1B12 and 11G2 assays do not detect these sequences when present in full-length mutant HTT. (A-B)** To verify sample genotypes and confirm that full-length mutant HTT was expressed in *Hdh*Q20, zQ175 and YAC128 mice, the D7F7-4C9 'full-length mutant HTT' assay was evaluated by HTRF **(A)** and MSD **(B)** using cortical lysates from heterozygous and homozygous *Hdh*Q20, heterozygous and homozygous zQ175 mice, and from transgenic YAC128 mice, alongside age-matched wild-type littermate controls at 2 months of age. **(C-F)** To detect aggregated HTT1a, reciprocal antibody pairings of 4C9-MW8, 4C9-1B12 and 4C9-11G2 were assessed by HTRF **(C)** and MSD **(D)** and of MW8-4C9, 1B12-4C9 and 11G2-4C9 were assessed by HTRF **(E)** and MSD **(F)** using the same cortical lysates (n = 3/gender/genotype). Any signal detected in wild-type mice by each assay configuration represented assay background, since the endogenous mouse wild-type HTT protein does not contain human HTT-specific sequences recognized by the 4C9 epitope. On both the HTRF and MSD platforms, assay signals were detected in YAC128 and zQ175 mice, and in zQ175 samples the signal intensity increased from heterozygous to homozygous animals. In contrast, a signal was not detected in heterozygous or homozygous *Hdh*Q20 mice, which appeared indistinguishable from wild-type controls. Assay specificity to exclusively detect HTT1a was confirmed, as the peptide sequences recognized by 4C9, MW8, 1B12 and 11G2 were not detected in *Hdh*Q20 mice when present within longer N-terminal proteolytic HTT fragments or full-length HTT protein isoforms. Data analysis was by one-way ANOVA with Bonferroni *post hoc* correction per mouse knock-in line, or a student's *t*-test for the YAC128. The test statistic, degrees of freedom and *P*-values for the ANOVA are reported in Supplementary Table 8. Error bars are mean ± SEM. *$P \leq 0.5$, **$P \leq 0.01$ and ***$P \leq 0.001$. Tb = terbium cryptate, ST = sulfo-tag, AU = arbitrary units, WT = wild type, HET = heterozygote, HOM = homozygote, TG = transgenic

**Supplementary Figure 10.**

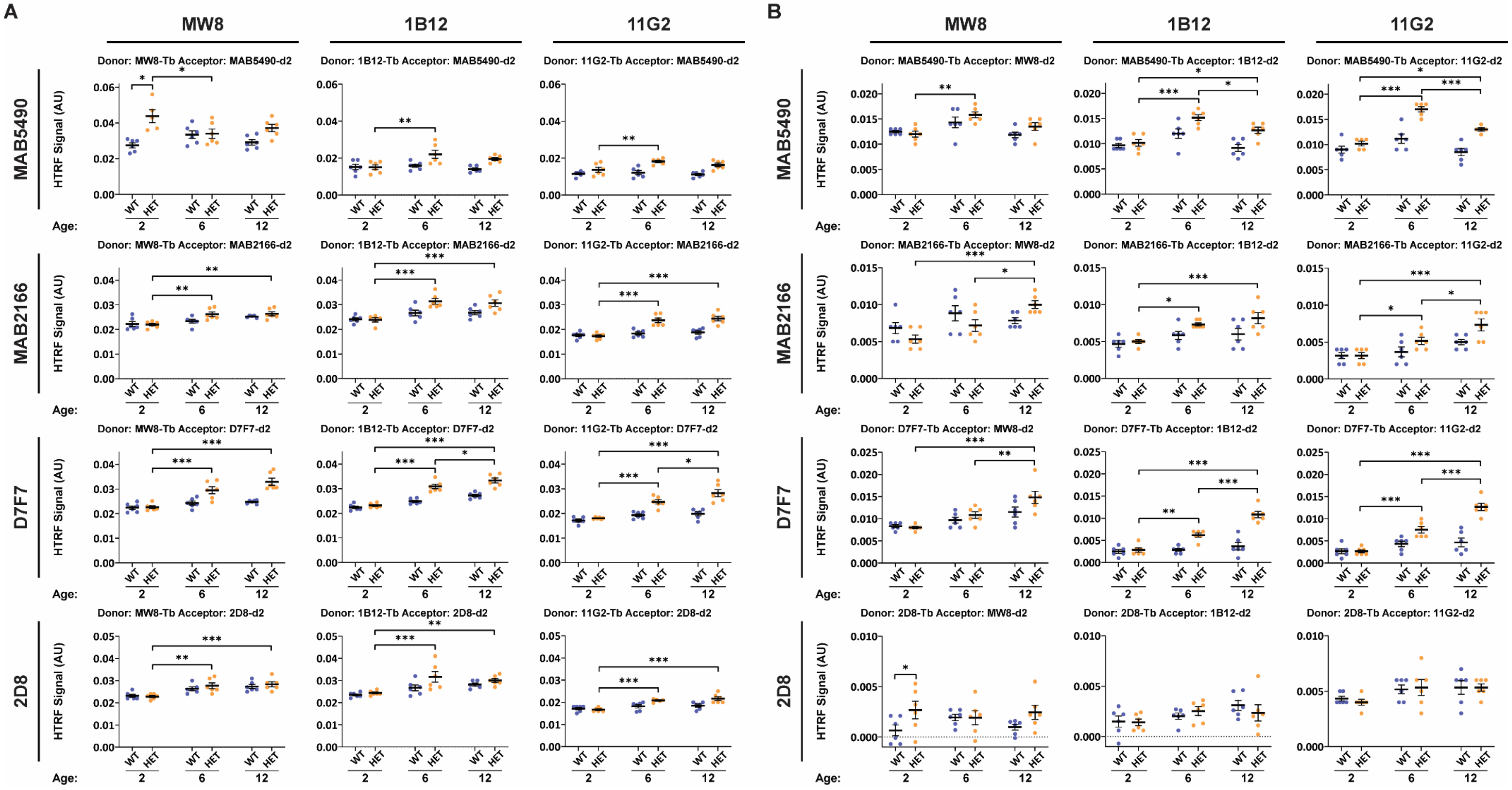

**Supplementary Figure 10. Detection of HTT fragments longer than HTT1a in HTT1a-containing aggregates by HTRF. (A-B)** Reciprocal antibody pairings of **(A)** MW8-MAB5490, 1B12-MAB5490, 11G2-MAB5490, MW8-MAB2166, 1B12-MAB2166, 11G2-MAB2166, MW8-D7F7, 1B12-D7F7, 11G2-D7F7, MW8-2D8, 1B12-2D8, 11G2-2D8 and of **(B)** MAB5490-MW8, MAB5490-1B12, MAB5490-11G2, MAB2166-MW8, MAB2166-1B12, MAB2166-11G2, D7F7-MW8, D7F7-1B12, D7F7-11G2, 2D8-MW8, 2D8-1B12, 2D8-11G2 were assessed by HTRF. Assays were performed using cortical lysates from heterozygous zQ175 mice at 2, 6, or 12 months of age, alongside age-matched wild-type littermate controls (n = 3/gender/genotype /age). Per dataset, the *y*-axis legends were scaled the same. Data analysis was by two-way ANOVA with Bonferroni *post hoc* correction. The test statistic, degrees of freedom and *P* values for the ANOVA are reported in Supplementary Table 9. Error bars are mean ± SEM. **P* ≤ 0.5, ***P* ≤ 0.01 and ****P* ≤ 0.001. Tb = terbium cryptate, AU = arbitrary units, WT = wild type, HET = heterozygote.

**Supplementary Table 4.** The test statistic, degrees of freedom and *P* values for the two-way ANOVA of data presented in **Figure 1I**.

| **Real-time quantitative PCR (qPCR)** | | |
|---|---|---|
| **Genotype x Age** | F (2, 53) = 3.3.082 | P=0.05 |
| **Genotype** | F (2, 53) = 3.074 | P=0.05 |
| **Age** | F (1, 53) = 832.8 | P<0.001 |

**Supplementary Table 5.** The test statistic, degrees of freedom and *P* values for the two-way ANOVA of data presented in **Figure 2** and **Supplementary Figure 5**.

| **HTRF** | | | | | | |
|---|---|---|---|---|---|---|
| **2B7-Tb : MW1-d2** | **zQ175** | | **YAC128** | | **N171-82Q** | |
| **Genotype x Age** | F (2, 30) = 336.2 | P<0.001 | F (2, 30) = 81.81 | P<0.001 | F (1, 19) = 531.6 | P<0.001 |
| **Genotype** | F (1, 30) = 6125 | P<0.001 | F (1, 30) = 48156 | P<0.001 | F (1, 19) = 28841 | P<0.001 |
| **Age** | F (2, 30) = 336.3 | P<0.001 | F (2, 30) = 82.07 | P<0.001 | F (1, 19) = 522.4 | P<0.001 |
| **2B7-Tb : MW8-d2** | **zQ175** | | **YAC128** | | **N171-82Q** | |
| **Genotype x Age** | F (2, 29) = 301.0 | P<0.001 | F (2, 28) = 13.30 | P<0.001 | F (1, 20) = 0.02463 | P=0.88 |
| **Genotype** | F (1, 29) = 2628 | P<0.001 | F (1, 28) = 327.5 | P<0.001 | F (1, 20) = 0.6158 | P=0.44 |
| **Age** | F (2, 29) = 318.0 | P<0.001 | F (2, 28) = 5.485 | P=0.010 | F (1, 20) = 8.892 | P=0.007 |
| **2B7-Tb : 1B12-d2** | **zQ175** | | **YAC128** | | | |
| **Genotype x Age** | F (2, 28) = 410.2 | P<0.001 | F (2, 30) = 59.46 | P<0.001 | F (1, 20) = 0.3226 | P=0.58 |
| **Genotype** | F (1, 28) = 3980 | P<0.001 | F (1, 30) = 1586 | P<0.001 | F (1, 20) = 5.161 | P=0.03 |
| **Age** | F (2, 28) = 404.7 | P<0.001 | F (2, 30) = 31.60 | P<0.001 | F (1, 20) = 1.290 | P=0.27 |
| **2B7-Tb : 11G2-d2** | **zQ175** | | **YAC128** | | **N171-82Q** | |
| **Genotype x Age** | F (2, 28) = 391.6 | P<0.001 | F (2, 30) = 50.13 | P<0.001 | F (1, 20) = 0.3543 | P=0.56 |
| **Genotype** | F (1, 28) = 3688 | P<0.001 | F (1, 30) = 1324 | P<0.001 | F (1, 20) = 3.189 | P=0.09 |
| **Age** | F (2, 28) = 403.6 | P<0.001 | F (2, 30) = 22.70 | P<0.001 | F (1, 20) = 0.9843 | P=0.33 |
| **MSD** | | | | | | |
| **Capture:2B7, Detection: MW1-ST** | **zQ175** | | **YAC128** | | **N171-82Q** | |
| **Genotype x Age** | F (2, 29) = 58.93 | P<0.001 | F (2, 28) = 1.390 | P=0.27 | F (1, 20) = 44.16 | P<0.001 |
| **Genotype** | F (1, 29) = 4396 | P<0.001 | F (1, 28) = 10245 | P<0.001 | F (1, 20) = 7280 | P<0.001 |
| **Age** | F (2, 29) = 58.62 | P<0.001 | F (2, 28) = 3.344 | P=0.05 | F (1, 20) = 44.75 | P<0.001 |
| **Capture: 2B7, Detection: MW8-ST** | **zQ175** | | **YAC128** | | **N171-82Q** | |
| **Genotype x Age** | F (2, 30) = 10.06 | P<0.001 | F (2, 29) = 0.2135 | P=0.81 | F (1, 20) = 1.028 | P=0.32 |
| **Genotype** | F (1, 30) = 63.05 | P<0.001 | F (1, 29) = 128.6 | P<0.001 | F (1, 20) = 1.045 | P=0.32 |
| **Age** | F (2, 30) = 22.98 | P<0.001 | F (2, 29) = 0.8222 | P=0.45 | F (1, 20) = 24.29 | P<0.001 |
| **Capture: 2B7, Detection: 1B12-ST** | **zQ175** | | **YAC128** | | **N171-82Q** | |
| **Genotype x Age** | F (2, 29) = 392.3 | P<0.001 | F (2, 30) = 84.89 | P<0.001 | F (1, 18) = 0.07987 | P=0.78 |
| **Genotype** | F (1, 29) = 3086 | P<0.001 | F (1, 30) = 2452 | P<0.001 | F (1, 18) = 3.782 | P=0.07 |
| **Age** | F (2, 29) = 392.5 | P<0.001 | F (2, 30) = 90.46 | P<0.001 | F (1, 18) = 0.7402 | P=0.40 |
| **Capture: 2B7, Detection: 11G2-ST** | **zQ175** | | **YAC128** | | **N171-82Q** | |
| **Genotype x Age** | F (2, 29) = 509.3 | P<0.001 | F (2, 30) = 69.08 | P<0.001 | F (1, 20) = 6.131e-005 | P>0.99 |
| **Genotype** | F (1, 29) = 3776 | P<0.001 | F (1, 30) = 1246 | P<0.001 | F (1, 20) = 0.004442 | P=0.95 |
| **Age** | F (2, 29) = 542.7 | P<0.001 | F (2, 30) = 89.45 | P<0.001 | F (1, 20) = 0.003422 | P=0.95 |

**Supplementary Table 6.** The test statistic, degrees of freedom and *P* values for the two-way ANOVA of data presented in **Figure 3** and **Supplementary Figure 6**.

| **HTRF** | | | | | | |
|---|---|---|---|---|---|---|
| **4C9-Tb : S830-d2** | **zQ175** | | **YAC128** | | **N171-82Q** | |
| **Genotype x Age** | F (2, 30) = 1021 | P<0.001 | F (2, 31) = 135.6 | P<0.001 | F (1, 20) = 11.81 | P=0.003 |
| **Genotype** | F (1, 30) = 6523 | P<0.001 | F (1, 31) = 4177 | P<0.001 | F (1, 20) = 251.3 | P<0.001 |
| **Age** | F (2, 30) = 1026 | P<0.001 | F (2, 31) = 145.7 | P<0.001 | F (1, 20) = 12.43 | P=0.002 |
| **MW8-Tb : 2B7-d2** | **zQ175** | | **YAC128** | | **N171-82Q** | |
| **Genotype x Age** | F (2, 28) = 105.7 | P<0.001 | F (2, 30) = 5.265 | P=0.01 | F (1, 20) = 0.03115 | P=0.86 |
| **Genotype** | F (1, 28) = 1250 | P<0.001 | F (1, 30) = 81.27 | P<0.001 | F (1, 20) = 6.106 | P=0.02 |
| **Age** | F (2, 28) = 106.9 | P<0.001 | F (2, 30) = 3.572 | P=0.04 | F (1, 20) = 1.526 | P=0.23 |
| **1B12-Tb : 2B7-d2** | **zQ175** | | **YAC128** | | **N171-82Q** | |
| **Genotype x Age** | F (2, 30) = 39.77 | P<0.001 | F (2, 30) = 10.27 | P<0.001 | F (1, 19) = 0.3004 | P=0.59 |
| **Genotype** | F (1, 30) = 1648 | P<0.001 | F (1, 30) = 682.0 | P<0.001 | F (1, 19) = 4.384 | P=0.05 |
| **Age** | F (2, 30) = 37.35 | P<0.001 | F (2, 30) = 11.39 | P<0.001 | F (1, 19) = 0.1137 | P=0.74 |
| **11G2-Tb : 2B7-d2** | **zQ175** | | **YAC128** | | **N171-82Q** | |
| **Genotype x Age** | F (2, 30) = 30.23 | P<0.001 | F (2, 30) = 14.63 | P<0.001 | F (1, 20) = 0.6716 | P=0.42 |
| **Genotype** | F (1, 30) = 1489 | P<0.001 | F (1, 30) = 1030 | P<0.001 | F (1, 20) = 0.07463 | P=0.79 |
| **Age** | F (2, 30) = 26.26 | P<0.001 | F (2, 30) = 14.51 | P<0.001 | F (1, 20) = 0.6716 | P=0.42 |
| **MSD** | | | | | | |
| **Capture: MW8, Detection: 2B7-ST** | **zQ175** | | **YAC128** | | **N171-82Q** | |
| **Genotype x Age** | F (2, 15) = 99.83 | P<0.001 | F (2, 27) = 29.03 | P<0.001 | F (1, 20) = 1.081 | P=0.31 |
| **Genotype** | F (1, 15) = 385.3 | P<0.001 | F (1, 27) = 154.9 | P<0.001 | F (1, 20) = 27.27 | P<0.001 |
| **Age** | F (2, 15) = 109.7 | P<0.001 | F (2, 27) = 12.66 | P<0.001 | F (1, 20) = 35.53 | P<0.001 |
| **Capture: 1B12, Detection: 2B7-ST** | **zQ175** | | **YAC128** | | **N171-82Q** | |
| **Genotype x Age** | F (2, 29) = 61.26 | P<0.001 | F (2, 29) = 7.623 | P=0.002 | F (1, 18) = 0.1897 | P=0.67 |
| **Genotype** | F (1, 29) = 1137 | P<0.001 | F (1, 29) = 202.2 | P<0.001 | F (1, 18) = 1.442 | P=0.25 |
| **Age** | F (2, 29) = 59.50 | P<0.001 | F (2, 29) = 7.444 | P=0.002 | F (1, 18) = 0.04873 | P=0.83 |
| **Capture: 11G2, Detection: 2B7-ST** | **zQ175** | | **YAC128** | | **N171-82Q** | |
| **Genotype x Age** | F (2, 28) = 188.2 | P<0.001 | F (2, 29) = 12.71 | P<0.001 | F (1, 19) = 0.4701 | P=0.50 |
| **Genotype** | F (1, 28) = 1758 | P<0.001 | F (1, 29) = 184.2 | P<0.001 | F (1, 19) = 0.04908 | P=0.83 |
| **Age** | F (2, 28) = 178.4 | P<0.001 | F (2, 29) = 16.02 | P<0.001 | F (1, 19) = 0.8495 | P=0.37 |

**Supplementary Table 7.** The test statistic, degrees of freedom and *P* values for the two-way ANOVA of data presented in **Figure 6** and **Supplementary Figure 8**.

| **HTRF** | | | | | | |
|---|---|---|---|---|---|---|
| **4C9-Tb : MW8-d2** | **zQ175** | | **YAC128** | | **N171-82Q** | |
| **Genotype x Age** | F (2, 28) = 499.4 | P<0.001 | F (2, 29) = 434.4 | P<0.001 | F (1, 20) = 20.36 | P<0.001 |
| **Genotype** | F (1, 28) = 1949 | P<0.001 | F (1, 29) = 1713 | P<0.001 | F (1, 20) = 130.2 | P<0.001 |
| **Age** | F (2, 28) = 497.3 | P<0.001 | F (2, 29) = 448.6 | P<0.001 | F (1, 20) = 18.65 | P<0.001 |
| **4C9-Tb : 1B12-d2** | | | | | | |
| **Genotype x Age** | F (2, 29) = 338.0 | P<0.001 | F (2, 29) = 161.2 | P<0.001 | F (1, 20) = 1.023 | P=0.32 |
| **Genotype** | F (1, 29) = 1893 | P<0.001 | F (1, 29) = 1191 | P<0.001 | F (1, 20) = 3.565 | P=0.07 |
| **Age** | F (2, 29) = 339.9 | P<0.001 | F (2, 29) = 172.2 | P<0.001 | F (1, 20) = 2.201 | P=0.15 |
| **4C9-Tb : 11G2-d2** | **zQ175** | | **YAC128** | | **N171-82Q** | |
| **Genotype x Age** | F (2, 29) = 590.2 | P<0.001 | F (2, 30) = 183.5 | P<0.001 | F (1, 20) = 0.009963 | P=0.92 |
| **Genotype** | F (1, 29) = 3141 | P<0.001 | F (1, 30) = 1260 | P<0.001 | F (1, 20) = 0.002491 | P=0.96 |
| **Age** | F (2, 29) = 588.8 | P<0.001 | F (2, 30) = 196.4 | P<0.001 | F (1, 20) = 3.985 | P=0.06 |
| **MW8-Tb : 4C9-d2** | **zQ175** | | **YAC128** | | **N171-82Q** | |
| **Genotype x Age** | F (2, 29) = 409.3 | P<0.001 | F (2, 29) = 5.305 | P=0.01 | F (1, 20) = 0.4685 | P=0.50 |
| **Genotype** | F (1, 29) = 1439 | P<0.001 | F (1, 29) = 15.71 | P<0.001 | F (1, 20) = 0.7627 | P=0.39 |
| **Age** | F (2, 29) = 377.7 | P<0.001 | F (2, 29) = 7.187 | P=0.003 | F (1, 20) = 1.685 | P=0.21 |
| **1B12-Tb : 4C9-d2** | **zQ175** | | **YAC128** | | **N171-82Q** | |
| **Genotype x Age** | F (2, 29) = 483.5 | P<0.001 | F (2, 30) = 2.957 | P=0.07 | F (1, 19) = 0.2500 | P=0.62 |
| **Genotype** | F (1, 29) = 2435 | P<0.001 | F (1, 30) = 76.84 | P<0.001 | F (1, 19) = 0.06635 | P=0.80 |
| **Age** | F (2, 29) = 507.0 | P<0.001 | F (2, 30) = 7.669 | P=0.002 | F (1, 19) = 9.837 | P=0.005 |
| **11G2-Tb : 4C9-d2** | **zQ175** | | **YAC128** | | **N171-82Q** | |
| **Genotype x Age** | F (2, 29) = 350.4 | P<0.001 | F (2, 29) = 3.962 | P=0.03 | F (1, 20) = 0.04043 | P=0.84 |
| **Genotype** | F (1, 29) = 1467 | P<0.001 | F (1, 29) = 39.82 | P<0.001 | F (1, 20) = 1.298 | P=0.27 |
| **Age** | F (2, 29) = 339.4 | P<0.001 | F (2, 29) = 12.13 | P<0.001 | F (1, 20) = 0.5436 | P=0.47 |
| **MSD** | | | | | | |
| **Capture: 4C9, Detection: MW8-ST** | **zQ175** | | **YAC128** | | **N171-82Q** | |
| **Genotype x Age** | F (2, 30) = 260.1 | P<0.001 | F (2, 30) = 21.94 | P<0.001 | F (1, 12) = 1.209 | P=0.29 |
| **Genotype** | F (1, 30) = 723.0 | P<0.001 | F (1, 30) = 102.3 | P<0.001 | F (1, 12) = 0.003980 | P=0.95 |
| **Age** | F (2, 30) = 256.4 | P<0.001 | F (2, 30) = 5.365 | P=0.01 | F (1, 12) = 3.620 | P=0.08 |
| **Capture: 4C9, Detection: 1B12-ST** | **zQ175** | | **YAC128** | | | |
| **Genotype x Age** | F (2, 30) = 420.0 | P<0.001 | F (2, 30) = 110.2 | P<0.001 | F (1, 12) = 0.2278 | P=0.64 |
| **Genotype** | F (1, 30) = 1133 | P<0.001 | F (1, 30) = 492.9 | P<0.001 | F (1, 12) = 0.6505 | P=0.44 |
| **Age** | F (2, 30) = 414.9 | P<0.001 | F (2, 30) = 60.24 | P<0.001 | F (1, 12) = 4.386 | P=0.06 |
| **Capture: 4C9, Detection: 11G2-ST** | **zQ175** | | **YAC128** | | **N171-82Q** | |
| **Genotype x Age** | F (2, 30) = 485.5 | P<0.001 | F (2, 30) = 49.13 | P<0.001 | F (1, 12) = 0.1015 | P=0.76 |
| **Genotype** | F (1, 30) = 1398 | P<0.001 | F (1, 30) = 212.3 | P<0.001 | F (1, 12) = 0.4305 | P=0.52 |
| **Age** | F (2, 30) = 485.2 | P<0.001 | F (2, 30) = 36.57 | P<0.001 | F (1, 12) = 0.6582 | P=0.43 |
| **Capture: MW8, Detection: 4C9-ST** | **zQ175** | | **YAC128** | | **N171-82Q** | |
| **Genotype x Age** | F (2, 28) = 364.5 | P<0.001 | F (2, 30) = 147.1 | P<0.001 | F (1, 16) = 0.3669 | P=0.55 |
| **Genotype** | F (1, 28) = 962.8 | P<0.001 | F (1, 30) = 783.4 | P<0.001 | F (1, 16) = 4.304 | P=0.05 |
| **Age** | F (2, 28) = 360.5 | P<0.001 | F (2, 30) = 127.5 | P<0.001 | F (1, 16) = 4.868 | P=0.04 |
| **Capture: 1B12, Detection: 4C9-ST** | **zQ175** | | **YAC128** | | **N171-82Q** | |
| **Genotype x Age** | F (2, 30) = 258.2 | P<0.001 | F (2, 30) = 33.86 | P<0.001 | F (1, 16) = 0.0002456 | P=0.99 |
| **Genotype** | F (1, 30) = 758.1 | P<0.001 | F (1, 30) = 408.4 | P<0.001 | F (1, 16) = 1.662 | P=0.22 |
| **Age** | F (2, 30) = 255.7 | P<0.001 | F (2, 30) = 30.93 | P<0.001 | F (1, 16) = 26.57 | P<0.001 |
| **Capture: 11G2, Detection: 4C9-ST** | **zQ175** | | **YAC128** | | **N171-82Q** | |
| **Genotype x Age** | F (2, 30) = 223.4 | P<0.001 | F (2, 30) = 40.67 | P<0.001 | F (1, 16) = 3.694 | P=0.07 |
| **Genotype** | F (1, 30) = 699.6 | P<0.001 | F (1, 30) = 480.1 | P<0.001 | F (1, 16) = 0.005324 | P=0.94 |
| **Age** | F (2, 30) = 221.9 | P<0.001 | F (2, 30) = 71.05 | P<0.001 | F (1, 16) = 38.88 | P<0.001 |

**Supplementary Table 8.** The test statistic, degrees of freedom and *P* values for the one-way ANOVA and two-tailed student's *t*-test of data presented in **Supplementary Figure 9**.

| | **One-Way ANOVA** | | | | **Student's *t*-test** | |
|---|---|---|---|---|---|---|
| **HTRF** | ***Hdh*Q20** | | **zQ175** | | **YAC128** | |
| **D7F7-Tb : 4C9-d2** | F (2, 15) = 998.6 | P<0.001 | F (2, 15) = 136.8 | P<0.001 | t=31.71 | df=10 |
| **4C9-Tb : MW8-d2** | F (2, 12) = 0.9600 | P=0.41 | F (2, 15) = 93.78 | P<0.001 | t=11.15 | df=10 |
| **4C9-Tb : 1B12-d2** | F (2, 12) = 0.0705 9 | P=0.93 | F (2, 15) = 127.7 | P<0.001 | t=14.71 | df=10 |
| **4C9-Tb : 11G2-d2** | F (2, 12) = 2.039 | P=0.17 | F (2, 15) = 130.5 | P<0.001 | t=12.52 | df=10 |
| **MW8-Tb : 4C9-d2** | F (2, 15) = 6.006 | P=0.01 | F (2, 14) = 69.04 | P<0.001 | t=7.385 | df=10 |
| **1B12-Tb : 4C9-d2** | F (2, 15) = 3.388 | P=0.06 | F (2, 14) = 88.05 | P<0.001 | t=7.777 | df=10 |
| **11G2-Tb : 4C9-d2** | F (2, 15) = 3.450 | P=0.06 | F (2, 15) = 93.34 | P<0.001 | t=6.612 | df=10 |
| **MSD** | ***Hdh*Q20** | | **zQ175** | | **YAC128** | |
| **Capture: D7F7, Detection: 4C9-ST** | F (2, 14) = 108.7 | P<0.001 | F (2, 15) = 7.221 | P=0.006 | t=18.59 | df=10 |
| **Capture: 4C9, Detection: MW8-ST** | F (2, 15) = 3.427 | P=0.06 | F (2, 15) = 265.5 | P<0.001 | t=2.647 | df=10 |
| **Capture: 4C9, Detection: 1B12-ST** | F (2, 15) = 3.088 | P=0.08 | F (2, 15) = 233.2 | P<0.001 | t=0.8073 | df=10 |
| **Capture: 4C9, Detection:11G2-ST** | F (2, 14) = 3.883 | P=0.05 | F (2, 15) = 189.0 | P<0.001 | t=0.6167 | df=10 |
| **Capture: MW8, Detection: 4C9-ST** | F (2, 15) = 7.056 | P=0.007 | F (2, 14) = 95.91 | P<0.001 | t=15.17 | df=10 |
| **Capture: 1B12, Detection: 4C9-ST** | F (2, 15) = 5.815 | P=0.01 | F (2, 15) = 226.8 | P<0.001 | t=9.310 | df=10 |
| **Capture: 11G2, Detection: 4C9-ST** | F (2, 14) = 1.727 | P=0.21 | F (2, 15) = 305.7 | P<0.001 | t=8.269 | df=10 |

**Supplementary Table 9.** The test statistic, degrees of freedom and *P* values for the two-way ANOVA of data presented in **Supplementary Figure 10**.

| **HTRF** | | |
|---|---|---|
| **MW8-Tb : MAB5490-d2** | **zQ175** | |
| **Genotype x Age** | F (2, 29) = 0.3587 | P=0.70 |
| **Genotype** | F (1, 29) = 20.51 | P<0.001 |
| **Age** | F (2, 29) = 6.391 | P=0.005 |
| **MAB5490-Tb : MW8-d2** | **zQ175** | |
| **Genotype x Age** | F (2, 30) = 1.596 | P=0.22 |
| **Genotype** | F (1, 30) = 2.602 | P=0.12 |
| **Age** | F (2, 30) = 10.27 | P<0.001 |
| **1B12-Tb : MAB5490-d2** | **zQ175** | |
| **Genotype x Age** | F (2, 30) = 3.503 | P=0.04 |
| **Genotype** | F (1, 30) = 12.75 | P=0.001 |
| **Age** | F (2, 30) = 4.274 | P=0.02 |
| **MAB5490-Tb : 1B12 -d2** | **zQ175** | |
| **Genotype x Age** | F (2, 30) = 2.998 | P=0.07 |
| **Genotype** | F (1, 30) = 18.98 | P<0.001 |
| **Age** | F (2, 30) = 15.93 | P<0.001 |
| **11G2-Tb : MAB5490-d2** | **zQ175** | |
| **Genotype x Age** | F (2, 29) = 2.373 | P=0.11 |
| **Genotype** | F (1, 29) = 34.20 | P<0.001 |
| **Age** | F (2, 29) = 3.824 | P=0.03 |
| **MAB5490-Tb :11G2 -d2** | **zQ175** | |
| **Genotype x Age** | F (2, 28) = 6.496 | P=0.005 |
| **Genotype** | F (1, 28) = 50.51 | P<0.001 |
| **Age** | F (2, 28) = 25.41 | P<0.001 |
| **MW8-Tb : MAB2166-d2** | **zQ175** | |
| **Genotype x Age** | F (2, 29) = 2.019 | P=0.15 |
| **Genotype** | F (1, 29) = 3.779 | P=0.06 |
| **Age** | F (2, 29) = 11.97 | P<0.001 |
| **MAB2166-Tb : MW8-d2** | **zQ175** | |
| **Genotype x Age** | F (2, 30) = 1.889 | P=0.17 |
| **Genotype** | F (1, 30) = 0.3383 | P=0.57 |
| **Age** | F (2, 30) = 11.36 | P<0.001 |
| **1B12-Tb : MAB2166-d2** | F (2, 29) = 3.117 | P=0.06 |
| **Genotype x Age** | F (1, 29) = 10.59 | P=0.003 |
| **Genotype** | F (2, 29) = 14.11 | P<0.001 |
| **Age** | | |
| **MAB2166-Tb : 1B12-d2** | **zQ175** | |
| **Genotype x Age** | F (2, 30) = 1.481 | P=0.24 |
| **Genotype** | F (1, 30) = 9.172 | P=0.005 |
| **Age** | F (2, 30) = 9.602 | P<0.001 |
| **11G2-Tb : MAB2166-d2** | **zQ175** | |
| **Genotype x Age** | F (2, 30) = 10.83 | P<0.001 |
| **Genotype** | F (1, 30) = 35.84 | P<0.001 |
| **Age** | F (2, 30) = 19.41 | P<0.001 |
| **MAB2166-Tb : 11G2-d2** | **zQ175** | |
| **Genotype x Age** | F (2, 30) = 2.367 | P=0.11 |
| **Genotype** | F (1, 30) = 8.292 | P=0.007 |
| **Age** | F (2, 30) = 15.38 | P<0.001 |
| **MAB2166-Tb : 11G2-d2** | **zQ175** | |
| **Genotype x Age** | F (2, 30) = 2.367 | P=0.11 |
| **Genotype** | F (1, 30) = 8.292 | P=0.007 |
| **Age** | F (2, 30) = 15.38 | P<0.001 |
| **MW8-Tb : D7F7-d2** | **zQ175** | |
| **Genotype x Age** | F (2, 30) = 8.492 | P=0.001 |
| **Genotype** | F (1, 30) = 32.25 | P<0.001 |
| **Age** | F (2, 30) = 22.01 | P<0.001 |
| **D7F7-Tb : MW8-d2** | **zQ175** | |
| **Genotype x Age** | F (2, 30) = 2.356 | P=0.11 |
| **Genotype** | F (1, 30) = 4.012 | P=0.05 |
| **Age** | F (2, 30) = 17.49 | P<0.001 |

| **1B12-Tb : D7F7-d2** | **zQ175** | |
|---|---|---|
| **Genotype x Age** | F (2, 30) = 10.88 | P<0.001 |
| **Genotype** | F (1, 30) = 64.51 | P<0.001 |
| **Age** | F (2, 30) = 68.27 | P<0.001 |
| **D7F7-Tb : 1B12-d2** | **zQ175** | |
| **Genotype x Age** | F (2, 27) = 14.82 | P<0.001 |
| **Genotype** | F (1, 27) = 58.65 | P<0.001 |
| **Age** | F (2, 27) = 28.70 | P<0.001 |
| **11G2-Tb : D7F7-d2** | **zQ175** | |
| **Genotype x Age** | F (2, 30) = 10.83 | P<0.001 |
| **Genotype** | F (1, 30) = 35.84 | P<0.001 |
| **Age** | F (2, 30) = 19.41 | P<0.001 |
| **D7F7-Tb : 11G2-d2** | **zQ175** | |
| **Genotype x Age** | F (2, 30) = 16.57 | P<0.001 |
| **Genotype** | F (1, 30) = 42.43 | P<0.001 |
| **Age** | F (2, 30) = 36.83 | P<0.001 |
| **MW8-Tb : 2D8-d2** | **zQ175** | |
| **Genotype x Age** | F (2, 30) = 0.4575 | P=0.64 |
| **Genotype** | F (1, 30) = 0.7843 | P=0.38 |
| **Age** | F (2, 30) = 15.71 | P<0.001 |
| **2D8-Tb : MW8-d2** | **zQ175** | |
| **Genotype x Age** | F (2, 30) = 1.492 | P=0.24 |
| **Genotype** | F (1, 30) = 5.405 | P=0.03 |
| **Age** | F (2, 30) = 0.09682 | P=0.91 |
| **1B12-Tb : 2D8-d2** | **zQ175** | |
| **Genotype x Age** | F (2, 30) = 1.564 | P=0.23 |
| **Genotype** | F (1, 30) = 6.475 | P=0.02 |
| **Age** | F (2, 30) = 11.96 | P<0.001 |
| **2D8-Tb : 1B12-d2** | **zQ175** | |
| **Genotype x Age** | F (2, 30) = 0.7175 | P=0.50 |
| **Genotype** | F (1, 30) = 0.06300 | P=0.80 |
| **Age** | F (2, 30) = 3.152 | P=0.06 |
| **11G2-Tb: 2D8-d2** | **zQ175** | |
| **Genotype x Age** | F (2, 30) = 10.83 | P<0.001 |
| **Genotype** | F (1, 30) = 35.84 | P<0.001 |
| **Age** | F (2, 30) = 19.41 | P<0.001 |
| **2D8-Tb : 11G2-d2** | **zQ175** | |
| **Genotype x Age** | F (2, 30) = 0.1528 | P=0.86 |
| **Genotype** | F (1, 30) = 0.02183 | P=0.88 |
| **Age** | F (2, 30) = 3.996 | P=0.03 |

**Supplementary Table 10.** The test statistic, degrees of freedom and *P* values for the two-way ANOVA of data presented in **Figure 7**.

| MSD | | |
|---|---|---|
| **Capture: MW8, Detection: MAB5490-ST** | **zQ175** | |
| **Genotype x Age** | F (2, 28) = 30.64 | P<0.001 |
| **Genotype** | F (1, 28) = 107.2 | P<0.001 |
| **Age** | F (2, 28) = 12.45 | P<0.001 |
| **Capture: MAB5490, Detection: MW8-ST** | **zQ175** | |
| **Genotype x Age** | F (2, 27) = 10.35 | P<0.001 |
| **Genotype** | F (1, 27) = 30.26 | P<0.001 |
| **Age** | F (2, 27) = 1.431 | P=0.26 |
| **Capture: 1B12, Detection: MAB5490-ST** | **zQ175** | |
| **Genotype x Age** | F (2, 30) = 102.7 | P<0.001 |
| **Genotype** | F (1, 30) = 247.3 | P<0.001 |
| **Age** | F (2, 30) = 51.72 | P<0.001 |
| **Capture: MAB5490, Detection: 1B12-ST** | **zQ175** | |
| **Genotype x Age** | F (2, 27) = 60.62 | P<0.001 |
| **Genotype** | F (1, 27) = 230.0 | P<0.001 |
| **Age** | F (2, 27) = 27.35 | P<0.001 |
| **Capture: 11G2, Detection: MAB5490-ST** | **zQ175** | |
| **Genotype x Age** | F (2, 30) = 106.6 | P<0.001 |
| **Genotype** | F (1, 30) = 333.5 | P<0.001 |
| **Age** | F (2, 30) = 31.50 | P<0.001 |
| **Capture: MAB5490, Detection: 11G2-ST** | **zQ175** | |
| **Genotype x Age** | F (2, 27) = 51.23 | P<0.001 |
| **Genotype** | F (1, 27) = 189.7 | P<0.001 |
| **Age** | F (2, 27) = 8.336 | P=0.002 |
| **Capture: MW8, Detection: MAB2166-ST** | **zQ175** | |
| **Genotype x Age** | F (2, 30) = 196.2 | P<0.001 |
| **Genotype** | F (1, 30) = 733.2 | P<0.001 |
| **Age** | F (2, 30) = 102.1 | P<0.001 |
| **Capture: MAB2166, Detection: MW8-ST** | **zQ175** | |
| **Genotype x Age** | F (2, 24) = 9.808 | P<0.001 |
| **Genotype** | F (1, 24) = 13.89 | P=0.001 |
| **Age** | F (2, 24) = 22.27 | P<0.001 |
| **Capture: 1B12, Detection: MAB2166-ST** | **zQ175** | |
| **Genotype x Age** | F (2, 30) = 234.5 | P<0.001 |
| **Genotype** | F (1, 30) = 566.2 | P<0.001 |
| **Age** | F (2, 30) = 180.2 | P<0.001 |
| **Capture: MAB2166, Detection: 1B12-ST** | **zQ175** | |
| **Genotype x Age** | F (2, 29) = 10.92 | P<0.001 |
| **Genotype** | F (1, 29) = 42.81 | P<0.001 |
| **Age** | F (2, 29) = 26.74 | P<0.001 |
| **Capture: 11G2, Detection: MAB2166-ST** | **zQ175** | |
| **Genotype x Age** | F (2, 30) = 126.1 | P<0.001 |
| **Genotype** | F (1, 30) = 455.4 | P<0.001 |
| **Age** | F (2, 30) = 150.9 | P<0.001 |
| **Capture: MAB2166, Detection: 11G2-ST** | **zQ175** | |
| **Genotype x Age** | F (2, 29) = 26.71 | P<0.001 |
| **Genotype** | F (1, 29) = 107.7 | P<0.001 |
| **Age** | F (2, 29) = 28.97 | P<0.001 |
| **Capture: MW8, Detection: D7F7-ST** | **zQ175** | |
| **Genotype x Age** | F (2, 26) = 48.15 | P<0.001 |
| **Genotype** | F (1, 26) = 137.4 | P<0.001 |
| **Age** | F (2, 26) = 37.85 | P<0.001 |
| **Capture: D7F7, Detection: MW8-ST** | **zQ175** | |
| **Genotype x Age** | F (2, 29) = 68.46 | P<0.001 |
| **Genotype** | F (1, 29) = 144.0 | P<0.001 |
| **Age** | F (2, 29) = 45.76 | P<0.001 |
| **Capture: 1B12, Detection: D7F7-ST** | **zQ175** | |
| **Genotype x Age** | F (2, 28) = 244.0 | P<0.001 |
| **Genotype** | F (1, 28) = 699.8 | P<0.001 |
| **Age** | F (2, 28) = 210.8 | P<0.001 |

| | | |
|---|---|---|
| **Capture: D7F7, Detection: 1B12-ST** | **zQ175** | |
| **Genotype x Age** | F (2, 28) = 70.23 | P<0.001 |
| **Genotype** | F (1, 28) = 415.9 | P<0.001 |
| **Age** | F (2, 28) = 72.42 | P<0.001 |
| **Capture: 11G2, Detection: D7F7-ST** | **zQ175** | |
| **Genotype x Age** | F (2, 29) = 214.1 | P<0.001 |
| **Genotype** | F (1, 29) = 700.4 | P<0.001 |
| **Age** | F (2, 29) = 194.2 | P<0.001 |
| **Capture: D7F7, Detection: 11G2-ST** | **zQ175** | |
| **Genotype x Age** | F (2, 30) = 269.6 | P<0.001 |
| **Genotype** | F (1, 30) = 1037 | P<0.001 |
| **Age** | F (2, 30) = 297.2 | P<0.001 |
| **Capture: MW8, Detection: 2D8-ST** | **zQ175** | |
| **Genotype x Age** | F (2, 28) = 6.136 | P=0.006 |
| **Genotype** | F (1, 28) = 44.57 | P<0.001 |
| **Age** | F (2, 28) = 9.236 | P<0.001 |
| **Capture: 2D8, Detection: MW8-ST** | **zQ175** | |
| **Genotype x Age** | F (2, 28) = 1.263 | P=0.30 |
| **Genotype** | F (1, 28) = 2.707 | P=0.11 |
| **Age** | F (2, 28) = 4.875 | P=0.02 |
| **Capture: 1B12, Detection: 2D8-ST** | **zQ175** | |
| **Genotype x Age** | F (2, 29) = 4.843 | P=0.02 |
| **Genotype** | F (1, 29) = 81.71 | P<0.001 |
| **Age** | F (2, 29) = 3.469 | P=0.04 |
| **Capture: 2D8, Detection: 1B12-ST** | **zQ175** | |
| **Genotype x Age** | F (2, 28) = 18.91 | P<0.001 |
| **Genotype** | F (1, 28) = 294.8 | P<0.001 |
| **Age** | F (2, 28) = 32.51 | P<0.001 |
| **Capture: 11G2, Detection: 2D8-ST** | **zQ175** | |
| **Genotype x Age** | F (2, 29) = 25.95 | P<0.001 |
| **Genotype** | F (1, 29) = 228.7 | P<0.001 |
| **Age** | F (2, 29) = 12.85 | P<0.001 |
| **Capture: 2D8, Detection: 11G2-ST** | **zQ175** | |
| **Genotype x Age** | F (2, 30) = 122.7 | P<0.001 |
| **Genotype** | F (1, 30) = 397.9 | P<0.001 |
| **Age** | F (2, 30) = 154.4 | P<0.001 |

**Supplementary Table 11.** The test statistic, degrees of freedom and *P* values for the one-way ANOVA and two-tailed student's *t*-test of data presented in **Figure 8**.

| | **One-Way ANOVA** | | | | | | | | | | | |
|---|---|---|---|---|---|---|---|---|---|---|---|---|
| **HTRF** | ***Hdh*Q20 11W** | | ***Hdh*Q50 11W** | | ***Hdh*Q80 11W** | | ***Hdh*Q111 11W** | | **CAG140 11W** | | **zQ175 11W** | |
| **2B7-Tb : 1B12-d2** | F (2, 15) =0.4648 | P=0.64 | F (2, 15) = 0.1364 | P=0.87 | F (2, 15) = 7.922 | P=0.004 | F (2, 15) = 4.719 | P=0.03 | F (2, 15) = 3.667 | P=0.05 | F (2, 14) = 3.556 | P=0.006 |
| **MSD** | ***Hdh*Q2011W** | | ***Hdh*Q50 11W** | | ***Hdh*Q80 11W** | | ***Hdh*Q111 11W** | | **CAG140 11W** | | **zQ175 11W** | |
| **Capture: 2B7, Detection: MW1-ST** | F (2, 14) = 1.993 | P=0.17 | F (2, 15) = 5.217 | P=0.02 | F (2, 15) = 4.552 | P=0.03 | F (2, 15) = 1.996 | P=0.17 | F (2, 15) = 2.938 | P=0.08 | F (2, 14) = 7.241 | P=0.007 |

| | **Student's *t*-test** | | | |
|---|---|---|---|---|
| **HTRF** | **YAC128 9W** | | **N171-82Q 7W** | |
| **4C9-Tb : 1B12-d2** | t=198.2 | df=10 | t=23.55 | df=10 |
| **MSD** | **YAC128 9W** | | **N171-82Q 7W** | |
| **Capture : 11G2, Detection: 4C9-ST** | t=21.91 | df=9 | t=0.9533 | df=10 |

| | **One-Way ANOVA** | | | | | | | | | | | |
|---|---|---|---|---|---|---|---|---|---|---|---|---|
| **HTRF** | ***Hdh*Q20 11W** | | ***Hdh*Q50 11W** | | ***Hdh*Q80 11W** | | ***Hdh*Q111 11W** | | **CAG140 11W** | | **zQ175 11W** | |
| **4C9-Tb : 1B12-d2** | F (2, 15) =5.374 | P=0.01 | F (2, 15) = 28.11 | P<0.001 | F (2, 15) = 301.2 | P=0.2 | F (2, 15) = 297.6 | P<0.001 | F (2, 15) = 568.8 | P=<0.001 | F (2, 14) = 631.2 | P<0.001 |
| **MSD** | ***Hdh*Q20 11W** | | ***Hdh*Q50 11W** | | ***Hdh*Q80 11W** | | ***Hdh*Q111 11W** | | **CAG140 11W** | | **zQ175 11W** | |
| **Capture: 11G2, Detection: 4C9-ST** | F (2, 15) = 4.312 | P=0.03 | F (2, 15) = 30.45 | P<0.001 | F (2, 15) = 218.7 | P<0.001 | F (2, 15) = 99.70 | P<0.001 | F (2, 15) = 142.5 | P<0.001 | F (2, 14) = 322.5 | P<0.001 |

| | **Student's *t*-test** | | | | | | | |
|---|---|---|---|---|---|---|---|---|
| **HTRF** | **zQ175 26W** | | **YAC128 9W** | | **YAC128 26W** | | **N171-82Q 14W** | |
| **4C9-Tb : 1B12-d2** | t=33.71 | df=10 | t=17.66 | df=10 | t=22.33 | df=10 | t=0.3612 | df=6 |
| **MSD** | **zQ175 26W** | | **YAC128 9W** | | **YAC128 26W** | | **N171-82Q 14W** | |
| **Capture: 11G2, Detection: 4C9-ST** | t=14.89 | df=10 | t=9.947 | df=10 | t=18.51 | df=10 | t=0.9012 | df=6 |
